# Targeting oncogenic KRasG13C with nucleotide-based covalent inhibitors

**DOI:** 10.1101/2022.07.25.501348

**Authors:** Lisa Goebel, Tonia Kirschner, Sandra Koska, Amrita Rai, Petra Janning, Stefano Maffini, Helge Vatheuer, Paul Czodrowski, Roger S. Goody, Matthias P. Müller, Daniel Rauh

**Affiliations:** Department of Chemistry and Chemical Biology, TU Dortmund University, Dortmund, Germany; Department of Structural Biochemistry, Max Planck Institute of Molecular Physiology, Dortmund, Germany; Department of Chemical Biology, Max Planck Institute of Molecular Physiology, Dortmund, Germany; Department of Mechanistic Cell Biology, Max Planck Institute of Molecular Physiology, Dortmund, Germany

**Keywords:** cancer, Ras, G13C, nucleotide analogues, covalent inhibitors

## Abstract

Mutations within Ras proteins represent major drivers in human cancer. In this study, we report the structure-based design, synthesis, as well as biochemical and cellular evaluation of nucleotide-based covalent inhibitors for KRasG13C, an important oncogenic mutant of Ras that has not been successfully addressed in the past. Mass spectrometry experiments and kinetic studies reveal promising molecular properties of these covalent inhibitors, and X-ray crystallographic analysis has yielded the first reported crystal structures of KRasG13C covalently locked with these GDP analogues. Importantly, KRasG13C covalently modified with these inhibitors can no longer undergo SOS-catalysed nucleotide exchange. As a final proof-of-concept, we show that in contrast to KRasG13C, the covalently locked protein is unable to induce oncogenic signalling in cells, further highlighting the possibility of using nucleotide-based inhibitors with covalent warheads in KRasG13C-driven cancer.

## Introduction

Ras proteins act as key regulators of many cellular processes by switching between inactive GDP-bound and active GTP-bound states, the latter specifically activating several downstream signalling pathways.^1^ Oncogenic Ras mutations that lead to dysregulation of the switch mechanism are found in about 25% of all human cancers, including three of the most lethal forms (lung, colon, and pancreatic cancer). Among the Ras proteins, KRas is the predominantly mutated isoform (85%), followed by NRas (11%) and HRas (4%), with mutational hotspots at amino acid positions G12, G13, and Q61.^1,2^ Because of their prominent role in cancer, Ras oncogenes were identified as attractive targets for cancer therapy since their initial discovery in 1981, but attempts to target Ras have been largely unsuccessful and Ras proteins were long considered undruggable. After decades of failure, new interest has recently arisen from selective targeting of the G12C oncogenic mutant of KRas.^3-11^ Inhibitors that bind irreversibly to the G12C mutated cysteine residue within a previously unknown switch-II pocket were originally identified and designed in the Shokat laboratory, and have been further developed within the academic and industrial world to advance candidates into the clinic.^12-16^ In May 2021 the first-in-class KRasG12C inhibitor Sotorasib (Amgen) was approved by the FDA for the treatment of non-small-cell lung cancer (NSCLC), confirming the therapeutic susceptibility of mutant KRas in cancer.^17^ A second approach for targeting mutant KRas has been described using nucleotide competitive inhibitors that can covalently bind to KRasG12C. ^18,19^ Strategies involving direct competition with nucleotide binding were originally set aside because of the high affinity of GDP/GTP for Ras and high cellular GDP/GTP concentrations. However, the combination of nucleotide competition with covalent binding of the inhibitors to the Ras protein has fuelled new hope. Gray and colleagues developed SML-8-73-1, a GDP derivative harbouring an electrophilic group on the β-phosphate to irreversibly bind to the mutant cysteine at position 12 within the P-loop of Ras.^18^ Unfortunately, the modification of the β-phosphate leads to a dramatic loss of affinity because of the loss of important interactions with the protein and the Mg^2+^ ion.^20^ In this publication, we demonstrate that GDP/GTP/GppCp analogues with an electrophilic group attached to the ribose interact with the necessary high reversible affinity to KRas and are able to react covalently with KRasG13C.

## Results and discussion

### Selective covalent modification of KRasG13C by 2’,3’-modified nucleotide analogues

Available structural information about the binding of GDP and GTP towards Ras provides a detailed understanding of the underlying high affinity of nucleotides through multiple reversible interactions. Based on published crystal structures of Ras proteins (PDB 4obe and 4nmm) and a multiple sequence alignment of the Ras small GTPase superfamily (Figure S1/S2), we designed and synthesized guanine nucleotide-based inhibitors with an additional Michael acceptor as a covalent warhead for targeting oncogenic KRas variants harboring cysteines in the P-loop (KRasG12C and KRasG13C). Whereas the G12C mutation has been successfully addressed in the past, KRasG13C is a largely unexplored target in cancer therapy. However, based on pKa calculations, we were able to show that the G13C mutation should also be generally addressable by appropriately positioned Michael acceptors (Table S1). In contrast to Gray and colleagues, we chose the 2’,3’-OH groups of the ribose for attachment of the warhead since modifications at this position do not significantly alter nucleotide affinity (Figure 1A/B).^21^ Nucleotide derivatives with different linkers (eda: ethylenediamine, pda: propylenediamine, bda: butylenediamine) were synthesised based on published procedures (Scheme S1), including GDP/GTP/GppCp analogues (R = OH), resulting in the formation of mixed 2’ and 3’-isomers, as well as dGTP analogues (R = H).^21,22^ In addition to acrylamide-bearing nucleotides that could potentially bind irreversibly to cysteine containing P-loop mutants via Michael addition, we prepared acetamide derivatives as non-reactive control analogues (Figure 1B). The cysteine light-version of KRas constructs lacking other cysteines (C51S, C80L, C118S) were used for initial MS experiments, which indicated that the acrylamide nucleotide derivatives can selectively react with KRasG13C_1-169_ (Cys-light), but not with KRasG12C_1-169_ (Cys-light) (Figure 1C/D/E). The eda linker led to the most efficient covalent protein modification (Figure 1C). This tendency is presumably because of a favourable orientation of the reactive warhead and/or a reduced flexibility. Using dGTP analogues, the rate of covalent protein modification was further increased in the case of the eda derivative, indicating that the linker in the 3’-position of the ribose is superior to the 2’-position with respect to targeting KRasG13C. On incubating KRasG13C_1-169_ (Cys-light) with the GTP analogues, we observed at intermediate stages a mixture of covalently bound GDP and GTP forms of KRasG13C_1-169_ (Cys-light), but ultimately the reaction yielded only the diphosphate form, indicating that the nucleotides were still hydrolysed after the covalent reaction and were properly positioned in the active site of KRas (Figure S7). The time-resolved labelling of KRasG13C_1-169_ (Cys-light) with either GDP or GTP derivatives led to comparable covalent protein modification rates (Figure S8). Upon incubating KRasG13C_1-169_ (Cys-light) with a non-cleavable GppCp derivative, we also observed covalent modification, but without subsequent hydrolysis of the nucleotide (Figure S9). To further investigate the specificity of the reaction towards KRasG13C_1-169_ (Cys-light), we also tested the wild type protein. For KRasWT_1-169_, very little unspecific labelling was observed at pH 9.5 compared to the G13C mutant (Figure 1E, Figure S10), thus showing that the warhead reacts preferentially with the cysteine at position 13, but not other cysteines in KRas nor the additional cysteine in KRasG12C. In addition, the multiple sequence alignment of the Ras small GTPase superfamily revealed that only about 7% of the GTPase members contain cysteines within the P-loop that might potentially be accessible by our linker design and only 3 contain Cys at the position equivalent to residue 13 in Ras (Arl4a, RheBL1, Rab21). However, since other cysteines that are located within the P-loop or other regions close to the nucleotide are mostly further remote compared to Cys13, suitable design and optimization of the linker will most likely allow sufficient specificity towards KRasG13C also *in vivo* (Figure S1/S2), similar to the observed preference of the cysteine in position 13 compared to position 12.

**Figure 1.**
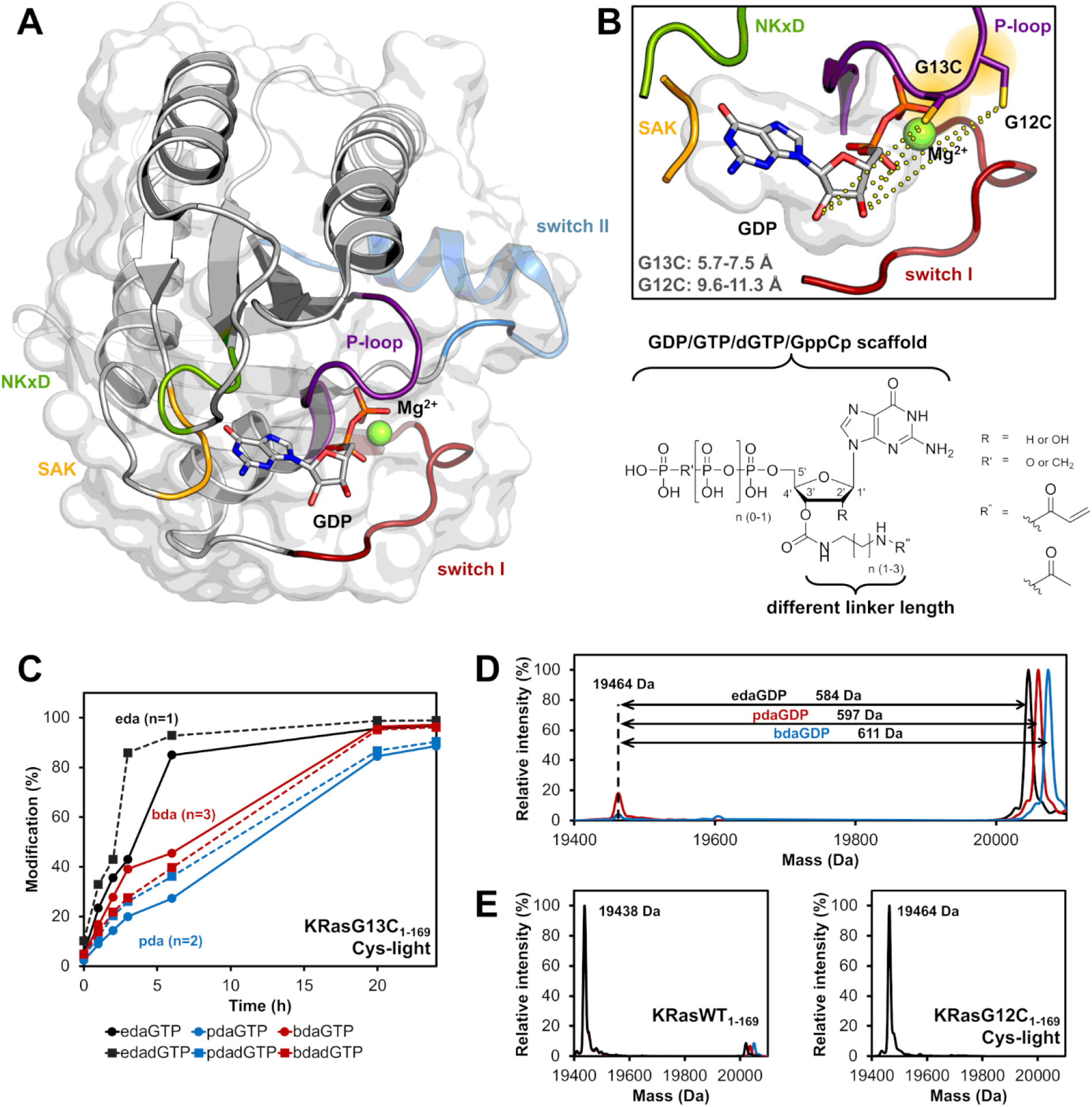
Rational design of nucleotide-based covalent KRasG13C inhibitors. (**A**) Structure of KRasWT in the GDP-bound state (grey, P-loop: violet, switch I: red, switch II: blue, NKxD: green, SAK: yellow; PDB 4obe). (**B**) Model of KRasG13C (model is based on PDB 4obe) showing the distances between the cysteine residue and the OH-groups of the ribose and the general structure of nucleotide derivatives bearing an electrophilic group at the 2’,3’-position of the ribose moiety, showing that position 12 is further remote compared to position 13 and thus explaining the observed specificity of covalent bond formation. (**C**) Time dependent analysis of the covalent modification of KRasG13C_1-169_ (Cys-light) at pH 9.5 using different linker lengths (eda: ethylenediamine, pda: propylenediamine and bda: butylenediamine). (**D**) Covalent modification of KRasG13C_1-169_ (Cys-light) proteins at pH 9.5 after 24 h at room temperature. (**E**) In contrast, KRasG12C_1-169_ (Cys-light) mutant was not modified by nucleotide derivatives and for KRasWT_1-169_ very little unspecific labelling at pH 9.5 was observed compared to the G13C mutant.

### Reversible affinities of the nucleotide analogues are comparable to those of the unmodified nucleotides

To evaluate the impact of the attached linker on nucleotide binding, we first determined the affinity and kinetics of the interaction of the nucleotide derivatives with Ras compared to unmodified GDP/GTP that have dissociation constants (K_D_) in the picomolar range. For this purpose, we measured the kinetics of the nucleotide association (k_on_) in a stopped-flow instrument using competition experiments between mantdGDP (2 μM) and increasing amounts of competing nucleotides (1, 2, and 6 μM).^20^ As shown in Figure S11, the competitive binding of the competing nucleotide and mantdGDP led to a significant decrease in the fluorescence signal because of smaller amounts of mantdGDP binding to KRas. By fitting the data to a previously described model^20^ (Figure S11B), we obtained the corresponding k_on_ values, and those for the nucleotide analogues were comparable to those of the unmodified nucleotide (GDP; Table S2). To further analyse the ability of the modified nucleotides to compete with GDP/GTP, KRasWT:GDP was mixed with equal amounts of the acetamide derivatives and incubated either for 7 days at room temperature in the absence of EDTA, for 24 h at 4 °C in the presence of EDTA, or for 1 h at room temperature in the presence of SOS (guanine nucleotide exchange factor) to increase the rate of nucleotide exchange and to allow the reaction to equilibrate. After this equilibration time and buffer exchange, the Ras proteins were concentrated and the nucleotide state was analysed by isocratic HPLC runs. By integrating the corresponding peaks for GDP and the guanosine nucleotide analogues after distinct time points, we observed that the modified nucleotides can indeed compete with GDP (Figure S12, Table S3). Based on the relative abundance of bound nucleotides determined by the HPLC assay, the dissociation constants for the pda and bda derivatives were calculated to be 8.6±1.3 pM and 9.6±0.5 pM, respectively (overlap with the GDP elution peak prevented accurate determination in the case of the eda-derivative) (SI, Results and Discussion, Table 1). Thus, the attached linker has very little impact on the reversible interaction and the affinity, an important fact that must be considered for the approach of using nucleotide-competitive inhibitors.^20^ In contrast, SML-8-73-1, a GDP derivative harbouring an electrophilic group on the β-phosphate showed a dramatic loss of reversible affinity (K_D_ = ∼140 nM).^23^

**Table 1.** Overview of the calculated kinetic parameters (K_D_, k_on_ and k_off_). K_D_ values of pdaGDP and bdaGDP obtained from an HPLC-based approach and show the average from the 3 different experiments in the absence or the presence of EDTA or SOS, respectively (see also Supplementary Table S3 for the individual results). k_on_ rate constants obtained from competitive binding experiments with mantdGDP. For reference, the K_D_ values of GDP and the nucleotide analogue SML-8-73-1 are also listed.^20,23^

| | $K_D$<br>[pM] | $k_{on}$<br>[$\mu M^{-1}s^{-1}$] | $k_{off}$<br>[$s^{-1}$] |
| --- | --- | --- | --- |
| **GDP** | 2.5 | 4.22 | $1.1 \times 10^{-5}$ |
| **pdaGDP** | $8.6 \pm 1.3$ | 3.34 | $2.9 \times 10^{-5}$ |
| **bdaGDP** | $9.6 \pm 0.5$ | 3.12 | $3.0 \times 10^{-5}$ |
| **SML-8-73-1** | ~140 nM | - | - |

### First crystal structures of the KRasG13C mutant

To gain further insight into the binding mode of the covalently bound nucleotides, we solved the first X-ray crystal structure of the oncogenic KRasG13C mutant, in this case with covalently bound edaGDP (PDB 7ok3) and bdaGDP (PDB 7ok4) (Figure 2, Table S4). The overall structures are very similar to the known structure of KRas:GDP (PDB 4obe), and the nucleotide scaffold, as well as the covalent linkage for both nucleotide analogues with cysteine at position 13 are well resolved in the electron density (Figure 2C/E, Figure S13). Interestingly, only the 3’-isomer of the nucleotide analogues was observed in the structures, consistent with results of the MS experiments comparing the efficiency of labelling of the dGTP derivative and the mixed isomers of the GTP derivative and showing a faster reaction for dGTP (Figure 1C). Both structures showed that the nucleotides are bound within the active site in a manner that is comparable to non-covalently bound GDP in other structures of Ras, with similar reversible interactions between the protein and the nucleotide and the additional well-resolved covalent link to Cys13. However, both structures lacked the Mg^2+^ ions in the active site despite a Mg^2+^ concentration of 2 mM in the Ras solution. The missing Mg^2+^-ions are probably a result of the crystallization buffer containing NH_4_F or NaF, leading to precipitation of poorly soluble MgF_2_, and this has also been observed in other PDB-deposited structures presumably because of similar effects of the reservoir solutions used in the crystallization process (e.g., PDB 4m1o, 4lyf, 4lyh, 4m21, 4m1s, 4m1t, 4m1t, 4m1y).^3^ In both crystal structures, the eda and bda linker are remote from the Mg^2+^ binding site and do not directly interfere with Mg^2+^ binding, suggesting that Mg^2+^ can generally bind. Thus, the structural analysis verified the nucleotide binding pose and the high reversible affinity comparable to the natural nucleotides and will guide design and optimisation of the linker and the warhead in further studies.

**Figure 2.**
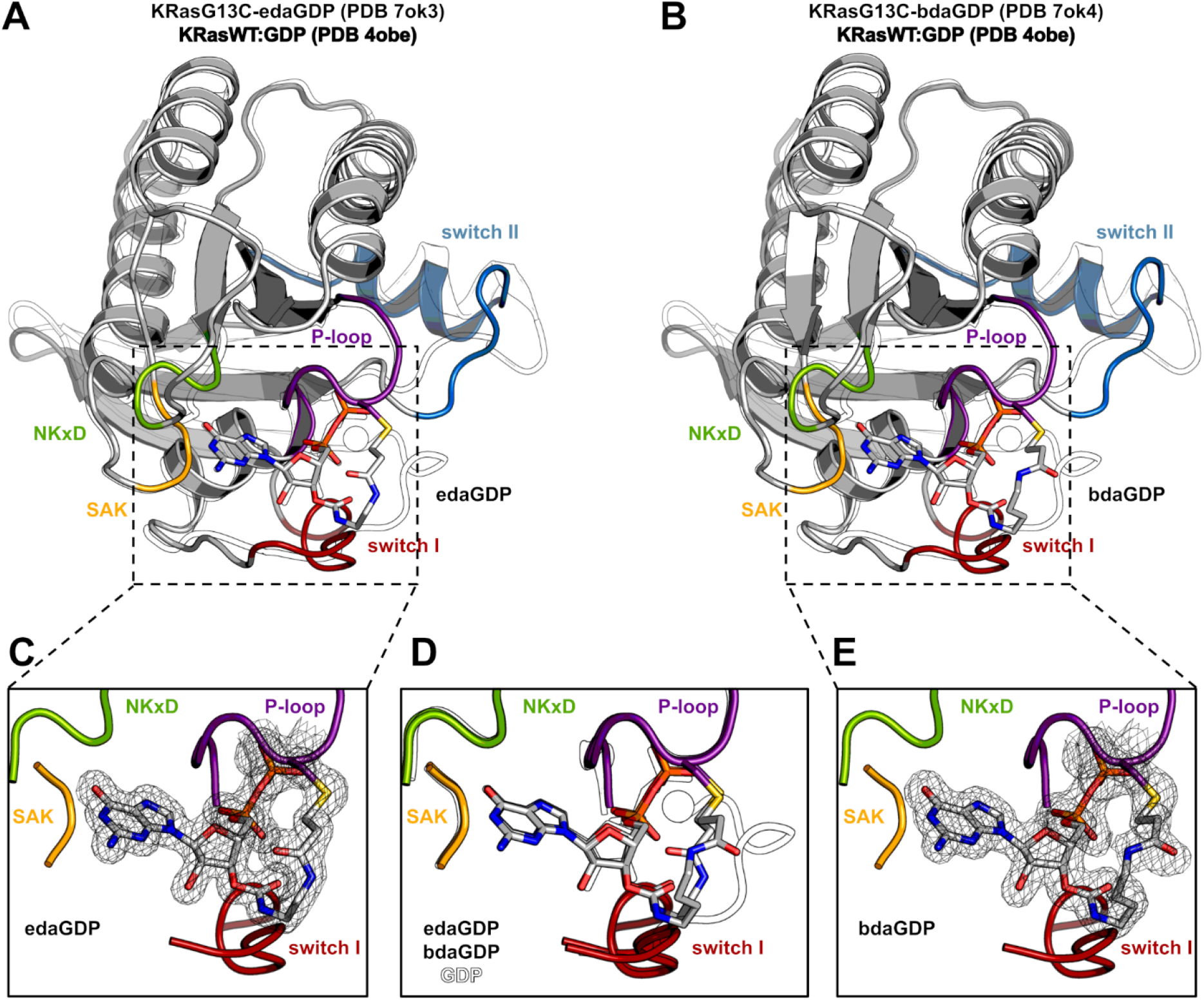
Crystal structure of KRasG13C covalently locked with either edaGDP or bdaGDP. (**A**) Comparison of KRasG13C covalently locked with edaGDP (grey, P-loop: violet, switch I: red, switch II: blue, NKxD: green, SAK: yellow, PDB 7ok3) and KRas:GDP (white, PDB 4obe). (**B**) Comparison of KRasG13C covalently locked with bdaGDP (grey, P-loop: violet, switch I: red, switch II: blue, NKxD: green, SAK: yellow, PDB 7ok4) and KRas:GDP (white, PDB 4obe). (**C**) Enlarged view of the nucleotide binding pocket in KRasG13C-edaGDP showing the 2Fo-Fc electron density map (countered at 1.0 σ). (**D**) Structural superposition of KRas:GDP and the covalently locked G13C mutants. (**E**) Enlarged view of the nucleotide binding pocket in KRasG13C-bdaGDP showing the 2Fo-Fc electron density map (countered at 1.0 σ).

### Inhibition of oncogenic KRasG13C signalling by covalent nucleotide analogues

Finally, we set out to test whether the GDP nucleotide derivatives we are focusing on for cancer therapy were indeed able to inhibit oncogenic signalling by KRasG13C. In a first experiment, we tested and compared the SOS-catalysed nucleotide exchange on Ras. Whereas SOS efficiently catalysed nucleotide exchange on KRasWT, KRasG13C and as a control on KRasG13C:acetyledaGDP, it was unable to do so with the covalently locked G13C-edaGDP mutant, showing that the protein is indeed locked in the inactive state (Figure 3, Figure S14A). Interestingly, in these experiments we also observed a drastically increased intrinsic nucleotide exchange rate for KRasG13C compared to KRasWT in the absence of SOS (Figure 3A, blue area), an effect that probably contributes to the increased signalling and oncogenic effect of this mutant, and also this effect is abrogated in the case of covalently locked KRasG13C-edaGDP.^24^ Similarly, no SOS-catalysed nucleotide exchange was observed in case of covalently modified KRasG13C-edaGppCp, indicating that the protein can also be trapped in the active conformation with the corresponding nucleoside-triphosphates (Figure S14B). Thus, the modified GppCp derivatives could also be used as artificial and irreversible activators and generally as tool compounds to further investigate the biological role of KRasG13C in cells. In addition to SOS-catalysed nucleotide exchange, GAP-stimulated GTP hydrolysis was also analysed. While the intrinsic GTP hydrolysis of KRasG13C-edaGTP (t_1/2_ = 187 min) is comparable to that of KRasWT:GTP (t_1/2_ = 126 min) and even faster than for the KRasG12C mutant (t_1/2_ = 300 min)^25^ (Figure S15), a drastically decreased GAP-stimulated GTP hydrolysis was observed for the G13C mutant as expected.^26^

**Figure 3.**
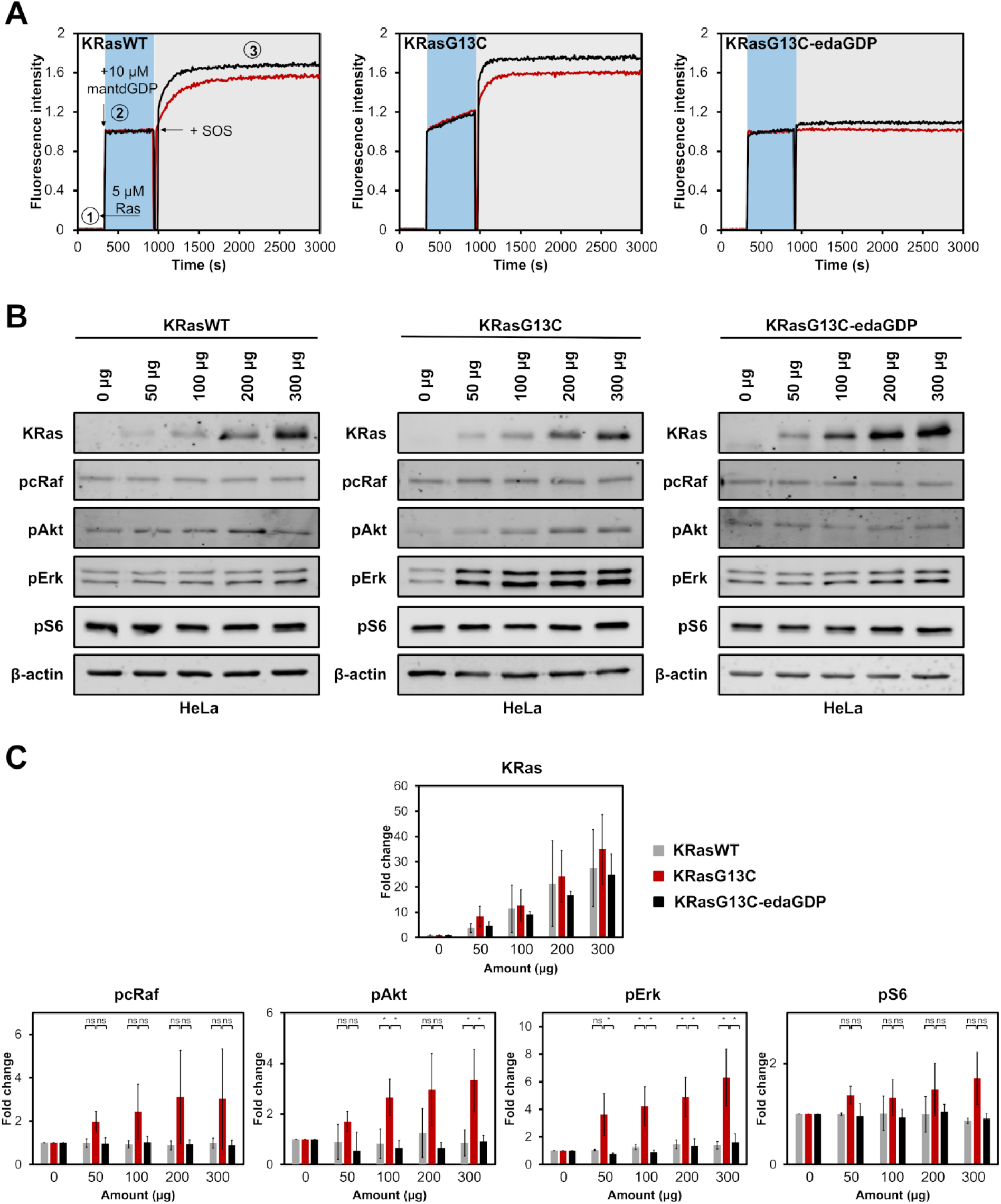
Cellular evaluation of nucleotide-based covalent inhibitors. (**A**) GEF-catalysed nucleotide exchange of KRasWT:GDP, KRasG13C:GDP and KRasG13C-edaGDP (step 1) which were mixed with an excess of mantdGDP (step 2) and subsequently with 0.25 µM (red curve) or 0.5 µM (black curve) of SOS (step 3). The intrinsic nucleotide exchange is depicted in the blue box whereas the SOS-catalysed nucleotide exchange is shown in the grey box. (**B**) Western blot analysis after electroporation of indicated amounts of full-length KRasWT, KRasG13C and KRasG13C-edaGDP into HeLa cells. (**C**) Quantification of protein levels from western blots was performed using Empiria Studio (Li-Cor). The mean fold change was plotted against increasing amounts of protein. Error bars indicate the standard deviation for each measurement (n = 3). One-way ANOVA was performed using GraphPad Prism.

After a detailed *in vitro* characterization of the nucleotide analogues, the next experiment was to investigate whether oncogenic signalling could also be inhibited *in vivo*. Since the nucleotides are unable to cross the cell membrane without loss of the phosphate groups and full labelling of the G13C mutant was only achieved at relatively high pH values, we used electroporation to deliver recombinant full-length KRasWT, KRasG13C and KRasG13C-edaGDP into human cells. Importantly, we confirmed that the covalently locked protein can still be fully farnesylated *in vitro* (Figure S16) and the covalent modification of full-length KRasG13C was further validated through MS/MS-analysis, which revealed selective labelling of the targeted cysteine residue at position 13 (Figure S17). Upon electroporation into HeLa cells, a concentration-dependent increase in abundance of KRas was observed for all variants, indicating successful delivery into cells (Figure 3, Figure S18-S19). However, whereas we observed a concentration-dependent activation of the downstream signalling and upregulation of pcRaf, pAkt, pErk and pS6 upon delivery of KRasG13C, the covalently locked variant was unable to induce these effects and, similarly to KRasWT, no increase in downstream signalling was observed (Figure 3, Figure S18-S19). In addition, upon electroporation of non-covalently modified KRasG13C:acetyledaGDP or covalently modified KRasG13C-edaGppCP, activation of the Ras pathway comparable to KRasG13C was observed, showing that covalent modification is essential for inhibition of oncogenic signalling and that artificial activation can be induced using non-hydrolyzable GTP derivatives, which potentially adds further possibilities of using these nucleotide analogues to study Ras biology (Figure S20-S21). Thus, our cellular data provide an additional proof-of-concept for the use of nucleotide-based covalent inhibitors and activators in KRasG13C-driven cancer.

## Conclusion

In summary, we have successfully developed nucleotide-based covalent inhibitors of oncogenic KRasG13C, a variant of KRas that is largely unexplored as a target even though it is a frequently observed mutation in cancer.^27^ Thorough biochemical and structural characterization revealed that the nucleotide analogues designed and synthesized in this publication have similar affinities towards Ras compared to their natural counterparts. Since we are currently unable to directly test the nucleotides in cells, we instead extensively tested them *in vitro* regarding their ability to covalently lock KRasG13C in the inactive state and to effectively inhibit oncogenic signalling. We could show that the nucleotides prohibit (SOS-mediated) nucleotide exchange and lead to an effective interference in the induction of oncogenic effects by KRasG13C in living cells. Thus, after the first successful examples of inhibition of KRasG12C by Shokat and colleagues, our study breaks ground to effectively inhibit another important oncogenic variant of Ras by using small molecules and further optimisation regarding the reactivity of the nucleotides using a structure-guided approach to optimize the orientation towards the relevant cysteine and the reactivity of the warhead as well as the delivery into cells by protecting esterification are currently underway.^28,29^ Additionally, it is possible that appropriate modification of the linker length and the electrophilic warhead would also make KRasG12C or other relevant mutants (e.g., HRasG12S in Costello syndrome)^30^ a possible target in a similar approach.

## Methods

### Protein expression and purification

KRasWT_1-169_ and KRasG13C_1-169_ Cys-light (C51S C80L C118S) were expressed in BL21 (DE3) *E. coli* whereas the full-length KRasWT and KRasG13C were expressed in BL21 (DE3) RIL *E. coli* at 37 °C. Protein expression was induced at A600 nm of 0.5 by addition of 0.2-0.3 mM isopropyl-b-D-thiogalactoside (IPTG), and growth was continued at 19 °C overnight. The bacteria were collected by centrifugation and the obtained pellet resuspended in Ni-NTA buffer (KRas_1-169_: 50 mM Tris pH 8.0, 250 mM NaCl, 40 mM imidazole, 4 mM MgCl_2_, 10 µM GDP, 1 mM dithiothreitol (DTT), and 5% glycerol; full-length KRas: 50 mM HEPES pH 7.2, 500 mM LiCl, 2 mM MgCl_2_, 10 µM GDP, 2 mM β-mercaptoethanol (βME)). The cells were lysed with a microfluidizer, and after addition of protease inhibitor cocktail (Roche complete EDTA free) and 1% CHAPS (w/v) stirring was continued for 1 h at 4 °C. The lysate was cleared by centrifugation (35,000 x g, 1 h) and the supernatant was loaded onto a Ni-affinity chromatography column (Qiagen Ni-NTA Superflow, 20 mL) pre-equilibrated with Ni-NTA buffer. KRas_1-169_ proteins were eluted with a linear gradient of imidazole buffer (40 mM – 500 mM), whereas the full-length KRas proteins were collected with a stepwise elution (2, 5, 10, 20, 30, 50 and 100% of Ni-NTA-buffer containing 500 mM imidazole). For cleavage of the N-terminal hexahistidine-tag, TEV protease was added to the pooled elution fractions and dialyzed overnight into dialysis buffer at 4 °C (KRas_1-169_: 25 mM Tris pH 8.0, 100 mM NaCl, 4 mM MgCl_2_, 10 µM GDP, 1 mM DTT, and 5% glycerol; full-length KRas: 20 mM HEPES pH 7.2, 200 mM NaCl, 2 mM MgCl_2_, 10 µM GDP, 2 mM βME and 5% glycerol). The cleaved protein was then applied to a reverse Ni-affinity chromatography column and finally purified by size-exclusion chromatography (GE HiLoad 16/60 Superdex 75 pg) in a final buffer containing 20 mM HEPES pH 7.5, 100 mM NaCl, 2 mM MgCl_2_, 10 µM GDP, 1 mM TCEP, and 5% glycerol.

### Synthesis of nucleotide-based covalent inhibitors

Synthesis of the nucleotide analogues was carried out according to the protocol established by Cremo et al. (1990) and described by Eberth et al. (2005).^21,22^ A strong cation exchanger (Ion exchanger I, Merck, Darmstadt, Germany) was used to prepare the tributylammonium salts of the nucleotides. The commercially available sodium salt of the nucleotide (0.5 mmol) was dissolved in 3 mL ddH_2_O and was applied on the column pre-equilibrated with pyridine/H_2_O (1:1). Nucleotide elution was achieved using methanol/H_2_O (1:1) and the eluate was dripped into 1 mL TBA. After monitoring the nucleotide elution by spotting samples onto a TLC plate with fluorescent indicator and removing of the methanol/H_2_O solution the mixture was dried by repeated rotary evaporation from dry DMF (3 × 20 mL). The remaining solid was dissolved in 20 mL dry DMF and CDI (2.5 mmol) was added under argon atmosphere. After stirring overnight at 4 °C to form the carbonate the reaction was quenched by the addition of absolute methanol (150 µL). The primary amine (eda, pda or bda; 2.5 mmol) dissolved in 5 mL dry DMF was slowly added to the carbonate mixture to prepare the phosphoramidate derivative and the resulting precipitate was recovered by centrifugation (10000 rpm, 10 min) and washed three times with DMF. The solid was dissolved in 20 mL ddH_2_O and the pH was adjusted to 1.5 with 0.25 M hydrochloric acid to hydrolyse the phosphoramidate. After stirring at 4 °C for 1-3 d the mixture was then raised to pH 7.5 using 0.25 M NaOH. The nucleotides were purified at 4 °C on a Q-Sepharose column (column volume: 130 mL) preequilibrated with 50 mM triethylammonium bicarbonate buffer (pH 7.6) and eluted by a linear gradient of 50 mM – 1 M TEAB over 600 min with a flow rate of 1 mL/min. The nucleotide containing fractions were analysed by HPLC 50 mM KPi pH 6.6, 10 mM TBAB, 16% ACN; column: ProntoSIL® 120-5-C18-AQ, Bischoff, Germany) and were lyophilized several times from ddH_2_O to remove the buffer. The nucleotide based covalent inhibitors were prepared by dissolving eda, pda or bda nucleotides in a small amount of tetraborate buffer (100 mM, pH 8.5) and adding N-acryloxysuccinimide (1 eq.) dissolved in 25 µL DMSO. The reaction mixture was stirred for 24 h at room temperature and the reaction progress was monitored using HPLC. The covalent nucleotide analogues were purified using Q-Sepharose as described above and were stored at -20 °C as concentrated solutions (∼ 100 mM) in 200 mM HEPES (pH 7.5).

### Nucleotide exchange

50 µM Ras protein was incubated with a 10-fold excess of nucleotides in 20 mM HEPES (pH 7.5), 100 mM NaCl, 2 mM MgCl_2_, 1 mM TCEP, 5% glycerol, and 10 mM EDTA for 3 h at 4 °C. Nucleotide exchange was terminated by the addition of 20 mM MgCl_2_ and Ras proteins were washed using centrifugal filter devices to remove any unbound nucleotide. Nucleotide exchange was controlled by isocratic HPLC runs.

### Covalent modification of proteins

To analyze the amount of covalent modification of KRasG13C_1-169_, 50 µM Ras protein was incubated with a 10-fold excess of the acryl-bearing nucleotides in 100 mM CHES (pH 9.5), 50 mM NaCl, 1 mM TCEP, and 1 mM EDTA. After incubation at room temperature for the appropriate time, the modification of the protein was controlled by ESI-MS. For covalent modification of full-length KRasG13C, a nucleotide exchange with a 10-fold excess of the acryl-bearing nucleotides at pH 7.5 was first performed following incubation at room temperature for 24 h at pH 9.5 for covalent protein modification. Covalent protein modification was controlled by ESI-MS. The MS spectra were recorded on a VelosPro IonTrap (Thermo Scientific) with an EC 50/3 Nucleodur C18 1.8 μm column (Macherey and Nagel) and a gradient of the mobile phase A (0.1 % formic acid in water) to B (0.1 % formic acid in acetonitrile).

### Stopped-flow experiments

The association kinetics of the nucleotide analogues were analyzed with a SX-20 stopped-flow instrument (Applied Photophysics) at 25 °C in a buffer consisting of 25 mM HEPES (pH 7.5), 100 mM NaCl, 1 mM MgCl_2_, and 0.5 mM TCEP. 1 µM of nucleotide-free KRas^20^ in one syringe was mixed rapidly with 2 µM mantdGDP in the other. In subsequent experiments, the second syringe contained competing nucleotides (1, 2, and 6 µM) in addition to 2 µM mantdGDP. The resulting progress curves were globally fit using KinTek Explorer to obtain the corresponding association rate constants as previously described (Fig. S11, Tab. S2).^31^

### HPLC-based approach for determination of affinities relative to GDP

The relative affinities of the nucleotide analogues were measured using an HPLC-based approach. 50 µM of KRasWT_1-169_:GDP was mixed with 50 µM of the acetamide derivatives and incubated either for 7 days at room temperature in the absence of EDTA, for 24 h at 4 °C in the presence of 10 mM EDTA, or for 1 h at room temperature in the presence of SOS in a buffer consisting of 20 mM HEPES (pH 7.5), 100 mM NaCl, 2 mM MgCl_2_, 1 mM TCEP and 5% glycerol. After incubation, the Ras proteins were washed 5 times with buffer (15 mL) using centrifugal filter devices to remove any unbound nucleotide, concentrated and analyzed by isocratic HPLC runs. The resulting curves were analyzed by integrating the corresponding peaks for GDP and the guanosine nucleotide analogues after distinct time points using the Agilent ChemStation Software (Fig. S12, Tab. S3).

### Crystallization and structure determination

KRasG13C_1-169_ Cys-light (C51S C80L C118S) was covalently modified by incubating 100 µM KRas with a 10-fold excess of the acryl-bearing nucleotide in 100 mM CHES (pH 9.5), 50 mM NaCl, 1 mM TCEP, and 1 mM EDTA at room temperature for 24 h. Modification of the protein was monitored by ESI-MS and terminated by the addition of 20 mM MgCl_2_. The protein was purified by size-exclusion chromatography (GE HiLoad 16/60 Superdex 75 pg) in a final buffer containing 20 mM HEPES pH 7.5, 100 mM NaCl, 2 mM MgCl_2_, 1 mM TCEP, and 5% glycerol, and subsequently concentrated to 67 mg/mL. To identify the initial crystallization conditions, commercially available protein crystallization screens (JCSG Core I –IV Suites, PEGs and PACT) were used. Using a TTP labtech Mosquito LCP crystal liquid-handling robot, 100 nL of protein solution was mixed with 100 nL reservoir solution in 96-well plates, and crystals were grown using the hanging-drop method at 20 °C. After 1 day of incubation, one successful crystallization condition was obtained for KRasG13C-edaGDP (0.2 M (NH_4_)F, 20% PEG3350) and KRasG13C-bdaGDP (0.2 M NaF, 20% PEG3350), and the crystals were cryoprotected in mother liquor and flash cooled in liquid nitrogen. The data sets were collected at the PXII X10SA beamline of the Swiss Light Source (Paul Scherrer Institute, Villigen, Switzerland) and indexed and scaled using XDS.^32^ The crystal structures were solved by molecular replacement with PHASER using PDB 4obe as a template.^33^ The manual modification of the molecule of the asymmetric unit was performed using the program COOT,^34^ and with the help of the Dundee PRODRG server,^35^ the inhibitor topology file was generated. For multiple cycles of refinement, PHENIX.refine^36^ was employed, the final structure was evaluated by the PDB_REDOserver,^37^ and crystal structures were visualized using PyMOL. Data collection, structure refinement statistics, and further details for data collection are provided in Table S4.

### Guanine nucleotide-exchange factor assay

SOS-catalysed nucleotide exchange was monitored at 25 °C in a FluoroMax-3 spectrofluorometer (excitation at 360 nm, emission at 440 nm) in 20 mM HEPES (pH 7.5), 100 mM NaCl, 2 mM MgCl_2_, and 1 mM TCEP. 5 µM KRas_1-169_ was mixed with 10 µM mantdGDP and subsequently with different concentrations of SOS (0.25 µM and 0.5 µM) (Fig. 3A; Fig. S14).

### Cell Culture

HeLa cells were obtained from the American Type Culture Collection (ATCC) and were cultured in DMEM medium (Gibco) supplemented with 10% fetal bovine serum (FBS) (PAN-Biotech) and 1% penicillin/streptomycin (Gibco). Cells were cultured in a humidified incubator at 37 °C in the presence of 5% CO_2_.

### Electroporation

Electroporation of full-length KRas constructs (KRasWT, KRasG13C, KRasG13C-edaGDP, KRasG13C:acetyledaGDP and KRasG13C-edaGppCp) was performed based on the protocol described by Alex et al. (2019) using the Neon Transfection System Kit (Thermo Fisher).^38^ For electroporation, 3 million cells per experiment were harvested by trypsinization, washed with PBS, and resuspended in 85 µL of the electroporation buffer R (Thermo Fisher). Increasing amounts of recombinant protein samples for each construct (0, 50, 100, 200 and 300 µg) were diluted 1:1 in buffer R followed by the addition of 30 µL of this protein master mix to the cell suspension. This cellular slurry was loaded into a 100 mL Neon Pipette Tip (Thermo Fisher) and electroporated with 2×35 msec pulses at 1000V. After electroporation, the cells were washed twice with PBS (15 mL) to remove non-internalized extracellular protein and the cell pellet was resuspended in 2 mL complete growth media. Cells were transferred into six-well tissue culture plates (Sarstedt) and incubated for 24 h at 37 °C and 5% CO_2_ in a humidified incubator for recovery before being processed for western blotting analysis.

### Western Blot analysis

After recovery of the electroporation, cells were washed twice with ice-cold PBS and lysed in 100 µL of phosphatase and protease inhibitor containing RIPA buffer (Cell Signaling Technology). Cells were incubated on ice for 30 min and then harvested by scraping followed by centrifugation at 14,000 rpm for 10 min at 4 °C. Protein concentrations were determined using the Pierce BCA protein assay (Thermo) following the manufacturer’s recommended procedure. Equal amounts of protein (10 µg) were analyzed by SDS-PAGE and transferred to Immobilon-FL PVDF membranes (Merck Millipore) using Pierce™ 1-step transfer buffer (Thermo) and the Pierce™ Power Blotter (Thermo). Membranes were washed with ddH_2_O for 5 min, blocked with Odyssey®Blocking Buffer TBS (Li-Cor) for 1 h at room temperature and then incubated with primary antibodies diluted in Odyssey®Blocking Buffer TBS overnight at 4 °C with gentle agitation. KRas (Sigma Aldrich, SAB1404011-100UG), pcRafS338 (CST, 9427), tAkt1 (CST, 2938), pAktS473 (CST, 4060), tErk (CST, 4696), pErkT202/Y204 (CST, 4370), pS6S235/236 (CST, 4858) and β-actin (CST, 4970/ Sigma-Aldrich, A5441) antibodies were used to detect the individual proteins. After primary antibody incubation, membranes were washed three times with TBS-T (50 mM Tris, 150 mM NaCl, 0.05% Tween 20, pH 7.4) for 5 min before being incubated with secondary antibodies (anti-mouse IgG (H+L) (DyLight™ 680 Conjugate) (CST, 5470) / anti-rabbit IgG (H+L) (DyLight™ 800 4X PEG Conjugate) (CST, 5151)) diluted in Odyssey®Blocking Buffer TBS for 1 h at room temperature with gentle agitation. After secondary antibody incubation, the membranes were washed three times for 5 min with TBS-T and then scanned using an Odyssey®CLx imaging system (Li-Cor). Quantification of protein levels from western blots was performed using Empiria Studio (Li-Cor) (Fig. 3B/C, Fig. S18-S21).

## Acknowledgements

This work was co-funded by the Deutsche Forschungsgemeinschaft (DFG; RA 1055/5-1 and GO 284/10-1) and the Drug Discovery Hub Dortmund (DDHD). We thank Nathalie Bleimling, Andreas Arndt and Paul Siebers for invaluable technical assistance. We acknowledge the Paul Scherrer Institut, Villigen, Switzerland for provision of synchrotron radiation beamtime at beamline X10SA of the SLS and would like to thank the staff of the SLS for assistance.

## Competing interests

The authors declare no competing interests.

## Notes

### Competing Interest Statement

The authors have declared no competing interest.

