## Supplementary information for "Targeting oncogenic KRasG13C with nucleotide-based covalent inhibitors"

### Table of Contents


**Experimental Procedures** 4

Synthesis of nucleotide-based covalent inhibitors. 4

Scheme S1. Synthesis scheme of nucleotide-based covalent KRasG13C inhibitors. 4

Scheme S2. Preparation of 5′-phosphorylimidazolidate 2’,3’-O-carbonate of nucleotides. 5

Scheme S3. Preparation of the respective 2’,3’-O-carbamates of nucleotides. 6

Scheme S4. Cleavage of the phosphoramidate. 7

Scheme S5. Insertion of the Michael acceptor or non-reactive acetamido derivative. 8

pKa calculations. 9

Sequence alignment. 9

GAP-stimulated GTP hydrolysis. 9

*In vitro* farnesylation. 10

MS/MS analysis. 10

**Results and Discussion** 12

Calculation of the $K_D$ values of the nucleotide analogues. 12

**Figures and Tables** 14

Figure S1. Rational design targeting KRasG13C. 14

Figure S2. Sequence alignment of Ras small GTPase superfamily. 15

Figure S3. Isocratic separation of the GDP-based nucleotide analogues. 22

Figure S4. Isocratic separation of the GTP-based nucleotide analogues. 23

Figure S5. Isocratic separation of the dGTP-based nucleotide analogues. 24

Figure S6. Isocratic separation of the GppCp-based nucleotide analogues. 25

Figure S7. Time dependent analysis of the covalent modification of KRasG13C. 26

Figure S8. Time dependent analysis of the covalent protein modification with edaGDP or edaGTP. 27

Figure S9. Covalent protein modification with edaGppCp. 28

Figure S10. pH dependent analysis of the covalent protein modification of KRasG13C and KRasWT. 29

Figure S11. Kinetics of the nucleotide association ($k_{on}$). 30

Figure S12. HPLC-based approach for determination of affinities relative to GDP. 31

Figure S13. Structural insights into the binding mode of nucleotide-based covalent inhibitors. 32

Figure S14. GEF-catalysed nucleotide exchange. 33

Figure S15. GAP-stimulated GTP hydrolysis. 34

Figure S16. Labelling and in vitro farnesylation experiments with full-length KRas. 35

Figure S17. MS/MS analysis. 36

Figure S18. In vitro immunoblotting experiments. 37

Figure S19. Western blot analysis and quantification. 38

Figure S20. Electroporation of KRasG13C:acetyledaGDP. 39

Figure S21. Electroporation of KRasG13C-edaGppCp. 40

Table S1. pKa calculations. . 41
Table S2. $k_{on}$ calculations. 42
Table S3. $K_D$ calculations. 43
Table S4. Data collection and refinement statistics for KRasG13C-edaGDP and KRasG13C-bdaGDP. 44

**Author Contributions** 45

**References** 45

### Experimental Procedures

#### Synthesis of nucleotide-based covalent inhibitors

Synthesis of the nucleotide analogues was carried out according to the protocol established by Cremo et al. (1990) and described by Eberth et al. (2005).[1,2]

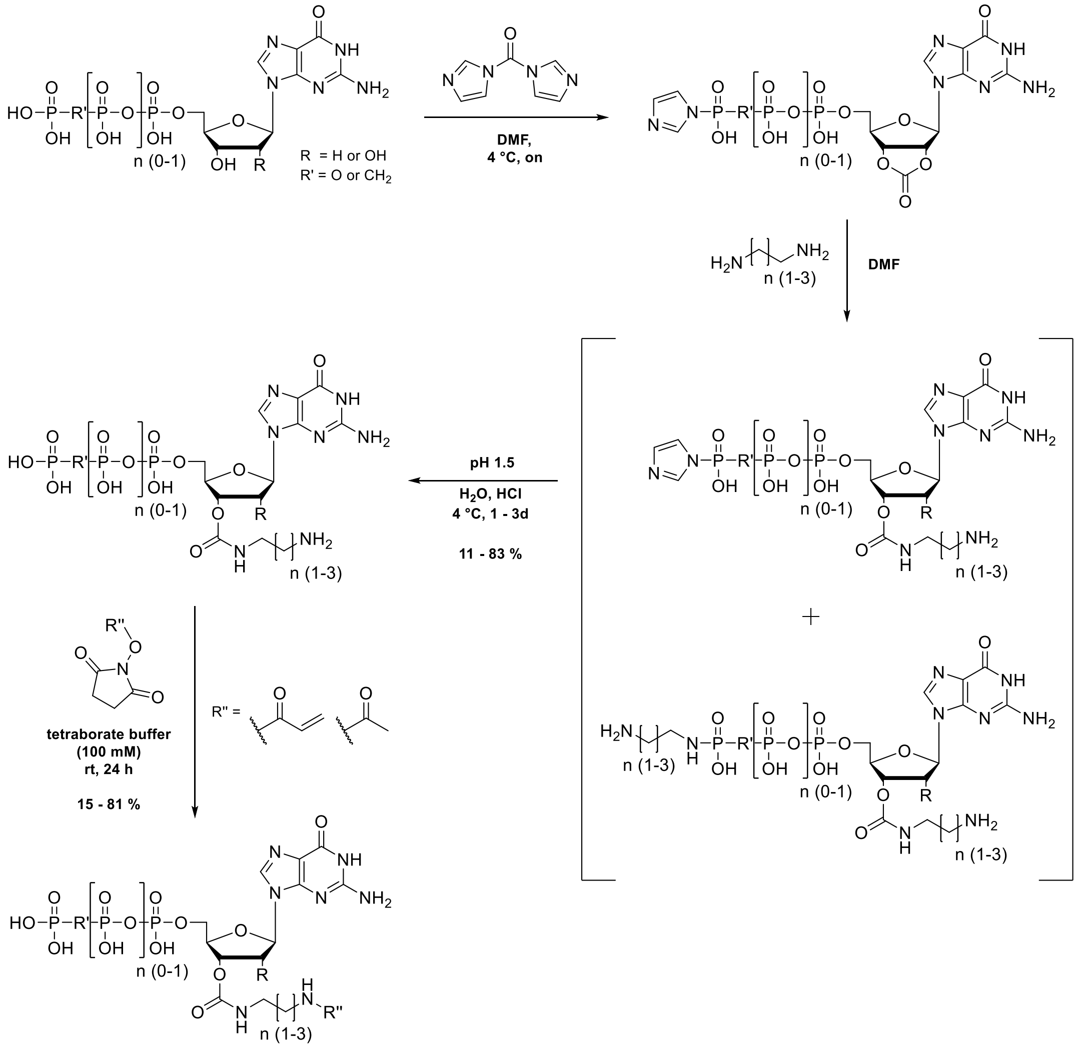


**Scheme S1**. Synthesis scheme of nucleotide-based covalent KRasG13C inhibitors. The nucleotide is mixed with CDI to prepare the 5'-phosphorylimidazolidate 2',3'-O-carbonate followed by the addition of the primary amine to obtain the respective 2',3'-O-carbamate. The phosphoramidate can be cleaved under acidic conditions (pH 1.5) to give the unmodified terminal phosphate group and in a final step the Michael acceptor is attached via an active ester.

A strong cation exchanger (Ion exchanger I, Merck, Darmstadt, Germany) was used to prepare the tributylammonium salts of the nucleotides. The commercially available sodium salt of the nucleotide (0.5 mmol) was dissolved in 3 mL dd$H_2O$ and was applied on the column pre-equilibrated with pyridine/$H_2O$ (1:1). Nucleotide elution was achieved using methanol/$H_2O$ (1:1) and the eluate was dripped into 1 mL TBA. After monitoring the nucleotide elution by spotting samples onto a TLC plate with fluorescent indicator and removing of the methanol/$H_2O$ solution the mixture was dried by repeated rotary evaporation from dry DMF (3 x 20 mL).

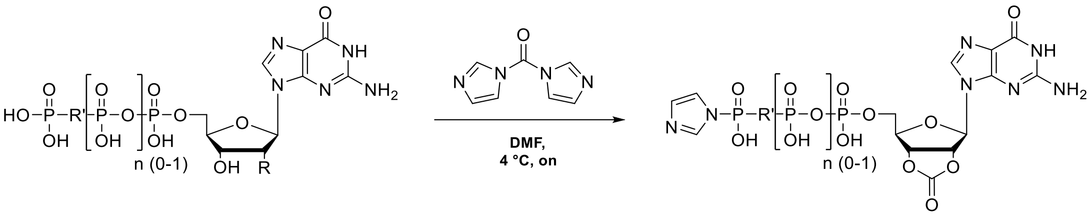


**Scheme S2.** Preparation of 5´-phosphorylimidazolidate 2’,3’-O-carbonate of nucleotides.

**HRMS (ESI-MS):**

| nucleotide | calculated | $[M\text{-}H]^-$ | found |
|---|---|---|---|
| GDP | 518.0232 | $C_{14}H_{14}O_{11}N_7P_2$ | 518.0220 |
| GTP | 597.9895 | $C_{14}H_{15}O_{14}N_7P_3$ | 597.9869 |
| dGTP | 650.0315 | $C_{17}H_{19}O_{13}N_9P_3$ | 650.0304 |
| GppCp* | 545.9828 | $C_{12}H_{15}O_{14}N_5P_3$ | 545.9829 |

*in case of GppCp, the cyclic carbonate was observed with a free terminal phosphate group.

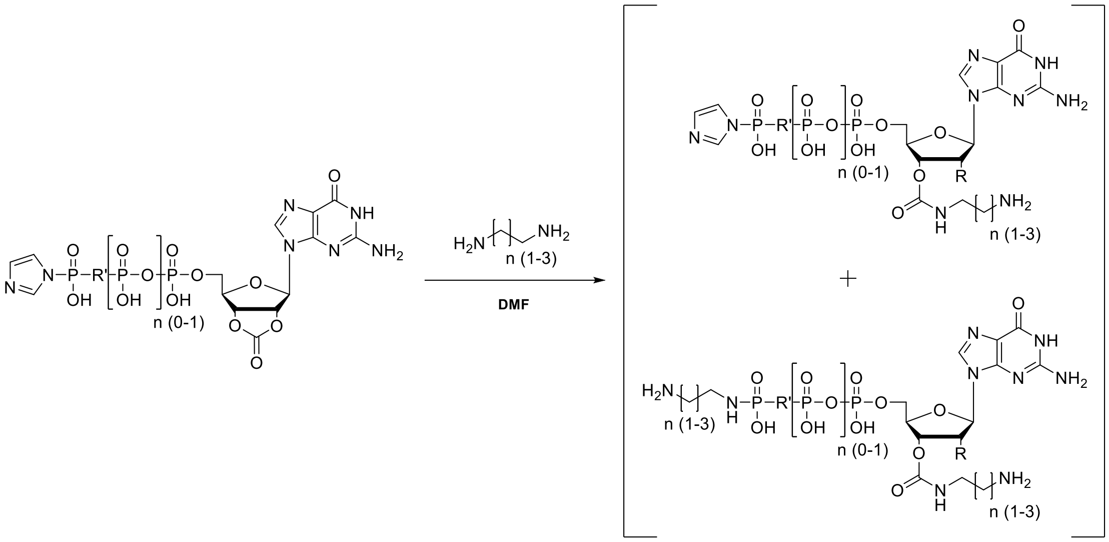


**Scheme S3.** Preparation of the respective 2’,3’-O-carbamates of nucleotides.

The primary amine (eda, pda or bda; 2.5 mmol) dissolved in 5 mL dry DMF was slowly added to the carbonate mixture to prepare the phosphoramidate derivative and the resulting precipitate was recovered by centrifugation (10000 rpm, 10 min) and washed three times with DMF.

**LCMS (ESI-MS):**

| nucleotide | linker | calculated | $[M\text{-}H]^-$ | found |
|---|---|---|---|---|
| GDP | eda | 578.1 | $C_{16}H_{22}N_9O_{11}P_2$ | 578.1 |
| | pda | 592.1 | $C_{17}H_{24}N_9O_{11}P_2$ | 592.1 |
| | bda | 606.1 | $C_{18}H_{26}N_9O_{11}P_2$ | 606.1 |
| GTP | eda | 658.1 | $C_{16}H_{23}N_9O_{14}P_3$ | 658.0 |
| | pda | 672.1 | $C_{17}H_{25}N_9O_{14}P_3$ | 672.1 |
| | bda | 686.1 | $C_{18}H_{27}N_9O_{14}P_3$ | 686.1 |
| dGTP | eda | 634.1 | $C_{15}H_{27}N_9O_{13}P_3$ | 634.1 |
| | pda | 662.1 | $C_{17}H_{31}N_9O_{13}P_3$ | 662.1 |
| | bda | 690.2 | $C_{19}H_{35}N_9O_{13}P_3$ | 690.2 |
| GppCp | eda | 648.1 | $C_{16}H_{29}N_9O_{13}P_3$ | 648.1 |

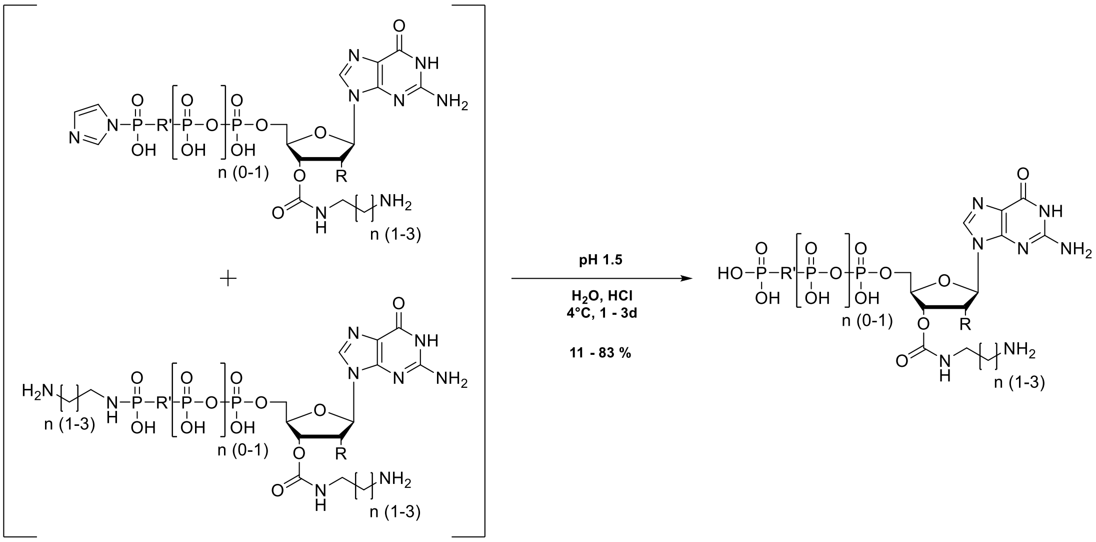


**Scheme S4.** Cleavage of the phosphoramidate.

The solid was dissolved in 20 mL dd$H_2O$ and the pH was adjusted to 1.5 with 0.25 M hydrochloric acid to hydrolyse the phosphoramidate. After stirring at 4 °C for 1-3 d the mixture was then raised to pH 7.5 using 0.25 M NaOH. The nucleotides were purified at 4 °C on a Q-Sepharose column (column volume: 130 mL) preequilibrated with 50 mM triethylammonium bicarbonate buffer (pH 7.6) and eluted by a linear gradient of 50 mM – 1 M TEAB over 600 min with a flow rate of 1 mL/min. The nucleotide containing fractions were analysed by HPLC 50 mM $KP_i$ pH 6.6, 10 mM TBAB, 16% ACN; column: ProntoSIL® 120-5-C18-AQ, Bischoff, Germany) and were lyophilized several times from dd$H_2O$ to remove the buffer

**HRMS (ESI-MS):**

| nucleotide | linker | calculated | $[M-H]^-$ | found | purity (%) | yield (%) |
|---|---|---|---|---|---|---|
| GDP | eda | 528.0651 | $C_{13}H_{20}O_{12}N_7P_2$ | 528.0654 | 90 | 38 |
| | pda | 542.0802 | $C_{14}H_{22}O_{12}N_7P_2$ | 542.0809 | > 95 | 65 |
| | bda | 556.0964 | $C_{15}H_{24}O_{12}N_7P_2$ | 556.0960 | > 95 | 29 |
| GTP | eda | 608.0314 | $C_{13}H_{21}O_{15}N_7P_3$ | 608.0289 | 89 | 56 |
| | pda | 662.0471 | $C_{14}H_{23}O_{15}N_7P_3$ | 662.0471 | 87 | 83 |
| | bda | 636.0621 | $C_{15}H_{25}N_7O_{15}P_3$ | 636.0603 | 94 | 67 |
| dGTP | eda | 592.0359 | $C_{13}H_{21}O_{14}N_7P_3$ | 592.0349 | 84 | 23 |
| | pda | 606.0516 | $C_{14}H_{23}O_{14}N_7P_3$ | 606.0521 | 93 | 11 |
| | bda | 620.0672 | $C_{15}H_{25}N_7O_{14}P_3$ | 620.0663 | 90 | 34 |
| GppCp | eda | 606.0516 | $C_{14}H_{23}N_7O_{14}P_3$ | 606.0513 | 89 | - |

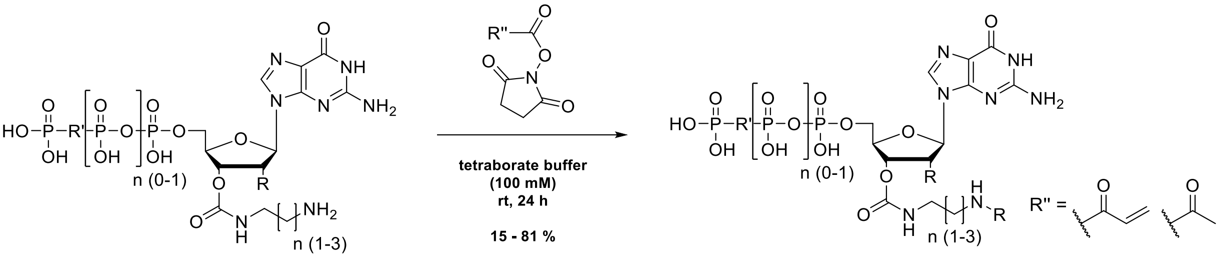

**Scheme S5.** Insertion of the Michael acceptor or non-reactive acetamido derivative.

The nucleotide based covalent inhibitors were prepared by dissolving eda, pda or bda nucleotides in a small amount of tetraborate buffer (100 mM, pH 8.5) and adding N-acryloxysuccinimide (1 eq.) dissolved in 25 µL DMSO. The reaction mixture was stirred for 24 hours at room temperature and the reaction progress was monitored using HPLC. The covalent nucleotide analogues were purified using Q-Sepharose as described above and were stored at -20 °C as concentrated solutions (~ 100 mM) in 200 mM HEPES (pH 7.5).

**HRMS (ESI-MS):**

| **Acrylamides** | **linker** | **calculated** | **$[M-H]^-$** | **found** | **purity (%)** | **yield (%)** |
|---|---|---|---|---|---|---|
| GDP | eda | 582.0751 | $C_{16}H_{22}N_7O_{13}P_2$ | 582.0741 | > 95 | 57 |
| | pda | 596.0907 | $C_{17}H_{24}N_7O_{13}P_2$ | 596.0907 | 87 | 65 |
| | bda | 610.1064 | $C_{18}H_{26}N_7O_{13}P_2$ | 610.1062 | > 95 | 78 |
| GTP | eda | 662.0414 | $C_{16}H_{23}N_7O_{16}P_3$ | 662.0394 | 93 | 54 |
| | pda | 676.0571 | $C_{17}H_{25}N_7O_{16}P_3$ | 676.0551 | > 95 | 63 |
| | bda | 690.0727 | $C_{18}H_{27}N_7O_{16}P_3$ | 690.0705 | 86 | 75 |
| dGTP | eda | 646.0465 | $C_{16}H_{23}N_7O_{15}P_3$ | 646.0452 | 93 | 63 |
| | pda | 660.0621 | $C_{17}H_{25}N_7O_{15}P_3$ | 660.0611 | 90 | 81 |
| | bda | 674.0778 | $C_{18}H_{27}N_7O_{15}P_3$ | 674.0771 | 93 | 74 |
| GppCp | eda | 660.0621 | $C_{17}H_{27}N_7O_{16}P_3$ | 660.0619 | 79 | 15* |
| **Acetylamides** | **linker** | **calculated** | **$[M-H]^-$** | **found** | **purity (%)** | **yield (%)** |
| GDP | eda | 570.0751 | $C_{15}H_{22}N_7O_{13}P_2$ | 570.0734 | > 95 | 57 |
| | pda | 584.0907 | $C_{16}H_{24}N_7O_{13}P_2$ | 584.0889 | 90 | 51 |
| | bda | 598.1064 | $C_{17}H_{26}N_7O_{13}P_2$ | 598.1042 | 79 | 73 |
| GTP | eda | 650.0414 | $C_{15}H_{23}N_7O_{16}P_3$ | 650.0392 | > 95 | 23 |
| | pda | 664.0571 | $C_{16}H_{25}N_7O_{16}P_3$ | 664.0553 | 94 | 40 |
| | bda | 678.0727 | $C_{17}H_{27}N_7O_{16}P_3$ | 678.0700 | 93 | 31 |

*in case of GppCp, the total yield was calculated over 4 steps.

**pKa calculations**

For the protein pKa prediction, a program from OpenEye based on the Zap finite difference Poisson–Boltzmann solver was used. Regarding partial charges of the protein, Delphi radii and CHARMM36 All-Hydrogen partial charges were utilized. The linker in the bda/edaGDP structures were removed. The am1bccsym method was used to assign appropriate partial charges to the ligands. An inner dielectric of 10, an ionic strength of 0.05 M and ionization (i.e. pH) of 7.5 was applied. For cysteine, a reference pKa of 8.6 was used. All hydrogen atoms were explicitly modeled, and except for the orientations of the OH and SH protons, which were sampled in 10° steps, the rest of the structure was static. Optimizing ionization state and SH orientation was achieved by applying ten million Monte Carlo steps.[3]

**Sequence alignment**

A multiple sequence alignment of the G domain Ras superfamily members was performed using Uniprot.[4] The uniport accession codes that were used for the sequence alignment of the GTPases can be taken from Fig. S2.

**GAP-stimulated GTP hydrolysis**

First, for KRasWT$_{1\text{-}169}$, nucleotide exchange with GTP was performed at pH 7.5, and for KRasG13C$_{1\text{-}169}$, covalent modification with a 10-fold excess of acryl-edaGTP at pH 9.5 was done, both in the presence of 50 mM EDTA to block intrinsic GTP hydrolysis. After incubation, the Ras proteins were washed 5 times with buffer (20 mM HEPES (pH 7.5), 100 mM NaCl, and 1 mM TCEP) using centrifugal filter devices to remove any unbound nucleotide. Nucleotide exchange of KRasWT$_{1\text{-}169}$ was controlled by isocratic HPLC runs, and covalent modification of KRasG13C$_{1\text{-}169}$ was verified by ESI-MS. GTP hydrolysis of Ras proteins was initiated by addition of 2 mM $MgCl_2$ in the absence or presence of Ras-GAP (1:1000 for KRasWT$_{1\text{-}169}$:GTP and 1:1 for KRasG13C1-169-edaGTP). At defined time points (0, 5, 10, 15, 20, 30, 45, 60, 90 and 120 min), samples were taken and immediately snap-frozen in liquid nitrogen. For KRasWT$_{1\text{-}169}$:GTP, samples were thawed and denatured for 5 min at 95 °C. Samples were centrifuged at 14.000 rpm for 10 min at 4 °C and subsequently analyzed by isocratic HPLC runs. For KRasG13C$_{1\text{-}169}$-edaGTP, samples were centrifuged and analyzed by ESI-MS. The relative amount of each nucleotide was determined by integrating the area of the GTP and GDP peaks using Origin (Fig. S15).[5]

#### *In vitro* farnesylation

50 µM of full-length KRas protein was mixed with 250 µM farnesyl pyrophosphate (FPP) and 10 µM farnesyltransferase (FTase). After incubation at room temperature for 1h the mixture was centrifugated at 14.000 rpm for 10 min at 4 °C and analyzed via ESI-MS (Fig. S16).

#### MS/MS analysis

For MS/MS analysis, the samples were dissolved in 100 mM TEAB and incubated for 1 h at 55 °C in the presence of 10 mM TCEP. 17 mM iodoacetamide was added and the samples were incubated in the dark at room temperature for 30 min. Samples were precipitated by adding pre-chilled acetone and stored overnight at -20 °C. After drying the pellets, Trypsin (Roche) was added and the samples were digested at 37 °C with 300 rpm shaking overnight. The digestion was quenched by the addition of 2% TFA and a stage tip purification[6] was performed, samples were evaporated to dryness and stored at -20 °C until MS/MS analysis. For nanoHPLC-MS/MS analysis samples were dissolved in 20 µL of 0.1% TFA in water and 3 µL were injected onto an UltiMateTM 3000 RSLCnano system (ThermoFisher Scientific, Germany) online coupled to a Q Exactive™ Plus Hybrid Quadrupole-Orbitrap Mass Spectrometer equipped with a nanospray source (Nanospray Flex Ion Source, Thermo Scientific). All solvents were LC-MS grade. Samples were injected onto a pre-column cartridge (5 µm, 100 Å, 300 µm ID * 5 mm, Dionex, Germany) using 0.1% TFA in water as eluent with a flow rate of 30 µL/min. Desalting was performed for 5 min with eluent flow to waste followed by back-flushing of the sample during the whole analysis from the pre-column to the PepMap100 RSLC C18 nano-HPLC column (2 µm, 100 Å, 75 µm ID × 50 cm, nanoViper, Dionex, Germany) using a linear gradient starting with 95% solvent A (water containing 0.1% formic acid) / 5% solvent B (acetonitrile containing 0.1% formic acid) and increasing to 30% solvent B in 90 min using a flow rate of 300 nL/min. Afterwards, the column was washed (two steps 60 and 95% solvent B) and re-equilibrated to starting conditions. The nanoHPLC was online coupled to the Quadrupole-Orbitrap Mass Spectrometer using a standard coated SilicaTip (ID 20 µm, Tip-ID 10 µM, New Objective, Woburn, MA, USA). Mass range of m/z 300 to 1650 was acquired with a resolution of 70000 for a full scan, followed by up to 10 high energy collision dissociation (HCD) MS / MS scans of the most intense at least doubly charged ions using a resolution of 17500 and a NCE energy of 25%. Data evaluation was performed using MaxQuant software[7] (v.1.6.3.4) including the Andromeda search algorithm[8] and searching the KRas sequence together with a database containing typical contaminants like keratins, trypsin etc., which is included in the MaxQuant software. The search was performed for full enzymatic trypsin cleavages allowing two miscleavages. For database search oxidation of methionine and N-terminal acetylation of proteins, carbamidomethylation of cysteines, and artificial modification of cysteines were defined as variable modifications. The mass accuracy for full mass spectra was set to 20 ppm (first search) and 4.5 ppm (second search), respectively and for MS/MS spectra to 20 ppm. The false discovery rates for peptide and protein identification were set to 1%. For further analysis, the peptide intensities of KRas were compared for modified and unmodified KRas. (Fig. S17).

#### Results and Discussion

##### Calculation of the $K_D$ values of the nucleotide analogues

Based on the percentage distribution determined by the HPLC assay, the relative association constants ($K_{relA}$) for pdaGDP and bdaGDP could subsequently be calculated. In contrast, the superposition of the GDP signal prevented an exact determination of the relative association constant for edaGDP. $K_{relA}$ values of 0.34 and 0.28 relative to GDP were determined for pdaGDP and bdaGDP, respectively. Finally, considering the $K_D$ value of 2.5 pM for GDP described by Jeganathan *et al.* the corresponding dissociation constant $K_D$ values could be determined, which are shown in Tab. S3.[9]

$$Ras + GDP \xrightarrow{k_{GDP}} Ras{:}GDP \tag{1a}$$

$$Ras + aGDP \xrightarrow{k_{aGDP}} Ras{:}aGDP \tag{1b}$$

By rearranging the equilibrium reactions listed in (1a) and (1b), the following relationships are obtained for $K_{GDP}$ (2a) and $K_{aGDP}$ (2b):

$$K_{GDP} = \frac{[Ras{:}GDP]}{[Ras][GDP]} \tag{2a}$$

$$K_{aGDP} = \frac{[Ras{:}aGDP]}{[Ras][aGDP]} \tag{2b}$$

Assuming that the concentration for aGDP used in the HPLC assay is 50 µM, the following GDP concentrations for pdaGDP (3a) and bdaGDP (3b) can be determined at time t=0 h using the percentages determined in Fig. S12:

$$GDP_{total} = \frac{\%_{GDP}}{\%_{pdaGDP}} = \frac{37\ \%}{63\ \%} \times 50\ \mu M = 29.4\ \mu M \tag{3a}$$

$$GDP_{total} = \frac{\%_{GDP}}{\%_{bdaGDP}} = \frac{43\ \%}{57\ \%} \times 50\ \mu M = 37.7\ \mu M \tag{3b}$$

The concentrations of the components listed in equation (2b) can be calculated as follows for pdaGDP (4a-d) and bdaGDP (5a-d) by considering the total GDP concentrations calculated previously:

$$[Ras{:}pdaGDP] = \frac{47}{100} \times 29.4\ \mu M = 13.8\ \mu M \tag{4a}$$

$$[Ras{:}GDP] = \frac{53}{100} \times 29.4\ \mu M = 15.6\ \mu M \tag{4b}$$

$$[pdaGDP] = 50\ \mu M - 13.8\ \mu M = 36.2\ \mu M \tag{4c}$$

$$[GDP] = 29.4\ \mu M - 15.6\ \mu M = 13.8\ \mu M \tag{4d}$$

$$[Ras{:}\,bdaGDP] = \frac{39.5}{100} \times 37.7\ \mu M = 14.9\ \mu M \tag{5a}$$

$$[Ras{:}\,GDP] = \frac{60.5}{100} \times 37.7\ \mu M = 22.8\ \mu M \tag{5b}$$

$$[bdaGDP] = 50\ \mu M - 14.9\ \mu M = 35.1\ \mu M \tag{5c}$$

$$[GDP] = 37.7\ \mu M - 22.8\ \mu M = 14.9\ \mu M \tag{5d}$$

The relative association constant $K_{relA}$ can be calculated by the following formula (6):

$$K_{relA} = \frac{K_{aGDP}}{K_{GDP}} = \frac{[Ras{:}\,aGDP] \times [GDP]}{[aGDP] \times [Ras{:}\,GDP]} \tag{6}$$

By inserting the concentrations of each component determined from (4a-d) and (5a-d) into (6), the relative association constants $K_{relA}$ for pdaGDP (7a) and bdaGDP (7b) can be calculated:

$$K_{relA} = \frac{K_{pdaGDP}}{K_{GDP}} = \frac{13.8\ \mu M \times 13.8\ \mu M}{36.2\ \mu M \times 15.6\ \mu M} = 0.34 \tag{7a}$$

$$K_{relA} = \frac{K_{bdaGDP}}{K_{GDP}} = \frac{14.9\ \mu M \times 14.9\ \mu M}{35.1\ \mu M \times 22.8\ \mu M} = 0.28 \tag{7b}$$

Based on the relative association constants $K_{relA}$, the $K_A$ values of the nucleotide analogues can be calculated by multiplication with the association constant of GDP:

$$K_A = K_A GDP \times K_{relA} \tag{8}$$

The $K_D$ value of 2.5 pM described by Jeganathan *et al.* was used as the reference value for GDP.[9] By substituting the reference value into $K_A$=1/$K_D$, a $K_A$ value for GDP of 0.4 $pM^{-1}$ was determined. Accordingly, the $K_A$ values for pdaGDP (9a) and bdaGDP (9b) are as follows:

$$K_A pdaGDP = 0.4\, pM^{-1} \times 0.34 = 0.136\, pM^{-1} \tag{9a}$$

$$K_A bdaGDP = 0.4\, pM^{-1} \times 0.28 = 0.112\, pM^{-1} \tag{9b}$$

By converting the determined $K_A$ values into the corresponding $K_D$ values using $K_D$=1/$K_A$, the following dissociation constants could be determined for pdaGDP (10a) and bdaGDP (10b):

$$K_D pdaGDP = \frac{1}{0.136\, pM^{-1}} = 7.4\, pM \tag{10a}$$

$$K_D bdaGDP = \frac{1}{0.112\, pM^{-1}} = 8.9\, pM \tag{10b}$$

### Figures and Tables

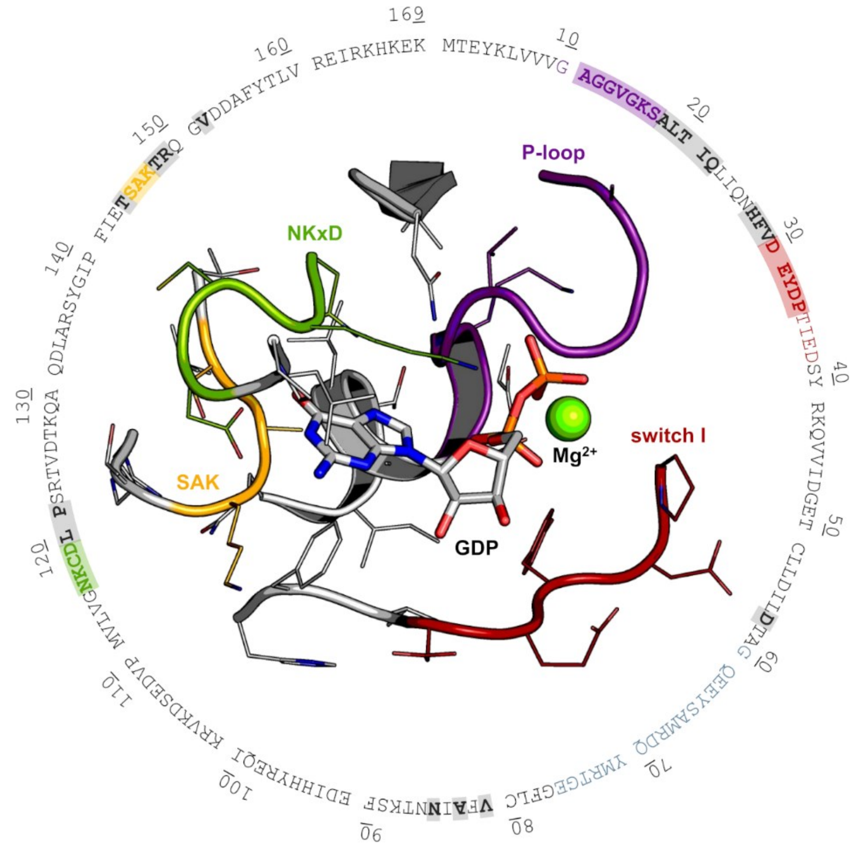


**Figure S1.** Rational design targeting KRasG13C. Based on the crystal structure of KRasWT (PDB 4obe), aa residues that are potentially accessible via the linker design (eda: ~10 Å, bda: ~12 Å) are marked (grey, P-loop: violet, switch I: red, NKxD: green, SAK: yellow) and used for multiple sequence alignment of the Ras small GTPase superfamily (Fig. S2).

```
ARF1     P84077 ------------------------------------MGNIFAN---------LFKGLFGKKEMRILMVGLDAAGKTTILYKLKLG--------------------EI-VTTIPTIG---- 50
ARF3     P61204 ------------------------------------MGNIFGN---------LLKSLIGKKEMRILMVGLDAAGKTTILYKLKLG--------------------EI-VTTIPTIG---- 50
ARF4     P18085 ------------------------------------MGLTISS---------LFSRLFGKKQMRILMVGLDAAGKTTILYKLKLG--------------------EI-VTTIPTIG---- 50
ARF5     P84085 ------------------------------------MGLTVSA---------LFSRIFGKKQMRILMVGLDAAGKTTILYKLKLG--------------------EI-VTTIPTIG---- 50
ARF6     P62330 ----------------------------------------MGK---------VLSKIFGNKEMRILMLGLDAAGKTTILYKLKLG--------------------QS-VTTIPTVG---- 46
ARL1     P40616 ------------------------------------MGGFFSS---------IFSSLFGTREMRILILGLDGAGKTTILYRLQVG--------------------EV-VTTIPTIG---- 50
ARL2     P36404 ------------------------------------MGLLTIL---------KKMKQ-KERELRLLMLGLDNAGKTTILKKFNGE--------------------DI-DTISPTLG---- 49
ARL3     P36405 ------------------------------------MGLLSIL---------RKLKSAPDQEVRILLLGLDNAGKTTLLKQLASE--------------------DI-SHITPTQG---- 50
ARL4A    P40617 ------------------------------------MGNGLSD----Q-TS-ILSNLPSFQSFHIVILGLDCAGKTTVLYRLQFN--------------------EF-VNTVPTKG---- 53
ARL4C    P56559 -----------------------------------------------M-GN-ISSNISAFQSLHIVMLGLDSAGKTTVLYRLKFN--------------------EF-VNTVPTIG---- 46
ARL4D    P49703 ------------------------------------MGNHLTE----M-APTASSFLPHFQALHVVVIGLDSAGKTSLLYRLKFK--------------------EF-VQSVPTKG---- 54
ARL5A    Q9Y689 ------------------------------------MGILFTR----------IWRLFNHQEHKVIIVGLDNAGKTTILYQFSMN--------------------EV-VHTSPTIG---- 49
ARL5B    Q96KC2 ------------------------------------MGLIFAK----------LWSLFCNQEHKVIIVGLDNAGKTTILYQFLMN--------------------EV-VHTSPTIG---- 49
ARL5C    A6NH57 ------------------------------------MGQLIAK----------LMSIFGNQEHTVIIVGLDNEGKTTILYRFLTN--------------------EV-VHMCPTIG---- 49
ARL6     Q9H0F7 ------------------------------------MGLLDRL---------SVLLGLKKKEVHVLCLGLDNSGKTTIINKLKPS-----------------N--AQS-QNILPTIG---- 52
ARL8A    Q96BM9 ------------------------------------MIALFNK----L--LDWFKALFWKEEMELTLVGLQYSGKTTFVNVIASG--------------------QFNEDMIPTVG---- 54
ARL8B    Q9NVJ2 ------------------------------------MLALISR----L--LDWFRSLFWKEEMELTLVGLQYSGKTTFVNVIASG--------------------QFSEDMIPTVG---- 54
ARL9     Q6T311 ------------------------------------MRPTWK--------ALSHPAWPEEKNKQILVLGLDGAGKTSVLHSLASN--------------------RVQHSVAPTQG---- 52
ARL10    Q8N8L6 ---------------D---RGEAWWGA------EAARLPEWDE-WDPEDEEDEEPALEELEQREVLVLGLDGAGKSTFLRVLSGK--------------------PPLEGHIPTWG---- 111
ARL11    Q969Q4 ------------------------------------MGSV--------------NSRGHKAEAQVVMMGLDSAGKTTLLYKLKGH--------------------QL-VETLPTVG---- 45
ARL13A   Q5H913 ------------------------------------MFRLLSSCCSCL-----RTTEETRRNVTIPIIGLNNSGKTVLVEAFQKL--------------------LP-SKTDHCMK---- 54
ARL13B   Q3SXY8 ------------------------------------MFSLMASCCGWF-----KRWREPVRKVTLLMVGLDNAGKTATAKGIQGE--------------------YP-EDVAPTVG---- 54
ARL14    Q8N4G2 ------------------------------------MGSLG-------------SKNPQTKQAQVLLLGLDSAGKSTLLYKLKLA--------------------KD-ITTIPTIG---- 46
ARL15    Q9NXU5 --------------------------I--------TEAFLYMD-YLCFRALCCKGPPPARPEYDLVCIGLTGSGKTSLLSKLCSE--------------------SP-DNVVSTTG---- 65
ARL16    Q0P5N6 --------------------------V--------AGGRALSR------GA-ELRVPGGAKHGMCLLLGATGVGKTLLVKRLQEV--SS--R------DGKGDLGEP-PPTRPTVG---- 66
ARL17    Q8IVW1 ------------------------------------MGNIFEK---------LFKSLLGKKKMRILILSLDTAGKTTILYKLKLG--------------------ET-VPAVPTVG---- 50
ARFRP1   Q13795 ------------------------------------MYTLLSG---------LYKYMFQKDEYCILILGLDNAGKTTFLEQSKTR--FN--K------NYKG--MSL-SKITTTVG---- 58
SAR1A    Q9NR31 ------------------------------------MSFIFEWIYNGF-SSVLQFLGLYKKSGKLVFLGLDNAGKTTLLHMLKDD--------------------RL-GQHVPTLH---- 58
SAR1B    Q9Y6B6 ------------------------------------MSFIFDWIYSGF-SSVLQFLGLYKKTGKLVFLGLDNAGKTTLLHMLKDD--------------------RL-GQHVPTLH---- 58
TRIM23   P36406 --------------------------L------DASIPVTFTK--------DNRVHIGPKMEIRVVTLGLDGAGKTTILFKLKQD--------------------EF-MQPIPTIG---- 437
IFT27    Q9BW83 ---------------------------------------------------------MVKLAAKCILAGDPAVGKTALAQIFRSDGAHF--Q-K----SY-----------TLTTG--MD 43
RAB1A    P62820 ---------------------------------------------------MSSMNPEYDYLFKLLLIGDSGVGKSCLLLRFADD--TY--T-E----SY-----------ISTIG--VD 47
RAB1B    Q9H0U4 ------------------------------------------------------MNPEYDYLFKLLLIGDSGVGKSCLLLRFADD--TY--T-E----SY-----------ISTIG--VD 44
RAB1C    Q92928 ------------------------------------------------------MNPGYDCLFKLLLIGDSGVGKSCLLLRFADD--PY--T-E----SY-----------ISTIG--VD 44
RAB2A    P61019 --------------------------------------------------------MAYAYLFKYIIIGDTGVGKSCLLLQFTDK--RF--Q-P----VH-----------DLTIG--VE 42
RAB2B    Q8WUD1 --------------------------------------------------------MTYAYLFKYIIIGDTGVGKSCLLLQFTDK--RF--Q-P----VH-----------DLTIG--VE 42
RAB3A    P20336 ------------------------------AS-------AT----DSRYGQKESSDQNFDYMFKILIIGNSSVGKTSFLFRYADD--SF--T-P----AF-----------VSTVG--ID 58
RAB3B    P20337 ------------------------------AS-------VT----DGKTGVKDASDQNFDYMFKLLIIGNSSVGKTSFLFRYADD--TF--T-P----AF-----------VSTVG--ID 58
RAB3C    Q96E17 ------------------------------AS-------AQ----DARYGQKDSSDQNFDYMFKLLIIGNSSVGKTSFLFRYADD--SF--T-S----AF-----------VSTVG--ID 66
RAB3D    O95716 ------------------------------AS-------AG----DTQAGPRDAADQNFDYMFKLLLIGNSSVGKTSFLFRYADD--SF--T-P----AF-----------VSTVG--ID 58
RAB4A    P20338 -------------------------------------------------MSQTAMSETYDFLFKFLVIGNAGTGKSCLLHQFIEK--KF--K-D----DS-----------NHTIG--VE 49
RAB4B    P61018 ------------------------------------------------------MAETYDFLFKFLVIGSAGTGKSCLLHQFIEN--KF--K-Q----DS-----------NHTIG--VE 44
RAB5A    P20339 -------------------------------------MASR----GA-TRPNGPNTGNKICQFKLVLLGESAVGKSSLVLRFVKG--QF--H-E----FQ-----------ESTIG--AA 56
RAB5B    P61020 -------------------------------------MTSR----ST-ARPNGQPQASKICQFKLVLLGESAVGKSSLVLRFVKG--QF--H-E----YQ-----------ESTIG--AA 56
RAB5C    P51148 ------------------------------------MAGRG----GA-ARPNGPAAGNKICQFKLVLLGESAVGKSSLVLRFVKG--QF--H-E----YQ-----------ESTIG--AA 57
RAB6A    P20340 ------------------------------------MSTGG----DFG---------NPLRKFKLVFLGEQSVGKTSLITRFMYD--SF--D-N----TY-----------QATIG--ID 49
RAB6B    Q9NRW1 ------------------------------------MSAGG----DFG---------NPLRKFKLVFLGEQSVGKTSLITRFMYD--SF--D-N----TY-----------QATIG--ID 49
RAB6C    Q9H0N0 ------------------------------------MSAGG----DFG---------NPLRKFKLVFLGEQSVAKTSLITRFRYD--SF--D-N----TY-----------QAIIG--ID 49
RAB7A    P51149 ------------------------------------------------------MTSRKKVLLKVIILGDSGVGKTSLMNQYVNK--KF--S-N----QY-----------KATIG--AD 44
RAB7B    Q96AH8 ------------------------------------------------------MNPRKKVDLKLIIVGAIGVGKTSLLHQYVHK--TF--Y-E----EY-----------QTTLG--AS 44
RAB8A    P61006 ------------------------------------------------------MAKTYDYLFKLLLIGDSGVGKTCVLFRFSED--AF--N-S----TF-----------ISTIG--ID 44
RAB8B    Q92930 ------------------------------------------------------MAKTYDYLFKLLLIGDSGVGKTCLLFRFSED--AF--N-T----TF-----------ISTIG--ID 44
RAB9A    P51151 -------------------------------------------------------MAGKSSLFKVILLGDGGVGKSSLMNRYVTN--KF--D-T----QL-----------FHTIG--VE 43
RAB9B    Q9NP90 -------------------------------------------------------MSGKSLLLKVILLGDGGVGKSSLMNRYVTN--KF--D-S----QA-----------FHTIG--VE 43
RAB10    P61026 -----------------------------------------------------MAKKTYDLLFKLLLIGDSGVGKTCVLFRFSDD--AF--N-T----TF-----------ISTIG--ID 45
RAB11A   P62491 ---------------------------------------------------MGTRDDEYDYLFKVVLIGDSGVGKSNLLSRFTRN--EF--N-L----ES-----------KSTIG--VE 47
RAB11B   Q15907 ---------------------------------------------------MGTRDDEYDYLFKVVLIGDSGVGKSNLLSRFTRN--EF--N-L----ES-----------KSTIG--VE 47
RAB12    Q6IQ22 ---------------------------------------GG----QGR--RRKQPPRPADFKLQVIIIGSRGVGKTSLMERFTDD--TF--C-E----AC-----------KSTVG--VD 78
RAB13    P51153 ------------------------------------------------------MAKAYDHLFKLLLIGDSGVGKTCLIIRFAED--NF--N-N----TY-----------ISTIG--ID 44
RAB14    P61106 ---------------------------------------------------MATAPYNYSYIFKYIIIGDMGVGKSCLLHQFTEK--KF--M-A----DC-----------PHTIG--VE 47
RAB15    P59190 ------------------------------------------------------MAKQYDVLFRLLLIGDSGVGKTCLLCRFTDN--EF--H-S----SH-----------ISTIG--VD 44
RAB17    Q9H0T7 ------------------------------------MAQAH----R---TPQPRAAPSQPRVFKLVLLGSGSVGKSSLALRYVKN--DF--K-S-----I-----------LPTVG--CA 54
RAB18    Q9NP72 ------------------------------------------------------MDEDVLTTLKILIIGESGVGKSSLLLRFTDD--TF--D-P----EL-----------AATIG--VD 44
RAB19    A4D1S5 ---------------------------------------------MHFSSSARAADENFDYLFKIILIGDSNVGKTCVVQHFKSG--VY--T-E----TQ-----------QNTIG--VD 53
RAB20    Q9NX57 ---------------------------------------------------------MRKPDSKIVLLGDMNVGKTSLLQRYMER--RF--P------DT-----------VSTVG--GA 40
RAB21    Q9UL25 ------------------------------------MAAAG----G---GGGGAAAAGRAYSFKVVLLGEGCVGKTSLVLRYCEN--KF--N-D----KH-----------ITTLQ--AS 55
RAB22A   Q9UL26 ---------------------------------------------------------MALRELKVCLLGDTGVGKSSIVWRFVED--SF--D-P----NI-----------NPTIG--AS 41
RAB23    Q9ULC3 -----------------------------------------------------MLEEDMEVAIKMVVVGNGAVGKSSMIQRYCKG--IF--T-K----DY-----------KKTIG--VD 45
RAB24    Q969Q5 -------------------------------------------------------MSGQRVDVKVVMLGKEYVGKTSLVERYVHD--RF--LVG----PY-----------QNTIG--AA 44
RAB25    P57735 --------------------------------------------------MGNGTEEDYNFVFKVVLIGESGVGKTNLLSRFTRN--EF--S-H----DS-----------RTTIG--VE 48
RAB26    Q9ULW5 ---------------------------------------PL----QPGRPSLGGGVDFYDVAFKVMLVGDSGVGKTCLLVRFKDG--AF--LAG----TF-----------ISTVG--ID 100
RAB27A   P51159 -----------------------------------------------------MSDGDYDYLIKFLALGDSGVGKTSVLYQYTDG--KF--N-S----KF-----------ITTVG--ID 45
RAB27B   O00194 -----------------------------------------------------MTDGDYDYLIKLLALGDSGVGKTTFLYRYTDN--KF--N-P----KF-----------ITTVG--ID 45
RAB28    P51157 --------------------------------------------------MSDSEEESQDRQLKIVVLGDGASGKTSLTTCFAQE--TF--G-K----QY-----------KQTIG--LD 48
RAB29    O14966 -------------------------------------------------------MGSRDHLFKVLVVGDAAVGKTSLVQRYSQD--SF--S-K----HY-----------KSTVG--VD 43
RAB30    Q15771 -----------------------------------------------------MSMEDYDFLFKIVLIGNAGVGKTCLVRRFTQG--LF--P-P----GQ-----------GATIG--VD 45
RAB31    Q13636 ---------------------------------------------------------MAIRELKVCLLGDTGVGKSSIVCRFVQD--HF--D-H----NI-----------SPTIG--AS 41
RAB32    Q13637 ---------------------------------------GD----PGLGAAAAPAPETREHLFKVLVIGELGVGKTSIIKRYVHQ--LF--S-Q----HY-----------RATIG--VD 61
RAB33A   Q14088 ------------------------------AA-------GL----ASLELDSSLDQYVQIRIFKIIVIGDSNVGKTCLTFRFCGG--TF--P-D----KT-----------EATIG--VD 72
RAB33B   Q9H082 ------------------------------FS-------SS----GAVSGASGFLPPARSRIFKIIVIGDSNVGKTCLTYRFCAG--RF--P-D----RT-----------EATIG--VD 69
RAB34    Q9BZG1 --PVRR---D--RVLA---ELPQCLRKEAALHGHKDFH-PR----VTCACQEHRTGTVGFKISKVIVVGDLSVGKTCLINRFCKD--TF--D-K----NY-----------KATIG--VD 88
RAB35    Q15286 ------------------------------------------------------MARDYDHLFKLLIIGDSGVGKSSLLLRFADN--TF--S-G----SY-----------ITTIG--VD 44
RAB36    O95755 --PVSR---D--RVIA---SFPKWYTPEACLQLREHFH-GQ----VSAACQRRNTGTVGLKLSKVVVVGDLYVGKTSLIHRFCKN--VF--D-R----DY-----------KATIG--VD 159
RAB37    Q96AX2 -------------------------------------------------P---PCSPSYDLTGKVMLLGDTGVGKTCFLIQFKDG--AF--LSG----TF-----------IATVG--ID 66
RAB38    P57729 -----------------------------------------------------MQAPHKEHLYKLLVIGDLGVGKTSIIKRYVHQ--NF--S-S----HY-----------RATIG--VD 45
RAB39A   Q14964 ------------------------------------------------------METIWIYQFRLIVIGDSTVGKSCLLHRFTQG--RF--PG-LRSPAC-----------DPTVG--VD 48
RAB39B   Q96DA2 ------------------------------------------------------MEAIWLYQFRLIVIGDSTVGKSCLIRRFTEG--RF--AQ-V----S-----------DPTVG--VD 44
RAB40A   Q8WXH6 ------------------------------------------------MSAPGSPDQAYDFLLKFLLVGDRDVGKSEILESLQDG--AA--E-S----PY-----------SHLGG--ID 50
RAB40B   Q12829 ------------------------------------------------MSALGSPVRAYDFLLKFLLVGDSDVGKGEILASLQDG--AA--E-S----PY-----------GHPAG--ID 50
RAB40C   Q96S21 ------------------------------------------------MGSQGSPVKSYDYLLKFLLVGDSDVGKGEILESLQDG--AA--E-S----PY-----------AYSNG--ID 50
RAB41    Q5JT25 ---------------------------MSAFGHDEAWMEAG----GFGLEAAERTEYQSLCKSKLLFLGEQSVGKTSIISRFMYN--SF--G-C----AC-----------QATVG--ID 67
RAB42    Q8N4Z0 -----------------------------------------------------MEAEGCRYQFRVALLGDAAVGKTSLLRSYVAG--AP--GAPEPEPEP-----------EPTVG--AE 50
RAB43    Q86YS6 ----------------------------------------M----AGPGPGPGDPDEQYDFLFKLVLVGDASVGKTCVVQRFKTG--AF--S-E----RQ-----------GSTIG--VD 54
RAB44    Q7Z6P3 --PHSR---E--PRAESRLEDPGMDSREAGLT-------PS----PGDPMAGGGPQANPDYLFHVIFLGDSNVGKTSFLHLLHQN--SF--A-T----GL-----------TATVG--VD 869
RASEF    Q8IZ41 ------------------------------AL-------SP----QTDLVDDNAKSFSSQKAYKIVLAGDAAVGKSSFLMRLCKN--EF--R-E----NI-----------SATLG--VD 577
DIRAS1   O95057 -------------------------------------------------------MPEQSNDYRVVVFGAGGVGKSSLVLRFVKG--TF--R-D----TY-----------IPTIE--DT 43
DIRAS2   Q96HU8 -------------------------------------------------------MPEQSNDYRVAVFGAGGVGKSSLVLRFVKG--TF--R-E----SY-----------IPTVE--DT 43
DIRAS3   O95661 LKRLRL------------LP--------------------------ALLILRAFKPHRKIRDYRVVVVGTAGVGKSTLLHKWASG--NF--R-H----EY-----------LPTIE--NT 73
ERAS     Q7Z444 -KPGTF-----------DLGLATWSPSFQG---------ET-----HRAQARRRDVGRQLPEYKAVVVGASGVGKSALTIQLNHQ--CF--V-E----DH-----------DPTIQ--DS 77
HRAS     P01112 -----------------------------------------------------------MTEYKLVVVGAGGVGKSALTIQLIQN--HF--V-D----EY-----------DPTIE--DS 39
KRAS     P01116 -----------------------------------------------------------MTEYKLVVVGAGGVGKSALTIQLIQN--HF--V-D----EY-----------DPTIE--DS 39
MRAS     O14807 -------------------------------------------------MATSAVPSDNLPTYKLVVVGDGGVGKSALTIQFFQK--IF--V-P----DY-----------DPTIE--DS 49
NKIRAS1  Q9NYS0 ----------------------------------------------------------MGKGCKVVVCGLLSVGKTAILEQLLYG--NHTIG-M----ED-----------CETME--DV 42
NKIRAS2  Q9NYR9 ----------------------------------------------------------MGKSCKVVVCGQASVGKTSILEQLLYG--NHVVG-S----EM-----------IETQE--DI 42
NRAS     P01111 -----------------------------------------------------------MTEYKLVVVGAGGVGKSALTIQLIQN--HF--V-D----EY-----------DPTIE--DS 39
RALA     P11233 ------------------------------------------------MAANKPKGQNSLALHKVIMVGSGGVGKSALTLQFMYD--EF--V-E----DY-----------EPTKA--DS 50
RALB     P11234 ------------------------------------------------MAANKSKGQSSLALHKVIMVGSGGVGKSALTLQFMYD--EF--V-E----DY-----------EPTKA--DS 50
RAP1A    P62834 -----------------------------------------------------------MREYKLVVLGSGGVGKSALTVQFVQG--IF--V-E----KY-----------DPTIE--DS 39
RAP1B    P61224 -----------------------------------------------------------MREYKLVVLGSGGVGKSALTVQFVQG--IF--V-E----KY-----------DPTIE--DS 39
RAP2A    P10114 -----------------------------------------------------------MREYKVVVLGSGGVGKSALTVQFVTG--TF--I-E----KY-----------DPTIE--DF 39
RAP2B    P61225 -----------------------------------------------------------MREYKVVVLGSGGVGKSALTVQFVTG--SF--I-E----KY-----------DPTIE--DF 39
RAP2C    Q9Y3L5 -----------------------------------------------------------MREYKVVVLGSGGVGKSALTVQFVTG--TF--I-E----KY-----------DPTIE--DF 39
RASD1    Q9Y272 MKLAAM------------IK--------------------------KMCPSDSELSIPAKNCYRMVILGSSKVGKTAIVSRFLTG--RF--E-D----AY-----------TPTIE--DF 60
RASD2    Q96D21 -----M------------MK--------------------------TLSSGNCTLSVPAKNSYRMVVLGASRVGKSSIVSRFLNG--RF--E-D----QY-----------TPTIE--DF 55
RASL10A  Q92737 ----------------------------------------------------------MGGSLRVAVLGAPGVGKTAIIRQFLFG--DY--P-E----RH-----------RPTDG--PR 40
RASL10B  Q96S79 ----------------------------------------------------------MVSTYRVAVLGARGVGKSAIVRQFLYN--EF--S-E----VC-----------VPTTA--RR 40
RASL11A  Q6T310 ---LSM-----------SGHFL-----LAPI--------PE-----------SSSDYLLPKDIKLAVLGAGRVGKSAMIVRFLTK--RF--I-G----DY-----------EPNTG--KL 63
RASL11B  Q9BPW5 ---IQN-----------MCTIA-----EYPA--------PG-----NAAASDCCVGAAGRRLVKIAVVGASGVGKTALVVRFLTK--RF--I-G----DY-----------ERNAG--NL 69
RasL12   Q9NYN1 -----------------MSSVF-----GKPR--------AG------------SGPQSAPLEVNLAILGRRGAGKSALTVKFLTK--RF--I-S----EY-----------DPNLE--DT 56
RERG     Q96A58 --------------------------------------------------------MAKSAEVKLAIFGRAGVGKSALVVRFLTK--RF--I-W----EY-----------DPTLE--ST 42
RERGL    G5EA41 -----------------------------------------------------------MNDVKLAVLGGEGTGKSALTVRFLTK--RF--I-G----EY-----------ASNFE--SI 39
RHEB     Q15382 --------------------------------------------------------MPQSKSRKIAILGYRSVGKSSLTIQFVEG--QF--V-D----SY-----------DPTIE--NT 42
RHEBL1   Q8TAI7 --------------------------------------------------------MPLVRYRKVVILGYRCVGKTSLAHQFVEG--EF--S-E----GY-----------DPTVE--NT 42
RIT1     Q92963 -----M-----------DSG-------------------TR-----PVGSC-CSSPAGLSREYKLVMLGAGGVGKSAMTMQFISH--RF--P-E----DH-----------DPTIE--DA 57
RIT2     Q99578 -----M-----------E---------------------VE-----NEASCSPGSASGGSREYKVVMLGAGGVGKSAMTMQFISH--QF--P-D----YH-----------DPTIE--DA 56
RRAS     P10301 -----M-----------SSGAAS----GTGR--------GR-----PRGGGPGPGDPPPSETHKLVVVGGGGVGKSALTIQFIQS--YF--V-S----DY-----------DPTIE--DS 65
```

```
RRAS2   P62070 ------------------------------------------------MAAAGWRDGSGQEKYRLVVVGGGGVGKSALTIQFIQS--YF--V-T----DY-----------DPTIE--DS 50
GEM     P55040 KEPHQYSHRNRHSATPEDHCRRSWSSDST-----------------DSVIS----SESGNTYYRVVLIGEQGVGKSTLANIFAGV--HDSMD-S----DC-----------EVLGE--DT 113
REM1    O75628 PRLGQSASLN-PPTQKPSPAPDDWSSESS-----------------DS-------EGSWEALYRVVLLGDPGVGKTSLASLFAGK--QERDL-------H-----------EQLGE--DV 116
REM2    Q8IYK8 -----L-------RRAQAVDELDWPPQASSS--------GS----SDSLGSGEAAPAQKDGIFKVMLVGESGVGKSTLAGTFGGL--QGDSA-H----E------------PENPE--DT 151
RRAD    P55042 LTPGALTAAA--AGTGTQGPRLDWPEDSE-----------------DSLSSGG--SDSDESVYKVLLLGAPGVGKSALARIFGGV--EDGPE-A----E--------------AAG--HT 126
CDC42   P60953 -----------------------------------------------------------MQTIKCVVVGDGAVGKTCLLISYTTN--KF--P-S----EY-----------VPTVF--DN 39
RAC1    P63000 -----------------------------------------------------------MQAIKCVVVGDGAVGKTCLLISYTTN--AF--P-G----EY-----------IPTVF--DN 39
RAC2    P15153 -----------------------------------------------------------MQAIKCVVVGDGAVGKTCLLISYTTN--AF--P-G----EY-----------IPTVF--DN 39
RAC3    P60763 -----------------------------------------------------------MQAIKCVVVGDGAVGKTCLLISYTTN--AF--P-G----EY-----------IPTVF--DN 39
RHOA    P61586 ---------------------------------------------------------MAAIRKKLVIVGDGACGKTCLLIVFSKD--QF--P-E----VY-----------VPTVF--EN 41
RHOB    P62745 ---------------------------------------------------------MAAIRKKLVVVGDGACGKTCLLIVFSKD--EF--P-E----VY-----------VPTVF--EN 41
RHOBTB1 O94844 ------------------------------------------------MDADMDYERPNVETIKCVVVGDNAVGKTRLICARACN--TT--L-T----QY-----QLLATHVPTVWAIDQ 58
RHOBTB2 Q9BYZ6 ------------------------------------------------MDSDMDYERPNVETIKCVVVGDNAVGKTRLICARACN--AT--L-T----QY-----QLLATHVPTVWAIDQ 58
RHOC    P08134 ---------------------------------------------------------MAAIRKKLVIVGDGACGKTCLLIVFSKD--QF--P-E----VY-----------VPTVF--EN 41
RHOD    O00212 ---------------------------------------------MTAAQAAGEEAPPGVRSVKVVLVGDGGCGKTSLLMVFADG--AF--P-E----SY-----------TPTVF--ER 53
RHOF    Q9HBH0 ------------------------------------M--DA----PGA-LAQTAAPGPGRKELKIVIVGDGGCGKTSLLMVYSQG--SF--P-E----HY-----------APSVF--EK 55
RHOG    P84095 -----------------------------------------------------------MQSIKCVVVGDGAVGKTCLLICYTTN--AF--P-K----EY-----------IPTVF--DN 39
RHOH    Q15669 ----------------------------------------------------------MLSSIKCVLVGDSAVGKTSLLVRFTSE--TF--P-E----AY-----------KPTVY--EN 40
RHOJ    Q9H4E5 -----------------------------------MN--CK----EGTDSSCGCRGNDEKKMLKCVVVGDGAVGKTCLLMSYAND--AF--P-E----EY-----------VPTVF--DH 57
RHOQ    P17081 -----------------------------------------------------MAHGPGALMLKCVVVGDGAVGKTCLLMSYAND--AF--P-E----EY-----------VPTVF--DH 45
RHOU    Q7L0Q8 ---RGG-------------R---------------GG--RG----PGEPGGRGRAGGAEGRGVKCVLVGDGAVGKTSLVVSYTTN--GY--P-T----EY-----------IPTAF--DN 85
RHOV    Q96L33 -----------------------------------------------------RRSAPPELGIKCVLVGDGAVGKSSLIVSYTCN--GY--P-A----RY-----------RPTAL--DT 67
RND1    Q92730 -----M-------------K---------------ER--RA----PQ----------PVVARCKLVLVGDVQCGKTAMLQVLAKD--CY--P-E----TY-----------VPTVF--EN 49
RND2    P52198 -------------------------------------------------------MEGQSGRCKIVVVGDAECGKTALLQVFAKD--AY--P-G----SY-----------VPTVF--EN 43
RND3    P61587 -----M-------------K---------------ER--RA----SQKLSSKSIMDPNQNVKCKIVVVGDSQCGKTALLHVFAKD--CF--P-E----NY-----------VPTVF--EN 59
RAN     P62826 ----------------------------------------------------MAAQGEPQVQFKLVLVGDGGTGKTTFVKRHLTG--EF--E-K----KY-----------VATLG--VE 46
RHOT1   Q8IXI2 ----------------------------------------------------------MKKDVRILLVGEPRVGKTSLIMSLVSE--EF--P-E----EV-----------PPRAE---E 39
RHOT2   Q8IXI1 ----------------------------------------------------------MRRDVRILLLGEAQVGKTSLILSLVGE--EF--P-E----EV-----------PPRAE---E 39
IFT22   Q9H7X7 -----------------------------------------------------------MLKAKILFVGPCESGKTVLANFLTES--SD--I-T----EY-----------SPTQG--VR 39
RABL2A  Q9UBK7 ------------------------------------MA-ED----KTKPSELDQGKYDADDNVKIICLGDSAVGKSKLMERFLMD--GF--Q-P----QQ-----------LSTYA--LT 57
RABL2B  Q9UNT1 ------------------------------------MA-ED----KTKPSELDQGKYDADDNVKIICLGDSAVGKSKLMERFLMD--GF--Q-P----QQ-----------LSTYA--LT 57
RABL3   Q5HYI8 --------------------------------------------------------MASLDRVKVLVLGDSGVGKSSLVHLLCQN--QVLGN-P----SW-----------TVGCS--VD 44
RABL6   Q3YEC7 ----SM----------------------------------------N-QALQRRFAKGVQYNMKIVIRGDRNTGKTALWHRLQGR--PF--V-E----EY-----------IPTQE--IQ 79
RRAGA   Q7L523 -------------------------------------------------------MPNTAMKKKVLLMGKSGSGKTSMRSIIFAN--YI--A------RDTRRLGAT------------- 42
RRAGB   Q5VZM2 ---------------SEKTTEKENLGPRM----DPPLGEPEGS--------LGWVLPNTAMKKKVLLMGKSGSGKTSMRSIIFAN--YI--A------RDTRRLGATILDRIHSLQINSS 88
RRAGC   Q9HB90 ---------------EEAAAAGGGVGAG-----------AGGGCG---------PGGADSSKPRILLMGLRRSGKSSIQKVVFHK--MS--P------N------ET-LFLESTNK---- 98
RRAGD   Q9NQL2 ---------------EDELVGLADYGDGP----DSSDADPDSGTEEGV--LDFSDPFSTEVKPRILLMGLRRSGKSSIQKVVFHK--MS--P------N------ET-LFLESTNK---- 99
```

```
ARF1    P84077 ---------------FNVETV----E-----Y---------------KNISFTVWDVGGQDKIR----------------------PLWRHYFQNTQGLIFVVDSNDR-ERVNEAREELM 108
ARF3    P61204 ---------------FNVETV----E-----Y---------------KNISFTVWDVGGQDKIR----------------------PLWRHYFQNTQGLIFVVDSNDR-ERVNEAREELM 108
ARF4    P18085 ---------------FNVETV----E-----Y---------------KNICFTVWDVGGQDRIR----------------------PLWKHYFQNTQGLIFVVDSNDR-ERIQEVADELQ 108
ARF5    P84085 ---------------FNVETV----E-----Y---------------KNICFTVWDVGGQDKIR----------------------PLWRHYFQNTQGLIFVVDSNDR-ERVQESADELQ 108
ARF6    P62330 ---------------FNVETV----T-----Y---------------KNVKFNVWDVGGQDKIR----------------------PLWRHYYTGTQGLIFVVDCADR-DRIDEARQELH 104
ARL1    P40616 ---------------FNVETV----T-----Y---------------KNLKFQVWDLGGQTSIR----------------------PYWRCYYSNTDAVIYVVDSCDR-DRIGISKSELV 108
ARL2    P36404 ---------------FNIKTL----E-----H---------------RGFKLNIWDVGGQKSLR----------------------SYWRNYFESTDGLIWVVDSADR-QRMQDCQRELQ 107
ARL3    P36405 ---------------FNIKSV----Q-----S---------------QGFKLNVWDIGGQRKIR----------------------PYWKNYFENTDILIYVIDSADR-KRFEETGQELA 108
ARL4A   P40617 ---------------FNTEKI----K-----VTL----------GNSKTVTFHFWDVGGQEKLR----------------------PLWKSYTRCTDGIVFVVDSVDV-ERMEEAKTELH 116
ARL4C   P56559 ---------------FNTEKI----K-----LSN----------GTAKGISCHFWDVGGQEKLR----------------------PLWKSYSRCTDGIIYVVDSVDV-DRLEEAKTELH 109
ARL4D   P49703 ---------------FNTEKI----R-----VPL----------GGSRGITFQVWDVGGQEKLR----------------------PLWRSYTRRTDGLVFVVDAAEA-ERLEEAKVELH 117
ARL5A   Q9Y689 ---------------SNVEEI----V-----I---------------NNTRFLMWDIGGQESLR----------------------SSWNTYYTNTEFVIVVVDSTDR-ERISVTREELY 107
ARL5B   Q96KC2 ---------------SNVEEI----V-----V---------------KNTHFLMWDIGGQESLR----------------------SSWNTYYSNTEFIILVVDSIDR-ERLAITKEELY 107
ARL5C   A6NH57 ---------------SNVEEI----I-----L---------------PKTHFFMWDIVRPEALS----------------------FIWNTYYSNTEFIILVIDSTDR-DRLLTTREELY 107
ARL6    Q9H0F7 ---------------FSIEKF----K-----S---------------SSLSFTVFDMSGQGRYR----------------------NLWEHYYKEGQAIIFVIDSSDR-LRMVVAKEELD 110
ARL8A   Q96BM9 ---------------FNMRKI----T-----K---------------GNVTIKLWDIGGQPRFR----------------------SMWERYCRGVSAIVYMVDAADQ-EKIEASKNELH 112
ARL8B   Q9NVJ2 ---------------FNMRKV----T-----K---------------GNVTIKIWDIGGQPRFR----------------------SMWERYCRGVNAIVYMIDAADR-EKIEASRNELH 112
ARL9    Q6T311 ---------------FHAVCI----N-----T---------------EDSQMEFLEIGGSKPFR----------------------SYWEMYLSKGLLLIFVVDSADH-SRLPEAKKYLH 110
ARL10   Q8N8L6 ---------------FNSVRL----P-----T---------------KDFEVDLLEIGGSQNLR----------------------FYWKEFVSEVDVLVFVVDSADR-LRLPWARQELH 169
ARL11   Q969Q4 ---------------FNVEPL----K-----AP--------------GHVSLTLWDVGGQAPLR----------------------ASWKDYLEGTDILVYVLDSTDE-ARLPESAAELT 104
ARL13A  Q5H913 ---------------SELTTL----L-----L---------------DEYELSIYDLNGDLKGR----------------------EAWPNYYAQAHGLVFVLDSSDI-RRMQEVKIILT 112
ARL13B  Q3SXY8 ---------------FSKINL----R-----Q---------------GKFEVTIFDLGGGIRIR----------------------GIWKNYYAESYGVIFVVDSSDE-ERMEETKEAMS 112
ARL14   Q8N4G2 ---------------FNVEMI----E-----LE--------------RNLSLTVWDVGGQEKMR----------------------TVWGCYCENTDGLVYVVDSTDK-QRLEESQRQFE 105
ARL15   Q9NXU5 ---------------FSIKAV----P-----F---------------QNAILNVKELGGADNIR----------------------KYWSRYYQGSQGVIFVLDSASSEDDLEAARNELH 124
ARL16   Q0P5N6 ---------------TNLTDI----V-----A----------------QRKITIRELGGC--MG----------------------PIWSSYYGNCRSLLFVMDASDP-TQLSASCVQLL 121
ARL17   Q8IVW1 ---------------FCVETV----E-----Y---------------KNNTFAVWDVGSHFKIR----------------------PLWQHFFQNTKGARSPG-STHQ-GSLASGVLPIK 107
ARFRP1  Q13795 ---------------LNIGTV----D-----V---------------GKARLMFWDLGGQEELQ----------------------SLWDKYYAECHGVIYVIDSTDE-ERLAESKQAFE 116
SAR1A   Q9NR31 ---------------PTSEEL----T-----I---------------AGMTFTTFDLGGHEQAR----------------------RVWKNYLPAINGIVFLVDCADH-SRLVESKVELN 116
SAR1B   Q9Y6B6 ---------------PTSEEL----T-----I---------------AGMTFTTFDLGGHVQAR----------------------RVWKNYLPAINGIVFLVDCADH-ERLLESKEELD 116
TRIM23  P36406 ---------------FNVETV----E-----Y---------------KNLKFTIWDVGGKHKLR----------------------PLWKHYYLNTQAVVFVVDSSHR-DRISEAHSELA 495
IFT27   Q9BW83 ---------------LVVKTV----P--------V--------PDTGDSVELFIFDSAGKELFS----------------------EMLDKLWESPNVLCLVYDVTNE-ESFNNCS-KWL 104
RAB1A   P62820 ---------------FKIRTI----E--------L----------DGKTIKLQIWDTAGQERFR----------------------TITSSYYRGAHGIIVVYDVTDQ-ESFNNVK-QWL 106
RAB1B   Q9H0U4 ---------------FKIRTI----E--------L----------DGKTIKLQIWDTAGQERFR----------------------TITSSYYRGAHGIIVVYDVTDQ-ESYANVK-QWL 103
RAB1C   Q92928 ---------------FKIQTI----E--------L----------DGKTIKLQIWDTAGQERFW----------------------TITSSYYRGAHGFLVVYDVTDQ-ESYANVK-QWL 103
RAB2A   P61019 ---------------FGARMI----T--------I----------DGKQIKLQIWDTAGQESFR----------------------SITRSYYRGAAGALLVYDITRR-DTFNHLT-TWL 101
RAB2B   Q8WUD1 ---------------FGARMV----N--------I----------DGKQIKLQIWDTAGQESFR----------------------SITRSYYRGAAGALLVYDITRR-ETFNHLT-SWL 101
RAB3A   P20336 ---------------FKVKTI----Y--------R----------NDKRIKLQIWDTAGQERYR----------------------TITTAYYRGAMGFILMYDITNE-ESFNAVQ-DWS 117
RAB3B   P20337 ---------------FKVKTV----Y--------R----------HEKRVKLQIWDTAGQERYR----------------------TITTAYYRGAMGFILMYDITNE-ESFNAVQ-DWA 117
RAB3C   Q96E17 ---------------FKVKTV----F--------K----------NEKRIKLQIWDTAGQERYR----------------------TITTAYYRGAMGFILMYDITNE-ESFNAVQ-DWS 125
RAB3D   O95716 ---------------FKVKTV----Y--------R----------HDKRIKLQIWDTAGQERYR----------------------TITTAYYRGAMGFLLMYDIANQ-ESFAAVQ-DWA 117
RAB4A   P20338 ---------------FGSKII----N--------V----------GGKYVKLQIWDTAGQERFR----------------------SVTRSYYRGAAGALLVYDITSR-ETYNALT-NWL 108
RAB4B   P61018 ---------------FGSRVV----N--------V----------GGKTVKLQIWDTAGQERFR----------------------SVTRSYYRGAAGALLVYDITSR-ETYNSLA-AWL 103
RAB5A   P20339 ---------------FLTQTV----C--------L----------DDTTVKFEIWDTAGQERYH----------------------SLAPMYYRGAQAAIVVYDITNE-ESFARAK-NWV 115
RAB5B   P61020 ---------------FLTQSV----C--------L----------DDTTVKFEIWDTAGQERYH----------------------SLAPMYYRGAQAAIVVYDITNQ-ETFARAK-TWV 115
RAB5C   P51148 ---------------FLTQTV----C--------L----------DDTTVKFEIWDTAGQERYH----------------------SLAPMYYRGAQAAIVVYDITNT-DTFARAK-NWV 116
RAB6A   P20340 ---------------FLSKTM----Y--------L----------EDRTVRLQLWDTAGQERFR----------------------SLIPSYIRDSTVAVVVYDITNV-NSFQQTT-KWI 108
RAB6B   Q9NRW1 ---------------FLSKTM----Y--------L----------EDRTVRLQLWDTAGQERFR----------------------SLIPSYIRDSTVAVVVYDITNL-NSFQQTS-KWI 108
RAB6C   Q9H0N0 ---------------FLSKTM----Y--------L----------EDGTIGLRLWDTAGQERLR----------------------SLIPRYIRDSAAAVVVYDITNV-NSFQQTT-KWI 108
RAB7A   P51149 ---------------FLTKEV----M--------V----------DDRLVTMQIWDTAGQERFQ----------------------SLGVAFYRGADCCVLVFDVTAP-NTFKTLD-SWR 103
RAB7B   Q96AH8 ---------------ILSKII----I--------L----------GDTTLKLQIWDTGGQERFR----------------------SMVSTFYKGSDGCILAFDVTDL-ESFEALD-IWR 103
RAB8A   P61006 ---------------FKIRTI----E--------L----------DGKRIKLQIWDTAGQERFR----------------------TITTAYYRGAMGIMLVYDITNE-KSFDNIR-NWI 103
RAB8B   Q92930 ---------------FKIRTI----E--------L----------DGKKIKLQIWDTAGQERFR----------------------TITTAYYRGAMGIMLVYDITNE-KSFDNIK-NWI 103
RAB9A   P51151 ---------------FLNKDL----E--------V----------DGHFVTMQIWDTAGQERFR----------------------SLRTPFYRGSDCCLLTFSVDDS-QSFQNLS-NWK 102
RAB9B   Q9NP90 ---------------FLNRDL----E--------V----------DGRFVTLQIWDTAGQERFK----------------------SLRTPFYRGADCCLLTFSVDDR-QSFENLG-NWQ 102
RAB10   P61026 ---------------FKIKTV----E--------L----------QGKKIKLQIWDTAGQERFH----------------------TITTSYYRGAMGIMLVYDITNG-KSFENIS-KWL 104
RAB11A  P62491 ---------------FATRSI----Q--------V----------DGKTIKAQIWDTAGQERYR----------------------AITSAYYRGAVGALLVYDIAKH-LTYENVE-RWL 106
RAB11B  Q15907 ---------------FATRSI----Q--------V----------DGKTIKAQIWDTAGQERYR----------------------AITSAYYRGAVGALLVYDIAKH-LTYENVE-RWL 106
RAB12   Q6IQ22 ---------------FKIKTV----E--------L----------RGKKIRLQIWDTAGQERFN----------------------SITSAYYRSAKGIILVYDITKK-ETFDDLP-KWM 137
RAB13   P51153 ---------------FKIRTV----D--------I----------EGKKIKLQVWDTAGQERFK----------------------TITTAYYRGAMGIILVYDITDE-KSFENIQ-NWM 103
RAB14   P61106 ---------------FGTRII----E--------V----------SGQKIKLQIWDTAGQERFR----------------------AVTRSYYRGAAGALMVYDITRR-STYNHLS-SWL 106
RAB15   P59190 ---------------FKMKTI----E--------V----------DGIKVRIQIWDTAGQERYQ----------------------TITKQYYRRAQGIFLVYDISSE-RSYQHIM-KWV 103
RAB17   Q9H0T7 ---------------FFTKVV----D--------V----------GATSLKLEIWDTAGQEKYH----------------------SVCHLYFRGANAALLVYDITRK-DSFLKAQ-QWL 113
RAB18   Q9NP72 ---------------FKVKTI----S--------V----------DGNKAKLAIWDTAGQERFR----------------------TLTPSYYRGAQGVILVYDVTRR-DTFVKLD-NWL 103
RAB19   A4D1S5 ---------------FTVRSL----D--------I----------DGKKVKMQVWDTAGQERFR----------------------TITQSYYRSAHAAIIAYDLTRR-STFESIP-HWI 112
RAB20   Q9NX57 ---------------FYLKQW----R--------S--------------YNISIWDTAGREQFH----------------------GLGSMYCRGAAAIILTYDVNHR-QSLVELE-DRF 95
RAB21   Q9UL25 ---------------FLTKKL----N--------I----------GGKRVNLAIWDTAGQERFH----------------------ALGPIYYRDSNGAILVYDITDE-DSFQKVK-NWV 114
RAB22A  Q9UL26 ---------------FMTKTV----Q--------Y----------QNELHKFLIWDTAGQERFR----------------------ALAPMYYRGSAAAIIVYDITKE-ETFSTLK-NWV 100
RAB23   Q9ULC3 ---------------FLERQI----Q--------V----------NDEDVRLMLWDTAGQEEFD----------------------AITKAYYRGAQACVLVFSTTDR-ESFEAVS-SWR 104
RAB24   Q969Q5 ---------------FVAKVM----S--------V----------GDRTVTLGIWDTAGSERYE----------------------AMSRIYYRGAKAAIVCYDLTDS-SSFERAK-FWV 103
RAB25   P57735 ---------------FSTRTV----M--------L----------GTAAVKAQIWDTAGLERYR----------------------AITSAYYRGAVGALLVFDLTKH-QTYAVVE-RWL 107
RAB26   Q9ULW5 ---------------FRNKVL----D--------V----------DGVKVKLQMWDTAGQERFR----------------------SVTHAYYRDAHALLLLYDVTNK-ASFDNIQ-AWL 159
RAB27A  P51159 ---------------FREKRV----V--------YRASGPDGATGRGQRIHLQLWDTAGQERFR----------------------SLTTAFFRDAMGFLLLFDLTNE-QSFLNVR-NWI 114
RAB27B  O00194 ---------------FREKRV----V--------YNAQGPNGSSGKAFKVHLQLWDTAGQERFR----------------------SLTTAFFRDAMGFLLMFDLTSQ-QSFLNVR-NWM 114
RAB28   P51157 ---------------FFLRR----IT--------L---------PGNLNVTLQIWDIGGQTIGG----------------------KMLDKYIYGAQGVLLVYDITNY-QSFENLE-DWY 108
RAB29   O14966 ---------------FALKVL----Q--------W---------SDYEIVRLQLWDIAGQERFT----------------------SMTRLYYRDASACVIMFDVTNA-TTFSNSQ-RWK 103
RAB30   Q15771 ---------------FMIKTV----E--------I----------NGEKVKLQIWDTAGQERFR----------------------SITQSYYRSANALILTYDITCE-ESFRCLP-EWL 104
RAB31   Q13636 ---------------FMTKTV----P--------C----------GNELHKFLIWDTAGQERFH----------------------SLAPMYYRGSAAAVIVYDITKQ-DSFYTLK-KWV 100
RAB32   Q13637 ---------------FALKVL----N--------W---------DSRTLVRLQLWDIAGQERFG----------------------NMTRVYYKEAVGAFVVFDISRS-STFEAVL-KWK 121
RAB33A  Q14088 ---------------FREKTV----E--------I----------EGEKIKVQVWDTAGQERFRK---------------------SMVEHYYRNVHAVVFVYDVTKM-TSFTNLK-MWI 132
RAB33B  Q9H082 ---------------FRERAV----E--------I----------DGERIKIQLWDTAGQERFRK---------------------SMVQHYYRNVHAVVFVYDMTNM-ASFHSLP-SWI 129
RAB34   Q9BZG1 ---------------FEMERF----E--------V----------LGIPFSLQLWDTAGQERFK----------------------CIASTYYRGAQAIIIVFNLNDV-ASLEHTK-QWL 147
RAB35   Q15286 ---------------FKIRTV----E--------I----------NGEKVKLQIWDTAGQERFR----------------------TITSTYYRGTHGVIVVYDVTSA-ESFVNVK-RWL 103
RAB36   O95755 ---------------FEIERF----E--------I----------AGIPYSLQIWDTAGQEKFK----------------------CIASAYYRGAQVIITAFDLTDV-QTLEHTR-QWL 218
RAB37   Q96AX2 ---------------FRNKVV----T--------V----------DGVRVKLQIWDTAGQERFR----------------------SVTHAYYRDAQALLLLYDITNK-SSFDNIR-AWL 125
RAB38   P57729 ---------------FALKVL----H--------W---------DPETVVRLQLWDIAGQERFG----------------------NMTRVYYREAMGAFIVFDVTRP-ATFEAVA-KWK 105
```

```
RAB39A   Q14964 ---------------FFSRLL----E--------I---------EPGKRIKLQLWDTAGQERFR---------------------SITRSYYRNSVGGFLVFDITNR-RSFEHVK-DWL 108
RAB39B   Q96DA2 ---------------FFSRLV----E--------I---------EPGKRIKLQIWDTAGQERFR---------------------SITRAYYRNSVGGLLLFDITNR-RSFQNVH-EWL 104
RAB40A   Q8WXH6 ---------------YKTTTI----L--------L----------DGQRVKLKLWDTSGQGRFC---------------------TIFRSYSRGAQGVILVYDIANR-WSFEGMD-RWI 109
RAB40B   Q12829 ---------------YKTTTI----L--------L----------DGRRVKLQLWDTSGQGRFC---------------------TIFRSYSRGAQGVILVYDIANR-WSFDGID-RWI 109
RAB40C   Q96S21 ---------------YKTTTI----L--------L----------DGRRVKLELWDTSGQGRFC---------------------TIFRSYSRGAQGILLVYDITNR-WSFDGID-RWI 109
RAB41    Q5JT25 ---------------FLSKTM----Y--------L----------EDQIVQLQLWDTAGQERFH---------------------SLIPSYIRDSTIAVVVYDITNI-NSFKETD-KWV 126
RAB42    Q8N4Z0 ---------------CYRRAL----Q--------L---------RAGPRVKLQLWDTAGHERFR---------------------CITRSFYRNVVGVLLVFDVTNR-KSFEHIQ-DWH 110
RAB43    Q86YS6 ---------------FTMKTL----E--------I----------QGKRVKLQIWDTAGQERFR---------------------TITQSYYRSANGAILAYDITKR-SSFLSVP-HWI 113
RAB44    Q7Z6P3 ---------------FRVKTL----L--------V----------DNKCFVLQLWDTAGQERYH---------------------SMTRQLLRKADGVVLMYDITSQ-ESFAHVR-YWL 928
RASEF    Q8IZ41 ---------------FQMKTL----I--------V----------DGERTVLQLWDTAGQERFR---------------------SIAKSYFRKADGVLLLYDVTCE-KSFLNIR-EWV 636
DIRAS1   O95057 ----------------YRQV----IS--------C----------DKSVCTLQITDTTGSHQFP---------------------AMQRLSISKGHAFILVFSVTSK-QSLEELG-PIY 101
DIRAS2   Q96HU8 ----------------YRQV----IS--------C----------DKSICTLQITDTTGSHQFP---------------------AMQRLSISKGHAFILVYSITSR-QSLEELK-PIY 101
DIRAS3   O95661 ----------------YCQL----LG--------C----------SHGVLSLHITDSKSGDGNR---------------------ALQRHVIARGHAFVLVYSVTKK-ETLEELK-AFY 131
ERAS     Q7Z444 ----------------YWKE----LT--------L----------DSGDCILNVLDTAGQAIHR---------------------ALRDQCLAVCDGVLGVFALDDP-SSLIQLQ-Q-- 133
HRAS     P01112 ----------------YRKQ----VV--------I----------DGETCLLDILDTAGQEEYS---------------------AMRDQYMRTGEGFLCVFAINNT-KSFEDIH-QYR 97
KRAS     P01116 ----------------YRKQ----VV--------I----------DGETCLLDILDTAGQEEYS---------------------AMRDQYMRTGEGFLCVFAINNT-KSFEDIH-HYR 97
MRAS     O14807 ----------------YLKH----TE--------I----------DNQWAILDVLDTAGQEEFS---------------------AMREQYMRTGDGFLIVYSVTDK-ASFEHVD-RFH 107
NKIRAS1  Q9NYS0 ----------------YMAS----VE--------T---------DRGVKEQLHLYDTRGSQEGV---------------------ELPKHYFSFADGFVLVYSVNNL-ESFQRVE-LLK 101
NKIRAS2  Q9NYR9 ----------------YVGS----IE--------T---------DRGVREQVRFYDTRGLRDGA---------------------ELPRHCFSCTDGYVLVYSTDSR-ESFQRVE-LLK 101
NRAS     P01111 ----------------YRKQ----VV--------I----------DGETCLLDILDTAGQEEYS---------------------AMRDQYMRTGEGFLCVFAINNS-KSFADIN-LYR 97
RALA     P11233 ----------------YRKK----VV--------L----------DGEEVQIDILDTAGQEDYA---------------------AIRDNYFRSGEGFLCVFSITEM-ESFAATA-DFR 108
RALB     P11234 ----------------YRKK----VV--------L----------DGEEVQIDILDTAGQEDYA---------------------AIRDNYFRSGEGFLLVFSITEH-ESFTATA-EFR 108
RAP1A    P62834 ----------------YRKQ----VE--------V----------DCQQCMLEILDTAGTEQFT---------------------AMRDLYMKNGQGFALVYSITAQ-STFNDLQ-DLR 97
RAP1B    P61224 ----------------YRKQ----VE--------V----------DAQQCMLEILDTAGTEQFT---------------------AMRDLYMKNGQGFALVYSITAQ-STFNDLQ-DLR 97
RAP2A    P10114 ----------------YRKE----IE--------V----------DSSPSVLEILDTAGTEQFA---------------------SMRDLYIKNGQGFILVYSLVNQ-QSFQDIK-PMR 97
RAP2B    P61225 ----------------YRKE----IE--------V----------DSSPSVLEILDTAGTEQFA---------------------SMRDLYIKNGQGFILVYSLVNQ-QSFQDIK-PMR 97
RAP2C    Q9Y3L5 ----------------YRKE----IE--------V----------DSSPSVLEILDTAGTEQFA---------------------SMRDLYIKNGQGFILVYSLVNQ-QSFQDIK-PMR 97
RASD1    Q9Y272 ----------------HRKF----YS--------I----------RGEVYQLDILDTSGNHPFP---------------------AMRRLSILTGDVFILVFSLDNR-DSFEEVQ-RLR 118
RASD2    Q96D21 ----------------HRKV----YN--------I----------RGDMYQLDILDTSGNHPFP---------------------AMRRLSILTGDVFILVFSLDNR-ESFDEVK-RLQ 113
RASL10A  Q92737 ---------------LYRPA----VL--------L----------DGAVYDLSIRDGDVAGPGSSPGG------P------EEWPDAKDWSLQDTDAFVLVYDICSP-DSFDYVK-ALR 108
RASL10B  Q96S79 ---------------LYLPA----VV--------M----------NGHVHDLQILDFPPIS-AFPVNT------L------QEWADTCCRGLRSVHAYILVYDICCF-DSFEYVK-TIR 107
RASL11A  Q6T310 ----------------YSRL----VY--------V----------EGDQLSLQIQDTPGGVQIQD---S-----------LPQVVDSLSKCVQWAEGFLLVYSITDY-DSYLSIR-PLY 128
RASL11B  Q9BPW5 ----------------YTRQ----VQ--------I----------EGETLALQVQDTPGIQVHEN---S------------LSCSEQLNRCIRWADAVVIVFSITDY-KSYELIS-QLH 133
RASL12   Q9NYN1 ----------------YSSE----ET--------V----------DHQPVHLRVMDTADLDTPR----------------------NCERYLNWAHAFLVVYSVDSR-QSFDSSS-SYL 113
RERG     Q96A58 ----------------YRHQ----AT--------I----------DDEVVSMEILDTAGQEDTI----------------------QREGHMRWGEGFVLVYDITDR-GSFEEVL-PLK 99
RERGL    G5EA41 ----------------YKKH----LC--------L----------ERKQLNLEIYDPCSQR---------------------------------------------------------- 62
RHEB     Q15382 ----------------FTKL----IT--------V----------NGQEYHLQLVDTAGQDEYS---------------------IFPQTYSIDINGYILVYSVTSI-KSFEVIK-VIH 100
RHEBL1   Q8TAI7 ----------------YSKI----VT--------L----------GKDEFHLHLVDTAGQDEYS---------------------ILPYSFIIGVHGYVLVYSVTSL-HSFQVIE-SLY 100
RIT1     Q92963 ----------------YKIR----IR--------I----------DDEPANLDILDTAGQAEFT---------------------AMRDQYMRAGEGFIICYSITDR-RSFHEVR-EFK 115
RIT2     Q99578 ----------------YKTQ----VR--------I----------DNEPAYLDILDTAGQAEFT---------------------AMREQYMRGGEGFIICYSVTDR-QSFQEAA-KFK 114
RRAS     P10301 ----------------YTKI----CS--------V----------DGIPARLDILDTAGQEEFG---------------------AMREQYMRAGHGFLLVFAINDR-QSFNEVG-KLF 123
RRAS2    P62070 ----------------YTKQ----CV--------I----------DDRAARLDILDTAGQEEFG---------------------AMREQYMRTGEGFLLVFSVTDR-GSFEEIY-KFQ 108
GEM      P55040 ----------------YERT----LM--------V----------DGESATIILLDMWENKG-EN---E----------------WLHDHCMQVGDAYLIVYSITDR-ASFEKAS-ELR 172
REM1     O75628 ----------------YERT----LT--------V----------DGEDTTLVVVDTWEAEKLDK---S----------------WSQESCLQGGSAYVIVYSIADR-GSFESAS-ELR 176
REM2     Q8IYK8 ----------------YERR----IM--------V----------DKEEVTLVVYDIWEQGD-AG---G----------------WLRDHCLQTGDAFLIVFSVTDR-RSFSKVP-ETL 210
RRAD     P55042 ----------------YDRS----IV--------V----------DGEEASLMVYDIWEQDG--G---R----------------WLPGHCMAMGDAYVIVYSVTDK-GSFEKAS-ELR 184
CDC42    P60953 ----------------YAVTV----M--------I----------GGEPYTLGLFDTAGQEDYD---------------------RLRPLSYPQTDVFLVCFSVVSP-SSFENVKEKWV 98
RAC1     P63000 ----------------YSANV----M--------V----------DGKPVNLGLWDTAGQEDYD---------------------RLRPLSYPQTDVFLICFSLVSP-ASFENVRAKWY 98
RAC2     P15153 ----------------YSANV----M--------V----------DSKPVNLGLWDTAGQEDYD---------------------RLRPLSYPQTDVFLICFSLVSP-ASYENVRAKWF 98
RAC3     P60763 ----------------YSANV----M--------V----------DGKPVNLGLWDTAGQEDYD---------------------RLRPLSYPQTDVFLICFSLVSP-ASFENVRAKWY 98
RHOA     P61586 ----------------YVADI----E--------V----------DGKQVELALWDTAGQEDYD---------------------RLRPLSYPDTDVILMCFSIDSP-DSLENIPEKWT 100
RHOB     P62745 ----------------YVADI----E--------V----------DGKQVELALWDTAGQEDYD---------------------RLRPLSYPDTDVILMCFSVDSP-DSLENIPEKWV 100
RHOBTB1  O94844 ----------------YRVCQ----EVLERSRDVV----------DEVSVSLRLWDTFGDHHKD-----------------------RRFAYGRSDVVVLCFSIANP-NSLNHVKSMWY 123
RHOBTB2  Q9BYZ6 ----------------YRVCQ----EVLERSRDVV----------DDVSVSLRLWDTFGDHHKD-----------------------RRFAYGRSDVVVLCFSIANP-NSLHHVKTMWY 123
RHOC     P08134 ----------------YIADI----E--------V----------DGKQVELALWDTAGQEDYD---------------------RLRPLSYPDTDVILMCFSIDSP-DSLENIPEKWT 100
RHOD     O00212 ----------------YMVNL----Q--------V----------KGKPVHLHIWDTAGQDDYD---------------------RLRPLFYPDASVLLLCFDVTSP-NSFDNIFNRWY 112
RHOF     Q9HBH0 ----------------YTASV----T--------V----------GSKEVTLNLYDTAGQEDYD---------------------RLRPLSYQNTHLVLICYDVMNP-TSYDNVLIKWF 114
RHOG     P84095 ----------------YSAQS----A--------V----------DGRTVNLNLWDTAGQEEYD---------------------RLRTLSYPQTNVFVICFSIASP-PSYENVRHKWH 98
RHOH     Q15669 ----------------TGVDV----F--------M----------DGIQISLGLWDTAGNDAFR---------------------SIRPLSYQQADVVLMCYSVANH-NSFLNLKNKWI 99
RHOJ     Q9H4E5 ----------------YAVTV----T--------V----------GGKQHLLGLYDTAGQEDYN---------------------QLRPLSYPNTDVFLICFSVVNP-ASYHNVQEEWV 116
RHOQ     P17081 ----------------YAVSV----T--------V----------GGKQYLLGLYDTAGQEDYD---------------------RLRPLSYPMTDVFLICFSVVNP-ASFQNVKEEWV 104
RHOU     Q7L0Q8 ----------------FSAVV----S--------V----------DGRPVRLQLCDTAGQDEFD---------------------KLRPLCYTNTDIFLLCFSVVSP-SSFQNVSEKWV 144
RHOV     Q96L33 ----------------FSVQV----L--------V----------DGAPVRIELWDTAGQEDFD---------------------RLRSLCYPDTDVFLACFSVVQP-SSFQNITEKWL 126
RND1     Q92730 ----------------YTACL----E--------T----------EEQRVELSLWDTSGSPYYD---------------------NVRPLCYSDSDAVLLCFDISRP-ETVDSALKKWR 108
RND2     P52198 ----------------YTASF----E--------I----------DKRRIELNMWDTSGSSYYD---------------------NVRPLAYPDSDAVLICFDISRP-ETLDSVLKKWQ 102
RND3     P61587 ----------------YTASF----E--------I----------DTQRIELSLWDTSGSPYYD---------------------NVRPLSYPDSDAVLICFDISRP-ETLDSVLKKWK 118
RAN      P62826 ---------------VHPLVF----H--------T----------NRGPIKFNVWDTAGQEKFG---------------------GLRDGYYIQAQCAIIMFDVTSR-VTYKNVP-NWH 105
RHOT1    Q8IXI2 ----------------ITIPA----D--------V----------TPERVPTHIVDYSEAEQSD---------------------EQLHQEISQANVICIVYAVNNK-HSIDKVTSRWI 98
RHOT2    Q8IXI1 ----------------ITIPA----D--------V----------TPEKVPTHIVDYSEAEQTD---------------------EELREEIHKANVVCVVYDVSEE-ATIEKIRTKWI 98
IFT22    Q9H7X7 ---------------ILEFENPHVTS--------N---------NKGTGCEFELWDCGGDAKFE---------------------SCWPALMKDAHGVVIVFNADIP-SHRKEME-MWY 103
RABL2A   Q9UBK7 ---------------LYKHTA----T--------V----------DGKTILVDFWDTAGQERFQ---------------------SMHASYYHKAHACIMVFDIQRK-VTYRNLS-TWY 116
RABL2B   Q9UNT1 ---------------LYKHTA----T--------V----------DGRTILVDFWDTAGQERFQ---------------------SMHASYYHKAHACIMVFDVQRK-VTYRNLS-TWY 116
RABL3    Q5HYI8 ---------------VRVHDY--KEG--------T---------PEEKTYYIELWDVGGSVGSAS-----------------SVKSTRAVFYNSVNGIIFVHDLTNK-KSSQNLR-RWS 110
RABL6    Q3YEC7 ---------------VTSIHW--SYK--------T----------TDDIVKVEVWDVVDKGKCKKRGDGLKMENDPQEAESEMALDAEFLDVYKNCNGVVMMFDITKQ-WTFNYIL-REL 162
RRAGA    Q7L523 ---------------IDVEHS----H-------V----------RFLGNLVLNLWDCGGQDTFME----------------NYFTSQRDNIFRNVEVLIYVFDVESR-E----LEKDMH 104
RRAGB    Q5VZM2 LSTYSLVDSVGNTKTFDVEHS----H-------V----------RFLGNLVLNLWDCGGQDTFME----------------NYFTSQRDNIFRNVEVLIYVFDVESR-E----LEKDMH 165
RRAGC    Q9HB90 ---------------IYKD-------------DI----------SNSSFVNFQIWDFPGQMDFFD----------------P--TFDYEMIFRGTGALIYVIDAQDD-YM--EALTRLH 158
RRAGD    Q9NQL2 ---------------ICRE-------------DV----------SNSSFVNFQIWDFPGQIDFFD----------------P--TFDYEMIFRGTGALIFVIDSQDD-YM--EALARLH 159

ARF1     P84077 R-------MLAEDEL-------------------RDAVL-LVFANKQDLPNAMNAAEI-------------------TDKLGLH------------SL-R---HRNW------------ 153
ARF3     P61204 R-------MLAEDEL-------------------RDAVL-LVFANKQDLPNAMNAAEI-------------------TDKLGLH------------SL-R---HRNW------------ 153
ARF4     P18085 K-------MLLVDEL-------------------RDAVL-LLFANKQDLPNAMAISEM-------------------TDKLGLQ------------SL-R---NRTW------------ 153
ARF5     P84085 K-------MLQEDEL-------------------RDAVL-LVFANKQDMPNAMPVSEL-------------------TDKLGLQ------------HL-R---SRTW------------ 153
ARF6     P62330 R-------IINDREM-------------------RDAII-LIFANKQDLPDAMKPHEI-------------------QEKLGLT------------RI-R---DRNW------------ 149
ARL1     P40616 A-------MLEEEEL-------------------RKAIL-VVFANKQDMEQAMTSSEM-------------------ANSLGLP------------AL-K---DRKW------------ 153
ARL2     P36404 S-------LLVEERL-------------------AGATL-LIFANKQDLPGALSSNAI-------------------REVLELD------------SI-R---SHHW------------ 152
ARL3     P36405 E-------LLEEEKL-------------------SCVPV-LIFANKQDLLTAAPASEI-------------------AEGLNLH------------TI-R---DRVW------------ 153
ARL4A    P40617 K-------ITRISEN-------------------QGVPV-LIVANKQDLRNSLSLSEI-------------------EKLLAMG------------EL-S--SSTPW------------ 162
ARL4C    P56559 K-------VTKFAEN-------------------QGTPL-LVIANKQDLPKSLPVAEI-------------------EKQLALH------------EL-I--PATTY------------ 155
ARL4D    P49703 R-------ISRASDN-------------------QGVPV-LVLANKQDQPGALSAAEV-------------------EKRLAVR------------EL-A--AATLT------------ 163
ARL5A    Q9Y689 K-------MLAHEDL-------------------RKAGL-LIFANKQDVKECMTVAEI-------------------SQFLKLT------------SI-K---DHQW------------ 152
ARL5B    Q96KC2 R-------MLAHEDL-------------------RKAAV-LIFANKQDMKGCMTAAEI-------------------SKYLTLS------------SI-K---DHPW------------ 152
ARL5C    A6NH57 K-------MLAHEAL-------------------QDASV-LIFANKQDVKDSMRMVEI-------------------SHFLTLS------------TI-K---DHSW------------ 152
ARL6     Q9H0F7 T-------LLNHPDI-----------------KHRRIPI-LFFANKMDLRDAVTSVKV-------------------SQLLCLE------------NI-K---DKPW------------ 157
ARL8A    Q96BM9 N-------LLDKPQL-------------------QGIPV-LVLGNKRDLPGALDEKEL-------------------IEKMNLS------------AI-Q---DREI------------ 157
ARL8B    Q9NVJ2 N-------LLDKPQL-------------------QGIPV-LVLGNKRDLPNALDEKQL-------------------IEKMNLS------------AI-Q---DREI------------ 157
ARL9     Q6T311 Q-------LIAANP---------------------VLPL-VVFANKQDLEAAYHITDI-------------------HEALALS------------EV-G--NDRKM------------ 154
ARL10    Q8N8L6 K-------LLDKDP---------------------DLPV-VVVANKQDLSEAMSMGEL-------------------QRELGLQ------------AI-D--NQREV------------ 213
ARL11    Q969Q4 E-------VLNDPNM-------------------AGVPF-LVLANKQEAPDALPLLKI-------------------RNRLSLE------------RF-Q---DHCW------------ 149
ARL13A   Q5H913 H-------LLSDKRV-------------------AGKPI-LILANKQDKKKALMPCDI-------------------IDYLLLK------------KL-VKENKCPC------------ 160
ARL13B   Q3SXY8 E-------MLRHPRI-------------------SGKPI-LVLANKQDKEGALGEADV-------------------IECLSLE------------KL-VNEHKCLC------------ 160
ARL14    Q8N4G2 H-------ILKNEHI-------------------KNVPV-VLLANKQDMPGALTAEDI-------------------TRMFKVK------------KL-C--SDRNW------------ 151
ARL15    Q9NXU5 S-------ALQHPQL-------------------CTLPF-LILANHQDKPAARSVQEI-------------------KKYFELE------------PL-A--RGKRW------------ 170
ARL16    Q0P5N6 G-------LLSAEQL-------------------AEASV-LILFNKIDLPCYMSTEEM-------------------KSLIRLP------------DI-IACAKQNI------------ 169
ARL17    Q8IVW1 C--SHVE-FGMWKGG-------------------RSHPF-LPHSSRCAG----SGGQL-------------------DSILP---------------H-Q---SPAW------------ 149
ARFRP1   Q13795 K-------VVTSEAL-------------------CGVPV-LVLANKQDVETCLSIPDI-------------------KTAFSDCTS----------KI-G---RRDC------------ 163
SAR1A    Q9NR31 A-------LMTDETI-------------------SNVPI-LILGNKIDRTDAISEEKL-------------------REIFGLYGQTTGKGNVTLKEL-N---ARPM------------ 173
SAR1B    Q9Y6B6 S-------LMTDETI-------------------ANVPI-LILGNKIDRPEAISEERL-------------------REMFGLYGQTTGKGSISLKEL-N---ARPL------------ 173
TRIM23   P36406 K-------LLTEKEL-------------------RDALL-LIFANKQDVAGALSVEEI-------------------TELLSLH------------KL-C--CGRSW------------ 541
IFT27    Q9BW83 E------KARSQAPG-------------------ISLPG-VLVGNKTDLAG------R-------------------RAVDSAEAR----------AW-----ALG-Q----------- 145
RAB1A    P62820 Q------EIDRYASE--------------------NVNK-LLVGNKCDLTT------K-------------------KVVDYTTAK----------EF-----ADS-L----------- 146
RAB1B    Q9H0U4 Q------EIDRYASE--------------------NVNK-LLVGNKSDLTT------K-------------------KVVDNTTAK----------EF-----ADS-L----------- 143
RAB1C    Q92928 Q------EIDRHASE--------------------NVNK-LLVGNKSDLTT------K-------------------KVVDNTTAK----------EF-----ADS-L----------- 143
RAB2A    P61019 E------DARQHSNS--------------------NMVI-MLIGNKSDLES------R-------------------REVKKEEGE----------AF-----ARE-H----------- 141
RAB2B    Q8WUD1 E------DARQHSSS--------------------NMVI-MLIGNKSDLES------R-------------------RDVKREEGE----------AF-----ARE-H----------- 141
RAB3A    P20336 T------QIKTYSWD--------------------NAQV-LLVGNKCDMED------E-------------------RVVSSERGR----------QL-----ADH-L----------- 157
RAB3B    P20337 T------QIKTYSWD--------------------NAQV-ILVGNKCDMEE------E-------------------RVVPTEKGQ----------LL-----AEQ-L----------- 157
RAB3C    Q96E17 T------QIKTYSWD--------------------NAQV-ILVGNKCDMED------E-------------------RVISTERGQ----------HL-----GEQ-L----------- 165
RAB3D    O95716 T------QIKTYSWD--------------------NAQV-ILVGNKCDLED------E-------------------RVVPAEDGR----------RL-----ADD-L----------- 157
RAB4A    P20338 T------DARMLASQ--------------------NIVI-ILCGNKKDLDA------D-------------------REVTFLEAS----------RF-----AQE-N----------- 148
RAB4B    P61018 T------DARTLASP--------------------NIVV-ILCGNKKDLDP------E-------------------REVTFLEAS----------RF-----AQE-N----------- 143
RAB5A    P20339 K------ELQRQASP--------------------NIVI-ALSGNKADLAN-------------------------KRAVDFQEAQ----------SY-----ADD-N----------- 155
RAB5B    P61020 K------ELQRQASP--------------------SIVI-ALAGNKADLAN-------------------------KRMVEYEEAQ----------AY-----ADD-N----------- 155
```

```
RAB5C   P51148 K------ELQRQASP--------------------NIVI-ALAGNKADLAS--------------------------KRAVEFQEAQ----------AY-----ADD-N----------- 156
RAB6A   P20340 D------DVRTERGS--------------------DVII-MLVGNKTDLAD--------------------------KRQVSIEEGE----------RK-----AKE-L----------- 148
RAB6A   Q9NRW1 D------DVRTERGS--------------------DVII-MLVGNKTDLAD--------------------------KRQITIEEGE----------QR-----AKE-L----------- 148
RAB6C   Q9H0N0 D------DVRTERGS--------------------DVII-TLVGNRTDLAD--------------------------KRQVSVEEGE----------RK-----AKG-L----------- 148
RAB7A   P51149 D------EFLIQASPR----------------DPENFPF-VVLGNKIDLEN---------------------------RQVATKRAQ----------AW-----CYS-K----------- 146
RAB7B   Q96AH8 G------DVLAKIVP-----------------MEQSYPM-VLLGNKIDLAD---------------------------RKVPQEVAQ----------GW-----CRE-K----------- 145
RAB8A   P61006 R------NIEEHASA--------------------DVEK-MILGNKCDVND------K--------------------RQVSKERGE----------KL-----ALD-Y----------- 143
RAB8B   Q92930 R------NIEEHASS--------------------DVER-MILGNKCDMND------K--------------------RQVSKERGE----------KL-----AID-Y----------- 143
RAB9A   P51151 K------EFIYYADVK----------------EPESFPF-VILGNKIDISE---------------------------RQVSTEEAQ----------AW-----CRD-N----------- 145
RAB9B   Q9NP90 K------EFIYYADVK----------------DPEHFPF-VVLGNKVDKED---------------------------RQVTTEEAQ----------TW-----CME-N----------- 145
RAB10   P61026 R------NIDEHANE--------------------DVER-MLLGNKCDMDD------K--------------------RVVPKGKGE----------QI-----ARE-H----------- 144
RAB11A  P62491 K------ELRDHADS--------------------NIVI-MLVGNKSDLRH------L--------------------RAVPTDEAR----------AF-----AEK-N----------- 146
RAB11B  Q15907 K------ELRDHADS--------------------NIVI-MLVGNKSDLRH------L--------------------RAVPTDEAR----------AF-----AEK-N----------- 146
RAB12   Q6IQ22 K------MIDKYASE--------------------DAEL-LLVGNKLDCET------D--------------------REITRQQGE----------KF-----AQQIT----------- 178
RAB13   P51153 K------SIKENASA--------------------GVER-LLLGNKCDMEA------K--------------------RKVQKEQAD----------KL-----ARE-H----------- 143
RAB14   P61106 T------DARNLTNP--------------------NTVI-ILIGNKADLEA------Q--------------------RDVTYEEAK----------QF-----AEE-N----------- 146
RAB15   P59190 S------DVDEYAPE--------------------GVQK-ILIGNKADEEQ------K--------------------RQVGREQGQ----------QL-----AKE-Y----------- 143
RAB17   Q9H0T7 K------DLEEELHP-------------------GEVLV-MLVGNKTDLSQ--------------------------EREVTFQEGK----------EF-----ADS-Q----------- 154
RAB18   Q9NP72 N------ELETYCTR-------------------NDIVN-MLVGNKIDKE-------N--------------------REVDRNEGL----------KF-----ARK-H----------- 143
RAB19   A4D1S5 H------EIEKYGAA--------------------NVVI-MLIGNKCDLWE------K--------------------RHVLFEDAC----------TL-----AEK-Y----------- 152
RAB20   Q9NX57 L------GLTDTASK--------------------DCLF-AIVGNKVDLTEEGALAGQEKEECSPNMDAGDRVSPRAPKQVQLEDAV----------AL-----YKKILKYKMLDEQDVP 173
RAB21   Q9UL25 K------ELRKMLGN--------------------EICL-CIVGNKIDLEK--------------------------ERHVSIQEAE----------SY-----AES-V----------- 154
RAB22A  Q9UL26 K------ELRQHGPP--------------------NIVV-AIAGNKCDLID--------------------------VREVMERDAK----------DY-----ADS-I----------- 140
RAB23   Q9ULC3 E------KVVAEVG---------------------DIPT-VLVQNKIDLLD------D--------------------SCIKNEEAE----------AL-----AKR-L----------- 143
RAB24   Q969Q5 K------ELRSLEEG---------------------CQI-YLCGTKSDLLEED----------------------RRRRRVDFHDVQ----------DY-----ADN-I----------- 146
RAB25   P57735 K------ELYDHAEA--------------------TIVV-MLVGNKSDLSQ------A--------------------REVPTEEAR----------MF-----AEN-N----------- 147
RAB26   Q9ULW5 T------EIHEYAQH--------------------DVAL-MLLGNKVDSAH------E--------------------RVVKREDGE----------KL-----AKE-Y----------- 199
RAB27A  P51159 S------QLQMHAYC-------------------ENPDI-VLCGNKSDLED------Q--------------------RVVKEEEAI----------AL-----AEK-Y----------- 155
RAB27B  O00194 S------QLQANAYC-------------------ENPDI-VLIGNKADLPD------Q--------------------REVNERQAR----------EL-----ADK-Y----------- 155
RAB28   P51157 T------VVK--KVSE----------------ESETQPLVALVGNKIDLE------HM--------------------RTIKPEKHL----------R-----FCQEN-----------G 152
RAB29   O14966 Q------DLDSKLTLP----------------NGEPVPC-LLANKCDLSPWA--------------------------V--SRDQID----------RF-----SKE-N----------- 146
RAB30   Q15771 R------EIEQYASN--------------------KVIT-VLVGNKIDLAE------R--------------------REVSQQRAE----------EF-----SEA-Q----------- 144
RAB31   Q13636 K------ELKEHGPE--------------------NIVM-AIAGNKCDLSD--------------------------IREVPLKDAK----------EY-----AES-I----------- 140
RAB32   Q13637 S------DLDSKVHLP----------------NGSPIPA-VLLANKCDQNKDS-------------------------SQ-SPSQVD----------QF-----CKE-H----------- 165
RAB33A  Q14088 Q------ECNGHAVP-------------------PLVPK-VLVGNKCDLRE------Q--------------------IQVPSNLAL----------KF-----ADA-H----------- 173
RAB33B  Q9H082 E------ECKQHLLA-------------------NDIPR-ILVGNKCDLRS------A--------------------IQVPTDLAQ----------KF-----ADT-H----------- 170
RAB34   Q9BZG1 A------DALKENDP-------------------SSVLL-FLVGSKKDLSTPA----Q--------------------YALMEKDAL----------QV-----AQE-M----------- 190
RAB35   Q15286 H------EINQNC-D--------------------DVCR-ILVGNKNDDPE------R--------------------KVVETEDAY----------KF-----AGQ-M----------- 142
RAB36   O95755 E------DALRENEA-------------------GSCFI-FLVGTKKDLLSGA----A--------------------CEQAEADAV----------HL-----ARE-M----------- 261
RAB37   Q96AX2 T------EIHEYAQR--------------------DVVI-MLLGNKADMSS------E--------------------RVIRSEDGE----------TL-----ARE-Y----------- 165
RAB38   P57729 N------DLDSKLSLP----------------NGKPVSV-VLLANKCDQGKDV-------------------------LMNNGLKMD----------QF-----CKE-H----------- 150
RAB39A  Q14964 E------EAKMYVQP-------------------FRIVF-LLVGHKCDLAS------Q--------------------RQVTREEAE----------KL-----SAD-C----------- 149
RAB39B  Q96DA2 E------ETKVHVQP-------------------YQIVF-VLVGHKCDLDT------Q--------------------RQVTRHEAE----------KL-----AAA-Y----------- 145
RAB40A  Q8WXH6 K------KIEEHAP---------------------GVPK-ILVGNRLHLAF------K--------------------RQVPREQAQ----------AY-----AER-L----------- 148
RAB40B  Q12829 K------EIDEHAP---------------------GVPK-ILVGNRLHLAF------K--------------------RQVPTEQAQ----------AY-----AER-L----------- 148
RAB40C  Q96S21 K------EIDEHAP---------------------GVPR-ILVGNRLHLAF------K--------------------RQVPTEQAR----------AY-----AEK-N----------- 148
RAB41   Q5JT25 E------HVRAERGD--------------------DVVI-MLLGNKIDLDN--------------------------KRQVTAEQGE----------EK-----SRN-L----------- 166
RAB42   Q8N4Z0 Q------EVMATQGP-------------------DKVIF-LLVGHKSDLQS------T--------------------RCVSAQEAE----------EL-----AAS-L----------- 151
RAB43   Q86YS6 E------DVRKYAGS--------------------NIVQ-LLIGNKSDLSE------L--------------------REVSLAEAQ----------SL-----AEH-Y----------- 153
RAB44   Q7Z6P3 D------CLQDAGSD--------------------GVVI-LLLGNKMDCEE------E--------------------RQVSVEAGQ----------QL-----AQE-L----------- 968
RASEF   Q8IZ41 D------MIEDAAHE--------------------TVPI-MLVGNKADIRDTAATEGQ--------------------KCVPGHFGE----------KL-----AMT-Y----------- 682
DIRAS1  O95057 K------LIV--QIKG--------------S-V-EDIPV-MLVGNKCDET------Q---------------------REVDTREAQ----------A-----VAQEW-----------K 143
DIRAS2  Q96HU8 E------QIC--EIKG--------------D-V-ESIPI-MLVGNKCDESP-----S---------------------REVQSSEAE----------A-----LARTW-----------K 144
DIRAS3  O95661 E------LIC--KIKG--------------NNL-HKFPI-VLVGNKSDDT------H---------------------REVALNDGA----------T-----CAMEW-----------N 174
ERAS    Q7Z444 --------IW--ATWG----------------PHPAQPL-VLVGNKCDLV------TT--------------------AGDAHAAAA----------A-----LAHSW-----------G 174
HRAS    P01112 E------QIK--RVKD----------------S-DDVPM-VLVGNKCDLA------A---------------------RTVESRQAQ----------D-----LARSY-----------G 138
KRAS    P01116 E------QIK--RVKD----------------S-EDVPM-VLVGNKCDLP------S---------------------RTVDTKQAQ----------D-----LARSY-----------G 138
MRAS    O14807 Q------LIL--RVKD----------------R-ESFPM-ILVANKVDLM------HL--------------------RKITREQGK----------E-----MATKH-----------N 149
NKIRAS1 Q9NYS0 K------EID--KFKD----------------K-KEVAI-VVLGNKIDLS------EQ--------------------RQVDAEVAQ----------Q-----WAKSE-----------K 143
NKIRAS2 Q9NYR9 K------EID--KSKD----------------K-KEVTI-VVLGNKCDLQ------EQ--------------------RRVDPDVAQ----------H-----WAKSE-----------K 143
NRAS    P01111 E------QIK--RVKD----------------S-DDVPM-VLVGNKCDLP------T---------------------RTVDTKQAH----------E-----LAKSY-----------G 138
RALA    P11233 E------QIL--RVKE----------------D-ENVPF-LLVGNKSDLE------DK--------------------RQVSVEEAK----------N-----RAEQW-----------N 150
RALB    P11234 E------QIL--RVKA----------------EEDKIPL-LVVGNKSDLE------ER--------------------RQVPVEEAR----------S-----KAEEW-----------G 151
RAP1A   P62834 E------QIL--RVKD----------------T-EDVPM-ILVGNKCDLE------DE--------------------RVVGKEQGQ----------N-----LARQW----------CN 140
RAP1B   P61224 E------QIL--RVKD----------------T-DDVPM-ILVGNKCDLE------DE--------------------RVVGKEQGQ----------N-----LARQW----------NN 140
RAP2A   P10114 D------QII--RVKR----------------Y-EKVPV-ILVGNKVDLE------SE--------------------REVSSSEGR----------A-----LAEEW-----------G 139
RAP2B   P61225 D------QII--RVKR----------------Y-ERVPM-ILVGNKVDLE------GE--------------------REVSYGEGK----------A-----LAEEW-----------S 139
RAP2C   Q9Y3L5 D------QIV--RVKR----------------Y-EKVPL-ILVGNKVDLE------PE--------------------REVMSSEGR----------A-----LAQEW-----------G 139
RASD1   Q9Y272 Q------QIL--DTKS--------CLKNKTKEN-VDVPL-VICGNKGDRDF-----Y---------------------REVDQREIE----------QLV---GDDPQ-----------R 170
RASD2   Q96D21 K------QIL--EVKS--------CLKNKTKEA-AELPM-VICGNKNDHGE-----LC--------------------RQVPTTEA-----------ELL---VSGDE-----------N 165
RASL10A Q92737 Q------RIAETRPAG----------------A-PEAPI-LVVGNKRDRQ------RL--------------------RFGPRRALA----------AL----VRRGW-----------R 153
RASL10B Q96S79 Q------QILETRVIG----------------T-SETPI-IIVGNKRDLQ------RG--------------------RVIPRWNVS----------HL----VRKTW-----------K 152
RASL11A Q6T310 Q------HIR--KVHP----------------D-SKAPV-IIVGNKGDLL------HA--------------------RQVQTQDGI----------Q-----LANEL-----------G 170
RASL11B Q9BPW5 Q------HVQ--QLHL----------------G-TRLPV-VVVANKADLL------HI--------------------KQVDPQLGL----------Q-----LASML-----------G 175
RASL12  Q9NYN1 E------LLALHAKET----------------Q-RSIPA-LLLGNKLDMA------QY--------------------RQVTKAEGV----------A-----LAGRF-----------G 157
RERG    Q96A58 N------ILD--EIKK----------------P-KNVTL-ILVGNKADLD------HS--------------------RQVSTEEGE----------K-----LATEL-----------A 141
RERGL   G5EA41 ------------------------------------------------------------------------------------------------------------------------
RHEB    Q15382 G------KLL--DMVG----------------K-VQIPI-MLVGNKKDLH------ME--------------------RVISYEEGK----------A-----LAESW-----------N 142
RHEBL1  Q8TAI7 Q------KLH--EGHG----------------K-TRVPV-VLVGNKADLS------PE--------------------REVQAVEGK----------K-----LAESW-----------G 142
RIT1    Q92963 Q------LIY--RVRR----------------T-DDTPV-VLVGNKSDLK------QL--------------------RQVTKEEGL----------A-----LAREF-----------S 157
RIT2    Q99578 E------LIF--QVRH----------------T-YEIPL-VLVGNKIDLE------QF--------------------RQVSTEEGL----------S-----LAQEY-----------N 156
RRAS    P10301 T------QIL--RVKD----------------R-DDFPV-VLVGNKADLE------SQ--------------------RQVPRSEAS----------A-----FGASH-----------H 165
RRAS2   P62070 R------QIL--RVKD----------------R-DEFPM-ILIGNKADLD------HQ--------------------RQVTQEEGQ----------Q-----LARQL-----------K 150
GEM     P55040 I------QLR--RARQ----------------T-EDIPI-ILVGNKSDLV------RC--------------------REVSVSEGR----------A-----CAVVF-----------D 214
REM1    O75628 I------QLR--RTHQ----------------A-DHVPI-ILVGNKADLA------RC--------------------REVSVEEGR----------A-----CAVVF-----------D 218
REM2    Q8IYK8 L------RLR--AGRP----------------H-HDLPV-ILVGNKSDLA------RS--------------------REVSLEEGR----------H-----LAGTL-----------S 252
RRAD    P55042 V------QLR--RARQ----------------T-DDVPI-ILVGNKSDLV------RS--------------------REVSVDEGR----------A-----CAVVF-----------D 226
CDC42   P60953 P------EITH---HC------------------PKTPF-LLVGTQIDLRDDPS-TIE-------KLAKN------KQKPITPETAE----------KL-----ARDL------------ 149
RAC1    P63000 P------EVRH---HC------------------PNTPI-ILVGTKLDLRDDKD-TIE-------KLKEK------KLTPITYPQGL----------AM-----AKEI------------ 149
RAC2    P15153 P------EVRH---HC------------------PSTPI-ILVGTKLDLRDDKD-TIE-------KLKEK------KLAPITYPQGL----------AL-----AKEI------------ 149
RAC3    P60763 P------EVRH---HC------------------PHTPI-LLVGTKLDLRDDKD-TIE-------RLRDK------KLAPITYPQGL----------AM-----AREI------------ 149
RHOA    P61586 P------EVKH---FC------------------PNVPI-ILVGNKKDLRNDEH-TRR-------ELAKM------KQEPVKPEEGR----------DM-----ANRI------------ 151
RHOB    P62745 P------EVKH---FC------------------PNVPI-ILVANKKDLRSDEH-VRT-------ELARM------KQEPVRTDDGR----------AM-----AVRI------------ 151
RHOBTB1 O94844 P------EIKH---FC------------------PRTPV-ILVGCQLDLRYADLEAVN-------RARRPLARPIKRGDILPPEKGR----------EV-----AKEL------------ 181
RHOBTB2 Q9BYZ6 P------EIKH---FC------------------PRAPV-ILVGCQLDLRYADLEAVN-------RARRPLARPIKPNEILPPEKGR----------EV-----AKEL------------ 181
RHOC    P08134 P------EVKH---FC------------------PNVPI-ILVGNKKDLRQDEH-TRR-------ELAKM------KQEPVRSEEGR----------DM-----ANRI------------ 151
RHOD    O00212 P------EVNH---FC------------------KKVPI-IVVGCKTDLCKDKS-LVN-------KLRRN------GLEPVTYHRGQ----------EM-----ARSV------------ 163
RHOF    Q9HBH0 P------EVTH---FC------------------RGIPM-VLIGCKTDLRKDKE-QLR-------KLRAA------QLEPITYMQGL----------SA-----CEQI------------ 165
RHOG    P84095 P------EVCH---HC------------------PDVPI-LLVGTKKDLRAQPD-TLR-------RLKEQ------GQAPITPQQGQ----------AL-----AKQI------------ 149
RHOH    Q15669 G------EIRS---NL------------------PCTPV-LVVATQTDQREMGP---------------H------RASCVNAMEGK----------KL-----AQDV------------ 143
RHOJ    Q9H4E5 P------ELKD---CM------------------PHVPY-VLIGTQIDLRDDPK-TLA-------RLLYM------KEKPLTYEHGV----------KL-----AKAI------------ 167
RHOQ    P17081 P------ELKE---YA------------------PNVPF-LLIGTQIDLRDDPK-TLA-------RLNDM------KEKPICVEQGQ----------KL-----AKEI------------ 155
RHOU    Q7L0Q8 P------EIRC---HC------------------PKAPI-ILVGTQSDLREDVK-VLI-------ELDKC------KEKPVPEEAAK----------LC-----AEEI------------ 195
RHOV    Q96L33 P------EIRT---HN------------------PQAPV-LLVGTQADLRDDVN-VLI-------QLDQG-----GREGPVPQPQAQ----------GL-----AEKI------------ 178
RND1    Q92730 T------EILD---YC------------------PSTRV-LLIGCKTDLRTDLS-TLM-------ELSHQ------KQAPISYEQGC----------AI-----AKQL------------ 159
RND2    P52198 G------ETQE---FC------------------PNAKV-VLVGCKLDMRTDLA-TLR-------ELSKQ------RLIPVTHEQGT----------VL-----AKQV------------ 153
RND3    P61587 G------EIQE---FC------------------PNTKM-LLVGCKSDLRTDVS-TLV-------ELSNH------RQTPVSYDQGA----------NM-----AKQI------------ 169
RAN     P62826 R------DLVRVCE----------------------NIPI-VLCGNKVDIKD---------------------------RKVKAKSIV----------FH-----RKK------------- 142
RHOT1   Q8IXI2 P------LINERTDKD------------------SRLPL-ILVGNKSDLVEYSS----------------------METILPI---------------M-----NQYT------------ 139
RHOT2   Q8IXI1 P------LVNGGTTQG------------------PRVPI-ILVGNKSDLRSGSS----------------------MEAVLPI---------------M-----SQFP------------ 139
IFT22   Q9H7X7 S------CFVQQPSLQ------------------DTQCM-LIAHHKPGSGDDKG--SL--------------------SLSPPLNKL----------KLVHSNLEDD------PEEI--R 158
RABL2A  Q9UBK7 T------ELREFRP----------------------EIPC-IVVANKIDDIN------------------------------VTQKSF----------NF-----AKK-F----------- 151
RABL2B  Q9UNT1 T------ELREFRP----------------------EIPC-IVVANKIDDIN------------------------------VTQKSF----------NF-----AKK-F----------- 151
RABL3   Q5HYI8 L------EALNRDLVPTGVLVTNGDYDQEQF-ADNQIPL-LVIGTKLDQIH-----ET--------------------KRHEVLTRT----------AF----LAEDFN----PEEINLD 179
RABL6   Q3YEC7 P------KVP------------------------THVPV-CVLGNYRDMG------EH--------------------RVILPDDVR----------DFI-DNLDRPP-----------G 203
RRAGA   Q7L523 YYQSCLEAILQN---------------------SPDAKI-FCLVHKMDLVQEDQRD-------------------------LIFKER-----EE---DL-R----RLS------------ 152
RRAGB   Q5VZM2 YYQSCLEAILQN---------------------SPDAKI-FCLVHKMDLVQEDQRD-------------------------LIFKER-----EE---DL-R----RLS------------ 213
RRAGC   Q9HB90 I-------TVSKAYK-----------------VNPDMNF-EVFIHKVDGLSDDHKIET--------------------QRDIHQRAN-DDLADAGLEKL-H----LSF------------ 215
RRAGD   Q9NQL2 L-------TVTRAYK-----------------VNTDINF-EVFIHKVDGLSDDHKIET--------------------QRDIHQRAN-DDLADAGLEKI-H----LSF------------ 216


ARF1    P84077 ----YIQATCAT-S---G------D-GL---YEG----LDWLSNQLRNQK---------------------------------------------------------------------- 181
ARF3    P61204 ----YIQATCAT-S---G------D-GL---YEG----LDWLANQLKNKK---------------------------------------------------------------------- 181
ARF4    P18085 ----YVQATCAT-Q---G------T-GL---YEG----LDWLSNELSKR----------------------------------------------------------------------- 180
ARF5    P84085 ----YVQATCAT-Q---G------T-GL---YDG----LDWLSHELSKR----------------------------------------------------------------------- 180
```

```
741
ARF6     P62330 ----YVQPSCAT-S---G------D-GL---YEG----LTWLTSNYKS------------------------------------------------------------------------ 175
ARL1     P40616 ----QIFKTSAT-K---G------T-GL---DEA----MEWLVETLKSRQ---------------------------------------------------------------------- 181
ARL2     P36404 ----CIQGCSAV-T---G------E-NL---LPG----IDWLLDDISSRIFTAD------------------------------------------------------------------ 184
ARL3     P36405 ----QIQSCSAL-T---G------E-GV---QDG----MNWVCKNVNAKKK--------------------------------------------------------------------- 182
ARL4A    P40617 ----HLQPTCAI-I---G------D-GL---KEG----LEKLHDMIIKRRKMLRQQKKKR------------------------------------------------------------ 200
ARL4C    P56559 ----HVQPACAI-I---G------E-GL---TEG----MDKLYEMILKRRKSLKQKKKR------------------------------------------------------------- 192
ARL4D    P49703 ----HVQGCSAV-D---G------L-GL---QQG----LERLYEMILKRKKAARGGKKRR------------------------------------------------------------ 201
ARL5A    Q9Y689 ----HIQACCAL-T---G------E-GL---CQG----LEWMMSRLKIR----------------------------------------------------------------------- 179
ARL5B    Q96KC2 ----HIQSCCAL-T---G------E-GL---CQG----LEWMTSRIGVR----------------------------------------------------------------------- 179
ARL5C    A6NH57 ----HIQGCCAL-T---R------E-GL---PAR----LQWMESQAAAN----------------------------------------------------------------------- 179
ARL6     Q9H0F7 ----HICASDAI-K---G------E-GL---QEG----VDWLQDQIQTVKT--------------------------------------------------------------------- 186
ARL8A    Q96BM9 ----CCYSISCK-E---K------D-NI---DIT----LQWLIQHSKSRRS--------------------------------------------------------------------- 186
ARL8B    Q9NVJ2 ----CCYSISCK-E---K------D-NI---DIT----LQWLIQHSKSRRS--------------------------------------------------------------------- 186
ARL9     Q6T311 ----FLFGTYLT-K---NGSEIPS---T---MQD----AKDLIAQLAADVQ--------------------------------------------------------------------- 187
ARL10    Q8N8L6 ----FLLAASIA-P---AGPTFEEP-GT---VHI----WKLLLELLS------------------------------------------------------------------------- 244
ARL11    Q969Q4 ----ELRGCSAL-T---G------E-GL---PEA----LQSLWSLLKSRSCMCLQARAHGAE---------------R-----------------------------GDSKRS------- 196
ARL13A   Q5H913 ----RVEPCSAI-R---NLERRNHQ-PI---VEG----LRWLLAVIDTCQLPPTSSISISKN---------------N-----------------------------TGSGERCS----- 215
ARL13B   Q3SXY8 ----QIEPCSAI-S---GYGKKIDK-SI---KKG----LYWLLHVIARDFDALNERIQKETT---------------E-----------------------------QRALEEQE----- 215
ARL14    Q8N4G2 ----YVQPCCAL-T---G------E-GL---AQG----FRKLTGFVKSHMKSRGDTLAFFKQ---------------N------------------------------------------ 192
ARL15    Q9NXU5 ----ILQPCSLD-D---M------D-AL---KDS----FSQLINLLEEKDHEAVRM---------------------------------------------------------------- 204
ARL16    Q0P5N6 ----TTAEISAR-E---G------T-GL---AGV----LAWLQATHRAND---------------------------------------------------------------------- 197
ARL17    Q8IVW1 ----GPWGCKDL-S---SG---FPS-FL---TSS----ILWKSAVVK------------------------------------------------------------------------- 177
ARFRP1   Q13795 ----LTQACSAL-T---G------K-GV---REG----IEWMVKCVVRNVHRPPRQRDIT------------------------------------------------------------ 201
SAR1A    Q9NR31 ----EVFMCSVL-K---R------Q-GY---GEG----FRWLSQYID------------------------------------------------------------------------- 198
SAR1B    Q9Y6B6 ----EVFMCSVL-K---R------Q-GY---GEG----FRWMAQYID------------------------------------------------------------------------- 198
TRIM23   P36406 ----YIQGCDAR-S---G------M-GL---YEG----LDWLSRQLVAAGVLDVA----------------------------------------------------------------- 574
IFT27    Q9BW83 -GL-ECFETSVK-E---M------E-NF---EAP----FHCLAKQFHQLYREKVEVFRALA----------------------------------------------------------- 186
RAB1A    P62820 -GI-PFLETSAK-N---A------T-NV---EQS----FMTMAAEIKKRMGPGATAGGAEK---S--NVKI---QS-------------------------------------------- 194
RAB1B    Q9H0U4 -GI-PFLETSAK-N---A------T-NV---EQA----FMTMAAEIKKRMGPGAASGG-ER---P--NLKI---DS-------------------------------------------- 190
RAB1C    Q92928 -GI-PFLETSAK-N---A------T-NV---EQA----FMTMAAEIKKQMGPGAASGG-ER---P--NLKI---DS-------------------------------------------- 190
RAB2A    P61019 -GL-IFMETSAK-T---A------S-NV---EEA----FINTAKEIYEKIQEGVFDINNEA-------NGI---KIGP-------------------------------QHAAT------ 194
RAB2B    Q8WUD1 -GL-IFMETSAK-T---A------C-NV---EEA----FINTAKEIYRKIQQGLFDVHNEA-------NGI---KIGP-------------------------------QQSIS------ 194
RAB3A    P20336 -GF-EFFEASAK-D---N------I-NV---KQT----FERLVDVICEKMSESLDTADPAV---TGAKQGP---QLSD------------------------------------------ 209
RAB3B    P20337 -GF-DFFEASAK-E---N------I-SV---RQA----FERLVDAICDKMSDSLDT-DPSM---LGSSKNT---RLSD------------------------------------------ 208
RAB3C    Q96E17 -GF-EFFETSAK-D---N------I-NV---KQT----FERLVDIICDKMSESLET-DPAI---TAAKQNT---RLKE------------------------------------------ 216
RAB3D    O95716 -GF-EFFEASAK-E---N------I-NV---KQV----FERLVDVICEKMNESLEPSSSS----GSNGKGP---AVGD------------------------------------------ 208
RAB4A    P20338 -EL-MFLETSAL-T---G------E-NV---EEA----FVQCARKILNKIESGELDPERMG-------SGI---QYGD-------------------------------AALRQ------ 201
RAB4B    P61018 -EL-MFLETSAL-T---G------E-NV---EEA----FLKCARTILNKIDSGELDPERMG-------SGI---QYGD-------------------------------ASLRQ------ 196
RAB5A    P20339 -SL-LFMETSAK-T---S------M-NV---NEI----FMAIAKKLPKNEPQNPG---A---NS-ARGRGV---DLTE------------------------------------------ 203
RAB5B    P61020 -SL-LFMETSAK-T---A------M-NV---NDL----FLAIAKKLPKSEPQNLG---G---AA-GRSRGV---DLHE------------------------------------------ 203
RAB5C    P51148 -SL-LFMETSAK-T---A------M-NV---NEI----FMAIAKKLPKNEPQNAT---G---AP-GRNRGV---DLQE------------------------------------------ 204
RAB6A    P20340 -NV-MFIETSAK-A---G------Y-NV---KQL----FRRVAAALPGMESTQDRSR-----------EDMIDIKLEK------------------------------------------ 195
RAB6B    Q9NRW1 -SV-MFIETSAK-T---G------Y-NV---KQL----FRRVASALPGMENVQEKSK-----------EGMIDIKLDK------------------------------------------ 195
RAB6C    Q9H0N0 -NV-TFIETRAK-A---G------Y-NV---KQL----FRRVAAALPGMESTQDGSR-----------EDMSDIKLEK------------------------------------------ 195
RAB7A    P51149 -NNIPYFETSAK-E---A------I-NV---EQA----FQTIARNALKQETEVELYNEFPEP------IKL---DKND------------------------------------------ 196
RAB7B    Q96AH8 --DIPYFEVSAK-N---D------I-NV---VQA----FEMLASRALSRYQSILE-N--HLT------ESI---KLS------------------------------------------- 190
RAB8A    P61006 -GI-KFMETSAK-A---N------I-NV---ENA----FFTLARDIKAKMDKKLEGNSPQ-----GSNQGV---KI-------------------------------------------- 191
RAB8B    Q92930 -GI-KFLETSAK-S---S------A-NV---EEA----FFTLARDIMTKLNRKMNDSNSA-----GAGGPV---KI-------------------------------------------- 191
RAB9A    P51151 -GDYPYFETSAK-D---A------T-NV---AAA----FEEAVRRVLATEDRSDHLI---QT------DTV---NLHR------------------------------------------ 192
RAB9B    Q9NP90 -GDYPYLETSAK-D---D------T-NV---TVA----FEEAVRQVLAVEEQLEHCM---LG------HTI---DLNS------------------------------------------ 192
RAB10    P61026 -GI-RFFETSAK-A---N------I-NI---EKA----FLTLAEDILRKTPVKEPNSENVD---ISSGGGV---T--------------------------------------------- 193
RAB11A   P62491 -GL-SFIETSAL-D---S------T-NV---EAA----FQTILTEIYRIVSQKQMSDRRENDMSPSNNV-V---PIHV------------------------------------------ 200
RAB11B   Q15907 -NL-SFIETSAL-D---S------T-NV---EEA----FKNILTEIYRIVSQKQIADRAAHDESPGNNV-V---DISV------------------------------------------ 200
RAB12    Q6IQ22 -GM-RFCEASAK-D---N------F-NV---DEI----FLKLVDDILKKMPLDILRNELS-------NSIL---SLQP-------------------------------EPE-------- 229
RAB13    P51153 -GI-RFFETSAK-S---S------M-NV---DEA----FSSLARDILLKSGGRRSGNGNKP---PSTD--L---K--------------------------------------------- 190
RAB14    P61106 -GL-LFLEASAK-T---G------E-NV---EDA----FLEAAKKIYQNIQDGSLDLNAAE-------SGV---QHKP-------------------------------SAPQG------ 199
RAB15    P59190 -GM-DFYETSAC-T---N------L-NI---KES----FTRLTELVLQAHRKELEGLRMRA---S-NELAL---AELE-------------------------------EEE-------- 197
RAB17    Q9H0T7 -KL-LFMETSAK-L---N------H-QV---SEV----FNTVAQELLQRSDEEGQA-----------LRGDAAVALN------------------------------------------- 200
RAB18    Q9NP72 -SM-LFIEASAK-T---C------D-GV---QCA----FEELVEKIIQTPGLWESENQ---------NKGV---KLSH-------------------------------REE-------- 192
RAB19    A4D1S5 -GLLAVLETSAK-E---S------K-NI---EEV----FVLMAKELIARNSLHLYGESALN--------GL---PLDS-------------------------------SPV-------- 203
RAB20    Q9NX57 AAEQMCFETSAK-T---G------Y-NV---DLL----FETLFDLVVPMILQQRAERPSHT---------V---DISS------------------------------------------ 221
RAB21    Q9UL25 -GA-KHYHTSAK-Q---N------K-GI---EEL----FLDLCKRMIETAQVDERAKGNGSSQPGTARRGVQIIDDEP------------------------------------------ 212
RAB22A   Q9UL26 -HA-IFVETSAK-N---A------I-NI---NEL----FIEISRRIPSTDANLPS---G--------GKGF---KLRR------------------------------------------ 184
RAB23    Q9ULC3 -KL-RFYRTSVK-E---D------L-NV---NEV----FKYLAEKYLQKLKQQIAEDPELTH-SSSNKIGV---FNTSG-----------------------------GSHSGQ-NSGTL 209
RAB24    Q969Q5 -KA-QLFETSSK-T---G------Q-SV---DEL----FQKVAEDYVSVAAFQVMTE----------DKGV---DLGQ------------------------------------------ 191
RAB25    P57735 -GL-LFLETSAL-D---S------T-NV---ELA----FETVLKEIFAKVSKQRQNSIRTNAITLGSAQ--------------------------------------------------- 196
RAB26    Q9ULW5 -GL-PFMETSAK-T---G------L-NV---DLA----FTAIAKELKQRSMKAPSEPRF---------------RLHD-------------------------------YVK-------- 245
RAB27A   P51159 -GI-PYFETSAA-N---G------T-NI---SQA----IEMLLDLIMKRMERCVDKSWIPEGVVRSNGH-A---STD------------------------------------------- 208
RAB27B   O00194 -GI-PYFETSAA-T---G------Q-NV---EKA----VETLLDLIMKRMEQCVEKTQIPDTVNGGNSG-N------------------------------------------------- 205
RAB28    P51157 FS---SHFVSAK-T---G------D-SV---FLC----FQKVAAEILGIKLNKAEI----EQ------SQR---VVKA--DIVNYN------------QE-------------------- 204
RAB29    O14966 -GFTGWTETSVK-E---N------K-NI---NEA----MRVLIEKMMRNSTEDIMSLSTQG-------DYI---NLQT------------------------------------------ 195
RAB30    Q15771 -DM-YYLETSAK-E---S------D-NV---EKL----FLDLACRLISEARQNTLVNNV--------------------------------------------------SSP-------- 186
RAB31    Q13636 -GA-IVVETSAK-N---A------I-NI---EEL----FQGISRQIPPLDPHENG---N--------NGTI---KVEK------------------------------------------ 184
RAB32    Q13637 -GFAGWFETSAK-D---N------I-NI---EEA----ARFLVEKILVNHQSFPNE-ENDV-------DKI---KLDQ------------------------------------------ 213
RAB33A   Q14088 -NM-LLFETSAK-DPKES------Q-NV---ESI----FMCLACRLKAQKSLLYRDAERQQ----GKVQKL---EF-------------------------------------------- 225
RAB33B   Q9H082 -SM-PLFETSAK-NPNDN------D-HV---EAI----FMTLAHKLKSHKPLMLSQPPDN-----G----I---IL-------------------------------------------- 217
RAB34    Q9BZG1 -KA-EYWAVSSL-T---G------E-NV---REF----FFRVAALTFEANVLAELEKSGARRI----GDVV---RINS------------------------------------------ 241
RAB35    Q15286 -GI-QLFETSAK-E---N------V-NV---EEM----FNCITELVLRAKKDNLAKQQQQQ---Q--NDVV---KL-------------------------------------------- 190
RAB36    O95755 -QA-EYWSVSAK-T---G------E-NV---KAF----FSRVAALAFEQSVLQDLERQSSARLQVGNGDLI---QMEG------------------------------------------ 316
RAB37    Q96AX2 -GV-PFLETSAK-T---G------M-NV---ELA----FLAIAKELKYRAGHQADEPSF---------------QIRD-------------------------------YVE-------- 211
RAB38    P57729 -GFVGWFETSAK-E---N------I-NI---DEA----SRCLVKHILANECDLMESIEPDV-------VKP---HLT------------------------------------------- 198
RAB39A   Q14964 -GM-KYIETSAK-D---A------T-NV---EES----FTILTRDIYELIKKGEICIQDGW-------EGV---KSGF------------------------------------------ 197
RAB39B   Q96DA2 -GM-KYIETSAR-D---A------I-NV---EKA----FTDLTRDIYELVKRGEITIQEGW-------EGV---KSGF------------------------------------------ 193
RAB40A   Q8WXH6 -GV-TFFEVSPL-C---N------F-NI---IES----FTELARIVLLRHRMNWLGRPSKV---------L---S--------------------------------------------- 191
RAB40B   Q12829 -GV-TFFEVSPL-C---N------F-NI---TES----FTELARIVLLRHGMDRLWRPSKV---------L---S--------------------------------------------- 191
RAB40C   Q96S21 -CM-TFFEVSPL-C---N------F-NV---IES----FTELSRIVLMRHGMEKIWRPNRV---------F---S--------------------------------------------- 191
RAB41    Q5JT25 -NV-MFIETSAK-T---G------Y-NV---KKL----FRRVASALLSTRTSPP-PK-----------EGTVEIELES------------------------------------------ 212
RAB42    Q8N4Z0 -GM-AFVETSVK-N---N------C-NV---DLA----FDTLADAIQQALQQGDIKLEEGW-------GGV---RLIH------------------------------------------ 199
RAB43    Q86YS6 -DILCAIETSAK-D---S------S-NV---EEA----FLRVATELIMRHGGPLFSEKSPD--------HI---QLNS-------------------------------KDI-------- 204
RAB44    Q7Z6P3 -GV-YFGECSAA-L---G------H-NI---LEP----VVNLARSLRMQEEGLKD-SL-----------V---KV--------------------------------------------1008
RASEF    Q8IZ41 -GA-LFCETSAK-D---G------S-NI---VEA----VLHLAREVKKRTDKDDSRSITNL---TGTN---------------------------------------------------- 727
DIRAS1   O95057 CA---FMETSAK-M---N------Y-NV---KEL----FQELLTLETRRNMSLN------ID------GKR---SGK----------------QKRTD---------------------- 189
DIRAS2   Q96HU8 CA---FMETSAK-L---N------H-NV---KEL----FQELLNLEKRRTVSLQ------ID------GKK---SKQ----------------QKRKE---------------------- 190
DIRAS3   O95661 CA---FMEISAK-T---D------V-NV---QEL----FHMLLNYKKKPTTGLQ------EP------EKK---SQM----------------PNTTE---------------------- 220
ERAS     Q7Z444 AH---FVETSAK-T---R------Q-GV---EEA----FSLLVHEIQRVQEAMAKE----PM------ARS---CREK----TRH----------------------------------- 221
HRAS     P01112 IP---YIETSAK-T---R------Q-GV---EDA----FYTLVREIRQHKLRKLNP----P--------------DES----GPG----------------------------------- 180
KRAS     P01116 IP---FIETSAK-T---R------Q-RV---EDA----FYTLVREIRQYRLKKISK----E---------------EK----TPG----------------------------------- 179
MRAS     O14807 IP---YIETSAKDP---P------L-NV---DKA----FHDLVRVIRQQIPEKSQK----K--------------KKK----TKWR------------GD-------------------- 195
NKIRAS1  Q9NYS0 VR---LWEVTVT-D---R------K-TL---IEP----FTLLASKLSQPQSKSSFP----LP------GRK---NKGNS------N---------------------------------- 189
NKIRAS2  Q9NYR9 VK---LWEVSVA-D---R------R-SL---LEP----FVYLASKMTQPQSKSAFP----LS------RKN---K-GSG------S---------------------------------- 188
NRAS     P01111 IP---FIETSAK-T---R------Q-GV---EDA----FYTLVREIRQYRMKKLNS----S--------------DDG----TQG----------------------------------- 180
RALA     P11233 VN---YVETSAK-T---R------A-NV---DKV----FFDLMREIRARKMEDSKE----KN------GKK---KRKS---------------------L-------------------- 195
RALB     P11234 VQ---YVETSAK-T---R------A-NV---DKV----FFDLMREIRTKKMSENKD----KN------GKK---SSKN------------------------------------------ 195
RAP1A    P62834 CA---FLESSAK-S---K------I-NV---NEI----FYDLVRQINRKTPV-------------------------------------------------------------------- 171
RAP1B    P61224 CA---FLESSAK-S---K------I-NV---NEI----FYDLVRQINRKTPV-------------------------------------------------------------------- 171
RAP2A    P10114 CP---FMETSAK-S---K------T-MV---DEL----FAEIVRQMNYAAQP-------------------------------------------------------------------- 170
RAP2B    P61225 CP---FMETSAK-N---K------A-SV---DEL----FAEIVRQMNYAAQP-------------------------------------------------------------------- 170
RAP2C    Q9Y3L5 CP---FMETSAK-S---K------S-MV---DEL----FAEIVRQMNYSSLP-------------------------------------------------------------------- 170
RASD1    Q9Y272 CA---YFEISAK-K---N------S-SL---DQM----FRALFAMAKLPSEMSPD-----LH------RKV---SVQYC-DV-------LHKKALRNKKLLRAGSGGGGGDPGDAFGIVA 247
RASD2    Q96D21 CA---YFEVSAK-K---N------T-NV---DEM----FYVLFSMAKLPHEMSPA-----LH------RKI---SVQYG-DA-------FHPRPFCMRRV----------KEMDAYGMVS 232
RASL10A  Q92737 CG---YLECSAK-Y---N------W-HV---LRL----FRELLRCA-LVRARPAHP-------------------------------------ALRLQGA-------------------- 194
RASL10B  Q96S79 CG---YVECSAK-Y---N------W-HI---LLL----FSELLKSVGCARCKHVHA-------------------------------------ALRFQGA-------------------- 194
RASL11A  Q6T310 SL---FLEISTSEN---Y------E-DV---CDV----FQHLCKEVSKMHG--LSG----ER------RRA---SIIPR-PRSPNM------------QD-------------------- 222
RASL11B  Q9BPW5 CS---FYEVSVSEN---Y------N-DV---YSA----FHVLCKEVSHKQQPSSTP----EK------RRT---SLIPR-PKSPNM------------QD-------------------- 229
RAS12    Q9NYN1 CL---FFEVSACLD---F------E-HV---QHV----FHEAVREARRELEKS--P----LT------RPL---FISEE-RALPHQ------------AP-------------------- 209
RERG     Q96A58 CA---FYECSACTG---E------G-NI---TEI----FYELCREVRRRRMVQ--G----KT------RRR---SSTTH----------------------------------------- 185
RERGL    G5EA41 ------------------------------------------------------------------------------------------------------------------------
RHEB     Q15382 AA---FLESSAK-E---N------Q-TA---VDV----FRRIILEAEKMDGAASQG----K----------------------------------------------------------- 178
RHEBL1   Q8TAI7 AT---FMESSAR-E---N------Q-LT---QGI----FTKVIQEIARVENSYGQE---------------------------------------------------------------- 177
RIT1     Q92963 CP---FFETSAA-Y---R------Y-YI---DDV----FHALVREIRRKEKEAVLA----ME------KKS---KPKN----SVWK------------RL-------------------- 207
RIT2     Q99578 CG---FFETSAA-L---R------F-CI---DDA----FHGLVREIRKKESMPS-L----ME------KKL---KRKD----SLWK------------KL-------------------- 205
RRAS     P10301 VA---YFEASAK-L---R------L-NV---DEA----FEQLVRAVRKYQEQELPP----S--------------PPS----AP------------------------------------ 206
RRAS2    P62070 VT---YMEASAK-I---R------M-NV---DQA----FHELVRVIRKFQEQECPP----S--------------PEP----TRK----------------------------------- 192
GEM      P55040 CK---FIETSAA-V---Q------H-NV---KEL----FEGIVRQVRLRRDSKEKN----ER-----------------RLAYQKRKESMPRKARRFWGK-------------------- 272
REM1     O75628 CK---FIETSAT-L---Q------H-NV---AEL----FEGVVRQLRLRRRDSAAK-----------------------EPPAPRRPASLAQRARRFLAR-------------------- 274
REM2     Q8IYK8 CK---HIETSAA-L---H------H-NT---REL----FEGAVRQIRLRRGRNHAG----GQ------RPD---PGSPEGPAPPARRESLTKKAKRFLAN-------------------- 318
```

```
RRAD     P55042 CK---FIETSAA-L---H------H-NV---QAL----FEGVVRQIRLRRDSKEAN----AR----------------RQAGTRRRESLGKKAKRFLGR-------------------- 284
CDC42    P60953 -KAVKYVECSAL-T---Q------K-GL---KNV----FDEAILAALEPPEPKK--S---R------------------------------------------------------------ 186
RAC1     P63000 -GAVKYLECSAL-T---Q------R-GL---KTV----FDEAIRAVLCPPPVKK-RK---R------------------------------------------------------------ 187
RAC2     P15153 -DSVKYLECSAL-T---Q------R-GL---KTV----FDEAIRAVLCPQPTRQ-QK---R------------------------------------------------------------ 187
RAC3     P60763 -GSVKYLECSAL-T---Q------R-GL---KTV----FDEAIRAVLCPPPVKK-PG---K------------------------------------------------------------ 187
RHOA     P61586 -GAFGYMECSAK-T---K------D-GV---REV----FEMATRAALQARRGKK-KS---------------------------------------------------------------- 188
RHOB     P62745 -QAYDYLECSAK-T---K------E-GV---REV----FETATRAALQKRYGSQ-N----------------------------------------------------------------- 187
RHOBTB1  O94844 -GL-PYYETSVF-D---Q------F-GI---KDV----FDNAIRAALISRRHLQFWKSHLK-------------KVQK-----------------P------------------------ 227
RHOBTB2  Q9BYZ6 -GI-PYYETSVV-A---Q------F-GI---KDV----FDNAIRAALISRRHLQFWKSHLR-------------NVQR-----------------P------------------------ 227
RHOC     P08134 -SAFGYLECSAK-T---K------E-GV---REV----FEMATRAGLQVRKNKR-RR--------------------------------------------------------------- 188
RHOD     O00212 -GAVAYLECSAR-L---H------D-NV---HAV----FQEAAEVALSSRGRNF-WRRIT------------------------------------------------------------ 203
RHOF     Q9HBH0 -RAALYLECSAK-F---R------E-NV---EDV----FREAAKVALSALKKAQ-RQKKRR----------------------------------------------------------- 206
RHOG     P84095 -HAVRYLECSAL-Q---Q------D-GV---KEV----FAEAVRAVLNPTPIKR--G---R------------------------------------------------------------ 186
RHOH     Q15669 -RAKGYLECSAL-S---N------R-GV---QQV----FECAVRTAVNQARRRN-RRRLFS------------------------------------------------------------ 184
RHOJ     Q9H4E5 -GAQCYLECSAL-T---Q------K-GL---KAV----FDEAILTIFHPKKKKK-RCSEGH------------------------------------------------------------ 208
RHOQ     P17081 -GACCYVECSAL-T---Q------K-GL---KTV----FDEAIIAILTPKKHTV-KKRIGS------------------------------------------------------------ 196
RHOU     Q7L0Q8 -KAASYIECSAL-T---Q------K-NL---KEV----FDAAIVAGIQYSDTQQ-QPKKSK-------------SRTP-----------------D------------------------ 241
RHOV     Q96L33 -RACCYLECSAL-T---Q------K-NL---KEV----FDSAILSAIEHKARLE------K-------------KLNA-----------------K------------------------ 219
RND1     Q92730 -GAEIYLEGSAF-T---S------EKSI---HSI----FRTASMLCLNKPSPLP-QKSPVR-------------SLSK-----------------R------------------------ 206
RND2     P52198 -GAVSYVECSSR-S---S------ERSV---RDV----FHVATVASLGRGHRQL-RRTDSR-------------RGMQ-----------------R------------------------ 200
RND3     P61587 -GAATYIECSAL-Q---S------ENSV---RDI----FHVATLACVNKTNKNV-KRNKSQ-------------RATK-----------------R------------------------ 216
RAN      P62826 -NL-QYYDISAK-S---N------Y-NF---EKP----FLWLARKLIGDPNLEFVAMPALAP------PEV---VMDP---------------------------------ALAAQ-YEHDL 201
RHOT1    Q8IXI2 -EIETCVECSAK-N---L------K-NI---SEL----FYYAQKAVLHPTGPLYCPE--EK-------------EMKP-----------------A------------------------ 184
RHOT2    Q8IXI1 -EIETCVECSAK-N---L------R-NI---SEL----FYYAQKAVLHPTAPLYDPE--AK-------------QLRP-----------------A------------------------ 184
IFT22    Q9H7X7 MEFIKYLKSIIN---------------------------SMSESRDREEMSIMT----------------------------------------------------------------- 185
RABL2A   Q9UBK7 -SL-PLYFVSAA-D---G------T-NV---VKL----FNDAIRLAVSYKQNSQDFMDEIFQ----ELENF---SLEQ---------------------------------E---------- 203
RABL2B   Q9UNT1 -SL-PLYFVSAA-D---G------T-NV---VKL----FNDAIRLAVSYKQNSQDFMDEIFQ----ELENF---SLEQ---------------------------------E---------- 203
RABL3    Q5HYI8 CTNPRYLAAGSS-N---A------V-KL---SRF----FDKVIEKRYFLREGNQIP------------------------GFPDR--------KRFG---------AGTLKSLHYD--- 236
RABL6    Q3YEC7 SSYFRYAESSMK-N---S------F-GLKYLHKFFNIPFLQLQRETLLRQLETNQLDMDATL------EEL---SVQQE------------------------------TEDQNYGIFL 272
RRAGA    Q7L523 ----RPLECACF-R---T--SIWDE-TL---YKA----WSSIVYQLIPNVQQLEMNLRNFAQ---------------I-----------------------------IEADEVLLFERAT 210
RRAGB    Q5VZM2 ----RPLECSCF-R---T--SIWDE-TL---YKA----WSSIVYQLIPNVQQLEMNLRNFAE---------------I-----------------------------IEADEVLLFERAT 271
RRAGC    Q9HB90 ----YLT--S-----------IYDH-SI---FEA----FSKVVQKLIPQLPTLENLLNIFIS---------------N-----------------------------SGIEKAFLFDVVS 266
RRAGD    Q9NQL2 ----YLT--S-----------IYDH-SI---FEA----FSKVVQKLIPQLPTLENLLNIFIS---------------N-----------------------------SGIEKAFLFDVVS 267


ARF1     P84077 ------------------------------------------------------------
ARF3     P61204 ------------------------------------------------------------
ARF4     P18085 ------------------------------------------------------------
ARF5     P84085 ------------------------------------------------------------
ARF6     P62330 ------------------------------------------------------------
ARL1     P40616 ------------------------------------------------------------
ARL2     P36404 ------------------------------------------------------------
ARL3     P36405 ------------------------------------------------------------
ARL4A    P40617 ------------------------------------------------------------
ARL4C    P56559 ------------------------------------------------------------
ARL4D    P49703 ------------------------------------------------------------
ARL5A    Q9Y689 ------------------------------------------------------------
ARL5B    Q96KC2 ------------------------------------------------------------
ARL5C    A6NH57 ------------------------------------------------------------
ARL6     Q9H0F7 ------------------------------------------------------------
ARL8A    Q96BM9 ------------------------------------------------------------
ARL8B    Q9NVJ2 ------------------------------------------------------------
ARL9     Q6T311 ------------------------------------------------------------
ARL10    Q8N8L6 ------------------------------------------------------------
ARL11    Q969Q4 ------------------------------------------------------------
ARL13A   Q5H913 SHSF------S---------TRTGMSK---------------EKRQH--LEQC--SIE-- 239
ARL13B   Q3SXY8 KQER------A---------ERVRKLR---------------EERKQNEQEQA--ELDGT 243
ARL14    Q8N4G2 ------------------------------------------------------------
ARL15    Q9NXU5 ------------------------------------------------------------
ARL16    Q0P5N6 ------------------------------------------------------------
ARL17    Q8IVW1 ------------------------------------------------------------
ARFRP1   Q13795 ------------------------------------------------------------
SAR1A    Q9NR31 ------------------------------------------------------------
SAR1B    Q9Y6B6 ------------------------------------------------------------
TRIM23   P36406 ------------------------------------------------------------
IFT27    Q9BW83 ------------------------------------------------------------
RAB1A    P62820 --------TP------------VKQSG----------------GGCC------------- 205
RAB1B    Q9H0U4 --------TP------------VKPAG----------------GGCC------------- 201
RAB1C    Q92928 --------TP------------VKPAG----------------GGCC------------- 201
RAB2A    P61019 -----NATHAG-------N-QGGQQAG----------------GGCC------------- 212
RAB2B    Q8WUD1 -----TSVGPSAS-QR--N-SRDIGSN----------------SGCC------------- 216
RAB3A    P20336 --------QQ-------------VPPH----------------QDCAC------------ 220
RAB3B    P20337 --------TP-------------PLLQ----------------QNCSC------------ 219
RAB3C    Q96E17 --------TP-------------PPPQ----------------PNCAC------------ 227
RAB3D    O95716 --------AP-------------APQP----------------SSCSC------------ 219
RAB4A    P20338 -----LRSPRR-------A-QAPNAQE----------------CGC-------------- 218
RAB4B    P61018 -----LRQPRS-------A-QAVAPQP----------------CGC-------------- 213
RAB5A    P20339 ------------------P-TQP--TR----------------NQCCSN----------- 215
RAB5B    P61020 ------------------Q-SQQ--NK----------------SQCCSN----------- 215
RAB5C    P51148 ------------------N-NPA--SR----------------SQCCSN----------- 216
RAB6A    P20340 ------------------P-QEQPVSE----------------GGCSC------------ 208
RAB6B    Q9NRW1 ------------------P-QEPPASE----------------GGCSC------------ 208
RAB6C    Q9H0N0 ------------------P-QEQTVSE----------------GGCSCYSPMS-----S- 214
RAB7A    P51149 ----------------------RAKAS----------------AESCSC----------- 207
RAB7B    Q96AH8 ----------------------PDQSR----------------SRCC------------- 199
RAB8A    P61006 --------TPD-------Q--QKRSSF----------------FRCVLL----------- 207
RAB8B    Q92930 --------TEN-------R--SKKTSF----------------FRCSLL----------- 207
RAB9A    P51151 ----------------------KPKPS----------------SSCC------------- 201
RAB9B    Q9NP90 ----------------------GSKAG----------------SSCC------------- 201
RAB10    P61026 ---------GW-------K--S---------------------KCC-------------- 200
RAB11A   P62491 --------PPT-------T-EN--KPK----------------VQCCQNI---------- 216
RAB11B   Q15907 --------PPT-------T-DGQKPNK----------------LQCCQNL---------- 218
RAB12    Q6IQ22 -------IPPE-------L--PPPRPH----------------VRCC------------- 244
RAB13    P51153 ------------------T--CDKKNT----------------NKCSLG----------- 203
RAB14    P61106 -----GRL-TS-------E-PQPQREG----------------CGC-------------- 215
RAB15    P59190 -------GKPE-------G--PAN-SS----------------KTCWC------------ 212
RAB17    Q9H0T7 ------------------K-GPA--RQ----------------AKCCAH----------- 212
RAB18    Q9NP72 ------------------G-Q-GGGAC----------------GGYCSVL---------- 206
RAB19    A4D1S5 --------LM---------AQGPSEK-----------------THCTC------------ 217
RAB20    Q9NX57 ------------------H-KPPKRTR----------------SGCCA------------ 234
RAB21    Q9UL25 ------------------Q-AQT--SG----------------GGCCSSG---------- 225
RAB22A   Q9UL26 ------------------Q-PSE--PK----------------RSCC------------- 194
RAB23    Q9ULC3 NGGDVINLRPN-K-QR--T-KKNRNPF----------------SSCSIP----------- 237
RAB24    Q969Q5 ------------------K-PNP--YF----------------YSCCHH----------- 203
RAB25    P57735 ---------AG-------Q-EPGPGEK----------------RACCISL---------- 213
RAB26    Q9ULW5 ----------------------REGRG----------------ASCCRP----------- 256
RAB27A   P51159 ------------------Q-LSEEKEK----------------GACGC------------ 221
RAB27B   O00194 ------------------L-DGEKPPE----------------KKCIC------------ 218
RAB28    P51157 ------------P-------MSRTVNPPRSS-------------MCAVQ----------- 221
RAB29    O14966 ------------------K-S----SS----------------WSCC------------- 203
RAB30    Q15771 --------LP----------GEGKSIS----------------YLTCCNFN--------- 203
RAB31    Q13636 ------------------P-TMQ--AS----------------RRCC------------- 194
RAB32    Q13637 ------------------E-TLRAENK----------------SQCC------------- 225
RAB33A   Q14088 --------------------PQEANSK----------------TSCPC------------ 237
RAB33B   Q9H082 --------------------KPEPKPA----------------MTCWC------------ 229
RAB34    Q9BZG1 ----------DDS-NL--Y-LTASKKK----------------PTCCP----------- 259
RAB35    Q15286 --------TK------------NSKRK----------------KRCC------------- 201
RAB36    O95755 ----------SPP-ET--Q-ESKRPSS----------------LGCC------------- 333
RAB37    Q96AX2 ----------------------SQKKR----------------SSCCSFM---------- 223
RAB38    P57729 --------------------STKVASC----------------SGCAKS----------- 211
RAB39A   Q14964 ----VPNTVHS-------S-EEAVKPR----------------KECFC------------ 217
RAB39B   Q96DA2 ----VPNVVHS-------S-EEVVKSE----------------RRCLC------------ 213
RAB40A   Q8WXH6 -------------------------LQ----------------DLCCRTIVSC-----TP 205
RAB40B   Q12829 -------------------------LQ----------------DLCCRAVVSC-----TP 205
```

```
RAB40C  Q96S21 -------------------------LQ----------------DLCCRAIVSC-----TP 205
RAB41   Q5JT25 ------------------F-E-----E----------------SGNRSYC---------- 222
RAB42   Q8N4Z0 ----KTQIPRS-------P--SRKQHS----------------GPCQC------------ 218
RAB43   Q86YS6 ------------------------GEG----------------WGCGC------------ 212
RAB44   Q7Z6P3 ------------------A-PKRPPKR----------------FGCCS------------ 1021
RASEF   Q8IZ41 --------------------SKKSPQM----------------KNCCNG----------- 740
DIRAS1  O95057 ---------------------------RVKG-------------KCTLM----------- 198
DIRAS2  Q96HU8 ---------------------------KLKG-------------KCVIM----------- 199
DIRAS3  O95661 ---------------------------KLLD-------------KCIIM----------- 229
ERAS    Q7Z444 -----------------------QKATCHC--------------GCSVA----------- 233
HRAS    P01112 --------------------------CMSCK--------------CVLS----------- 189
KRAS    P01116 --------------------------CVKIK-------------KCIIM----------- 189
MRAS    O14807 ----------------------RATGTHKL--------------QCVIL----------- 208
NKIRAS1 Q9NYS0 ------------S--------------------------------EN------------- 192
NKIRAS2 Q9NYR9 ------------L--------------------------------DG------------- 191
NRAS    P01111 --------------------------CMGLP--------------CVVM----------- 189
RALA    P11233 ------------A-------K-----RIRE--------------RCCIL----------- 206
RALB    P11234 ------------K-------K-----SFKE--------------RCCLL----------- 206
RAP1A   P62834 -----------------------EKKKPKKK-------------SCLLL----------- 184
RAP1B   P61224 -----------------------PGKARKKS-------------SCQLL----------- 184
RAP2A   P10114 -----------------------DKDDPCCS-------------ACNIQ----------- 183
RAP2B   P61225 -----------------------NGDEGCCS-------------ACVIL----------- 183
RAP2C   Q9Y3L5 -----------------------EKQDQCCT-------------TCVVQ----------- 183
RASD1   Q9Y272 PFARRPS-VHSDLMYIREK-ASAGSQAKDKE-------------RCVIS----------- 281
RASD2   Q96D21 PFARRPS-VNSDLKYIKAK-VLREGQARERD-------------KCTIQ----------- 266
RASL10A Q92737 ---------------------------LHPA-------------RCSLM----------- 203
RASL10B Q96S79 ---------------------------LRRN-------------RCAIM----------- 203
RASL11A Q6T310 ------------L-------KRRFKQALS----------------PKVKAPSA------- 240
RASL11B Q9BPW5 ------------L-------KRRFKQALS----------------AKVRTVTS------- 247
RASL12  Q9NYN1 ------------L-------TARHGLA-S----------------CTFNTLST------- 226
RERG    Q96A58 -----------------------VKQAIN----------------KMLTKISS------- 199
RERGL   G5EA41 ------------------------------------------------------------
RHEB    Q15382 ------------------------------S-------------SCSVM----------- 184
RHEBL1  Q8TAI7 ------------------------------R-------------RCHLM----------- 183
RIT1    Q92963 ------------K-------SPFRKKKDSVT----------------------------- 219
RIT2    Q99578 ------------K-------GSLKKKRENMT----------------------------- 217
RRAS    P10301 -----------------------RKKGGGC--------------PCVLL----------- 218
RRAS2   P62070 -----------------------EKDKKGC--------------HCVIF----------- 204
GEM     P55040 ------------IVAKNNK-NMAFKLK--SK-------------SCHDLSVL-------- 296
REM1    O75628 ------------LTARSAR-RRALKAR--SK-------------SCHNLAVL-------- 298
REM2    Q8IYK8 ------------LVPRNAK---FFKQR--SR-------------SCHDLSVL-------- 340
RRAD    P55042 ------------IVARNSR-KMAFRAK--SK-------------SCHDLSVL-------- 308
CDC42   P60953 -----------------------------------------RCVLL-------------- 191
RAC1    P63000 -----------------------------------------KCLLL-------------- 192
RAC2    P15153 -----------------------------------------ACSLL-------------- 192
RAC3    P60763 -----------------------------------------KCTVF-------------- 192
RHOA    P61586 -----------------------------------------GCLVL-------------- 193
RHOB    P62745 -----------------------------------------GCINCCKVL---------- 196
RHOBTB1 O94844 -LLQAPFLPPKAP-------PPVIKIPECPSMGTNEAACLLDNPLCADVLFI-------- 271
RHOBTB2 Q9BYZ6 -LLQAPFLPPKPP-------PPIIVVPDPPSSSEECPAHLLEDPLCADVILV-------- 271
RHOC    P08134 -----------------------------------------GCPIL-------------- 193
RHOD    O00212 -------------------------------------------QGFCVVT---------- 210
RHOF    Q9HBH0 ---------------------------------------------LCLLL---------- 211
RHOG    P84095 -----------------------------------------SCILL-------------- 191
RHOH    Q15669 -------------------------------------------INECKIF---------- 191
RHOJ    Q9H4E5 -----------------------------------------S---CCSII---------- 214
RHOQ    P17081 -----------------------------------------RCINCCLIT---------- 205
RHOU    Q7L0Q8 -KMKNLSKSW-------------------------------WKKYCCFV----------- 258
RHOV    Q96L33 -GVRTLSRCR-------------------------------WKKFFCFV----------- 236
RND1    Q92730 -LLHLPSRSE---------------------LI--SSTFKKEKAKSCSIM---------- 232
RND2    P52198 -SAQLSGRPD---------------------RGN-EGEIHKDRAKSCNLM---------- 227
RND3    P61587 -ISHMPSRPE---------------------LSAVATDLRKDKAKSCTVM---------- 244
RAN     P62826 EVAQTTALPDE-------------D----------------------------------- 213
RHOT1   Q8IXI2 -CIKALTR--------------IFKISDQDNDGTLNDAELNFFQRICFNTPLA------P 223
RHOT2   Q8IXI1 -CAQALTR--------------IFRLSDQDLDQALSDEELNAFQKSCFGHPLA------P 223
IFT22   Q9H7X7 ------------------------------------------------------------
RABL2A  Q9UBK7 -EEDVPDQEQS-------S--SIETPS----------------EEVASPHS--------- 228
RABL2B  Q9UNT1 -EEDVPDQEQS-------S--SIETPS----------------EEAASPHS--------- 228
RABL3   Q5HYI8 ------------------------------------------------------------
RABL6   Q3YEC7 EMMEARSRGHASPLAANGQSPSPGSQSPVVPAGAVST---GSSSPGTPQPAPQ------- 322
RRAGA   Q7L523 FLVISHYQC-K---------EQRDVHR---------------FEKISNIIKQF--KLSCS 243
RRAGB   Q5VZM2 FLVISHYQC-K---------EQRDAHR---------------FEKISNIIKQF--KLSCS 304
RRAGC   Q9HB90 KIYIAT--D-S---------SPVDMQS---------------YELCC-DMIDVVIDVSCI 298
RRAGD   Q9NQL2 KIYIAT--D-S---------TPVDMQT---------------YELCC-DMIDVVIDISCI 299
```

**Figure S2.** Sequence alignment of Ras small GTPase superfamily. Multiple sequence alignment of the G domain Ras superfamily members using Uniprot (the uniport accession codes can be taken from this figure). Based on the marked areas that are shown in Fig. S1, cysteines that are within the reach of the warhead of the eda linker are highlighted in gray. In contrast, regions that are also within reach of the bda linker are highlighted in blue. The cysteines located within these regions are marked in red. Only ~ 7% of the GTPases have cysteines within the P-loop and are therefore potential off-targets of the designed nucleotide derivatives.

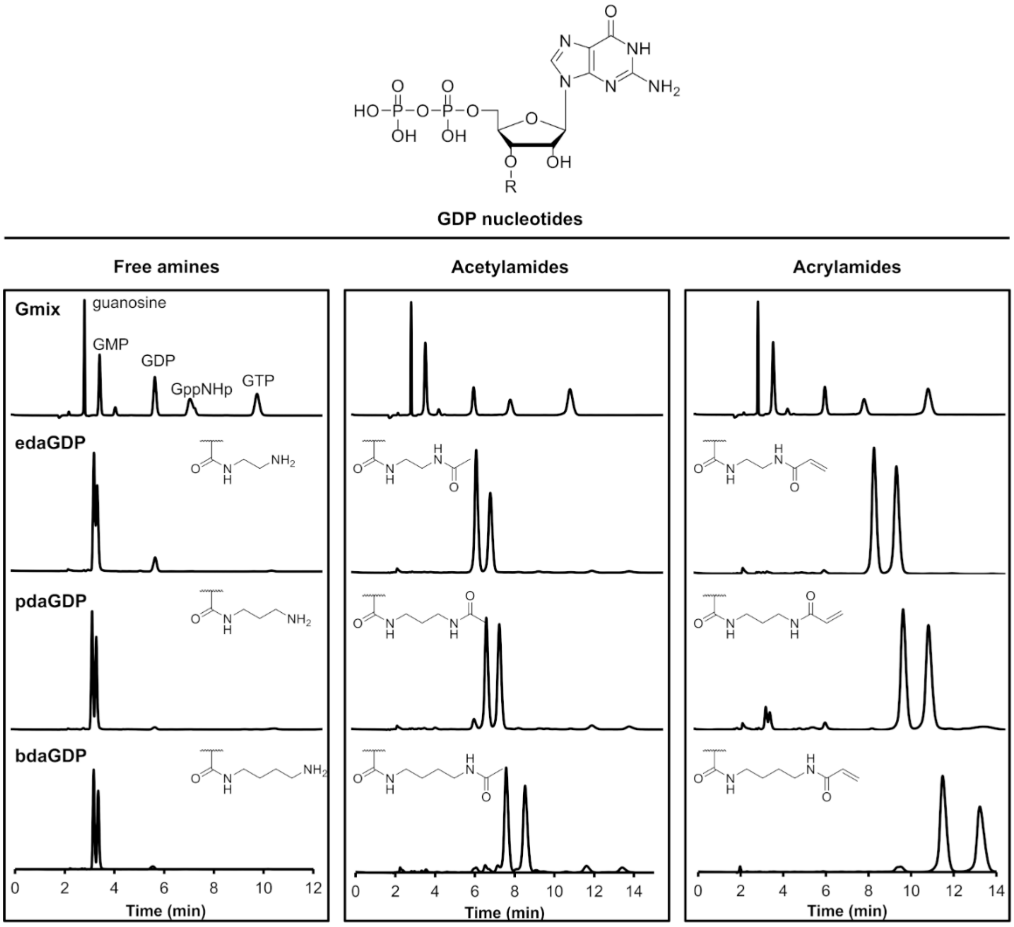


**Figure S3**. Isocratic separation of the GDP-based nucleotide analogues. Isocratic HPLC runs of (left) free amines, (middle) acetylamides and (right) acrylamides of the GDP derivatives resulting in the formation of two partially overlapping peaks corresponding to the 2'- and 3'-isomers.

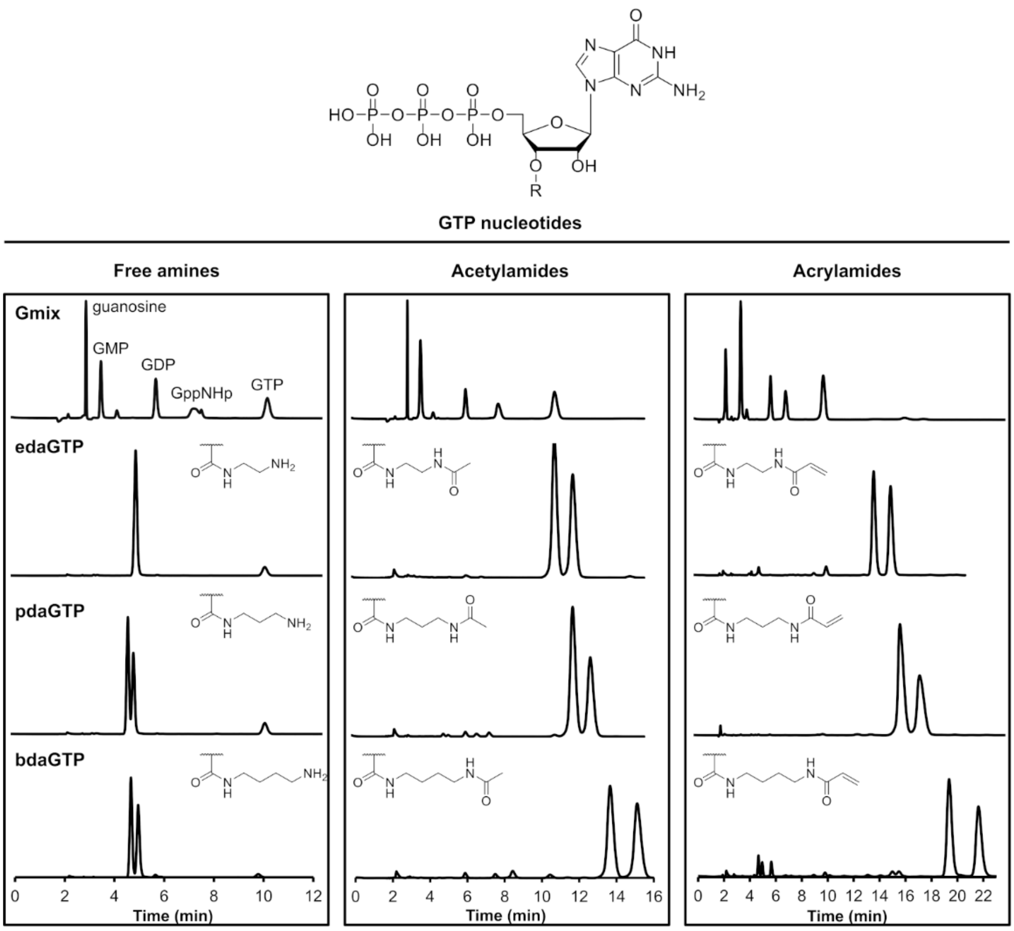


**Figure S4.** Isocratic separation of the GTP-based nucleotide analogues. Isocratic HPLC runs of (left) free amines, (middle) acetylamides and (right) acrylamides of the GTP derivatives resulting in the formation of two overlapping peaks corresponding to the 2'- and 3'-isomers.

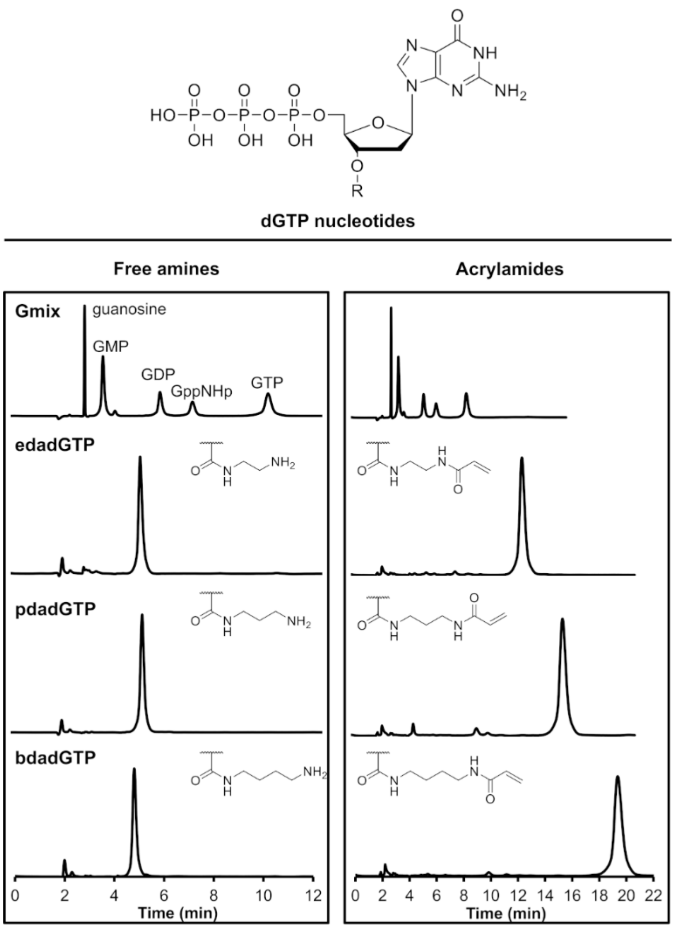


**Figure S5.** Isocratic separation of the dGTP-based nucleotide analogues. Isocratic HPLC runs of (left) free amines, (middle) acetylamides and (right) acrylamides of the dGTP derivatives resulting in the formation of one peak corresponding to the 3'-isomers.

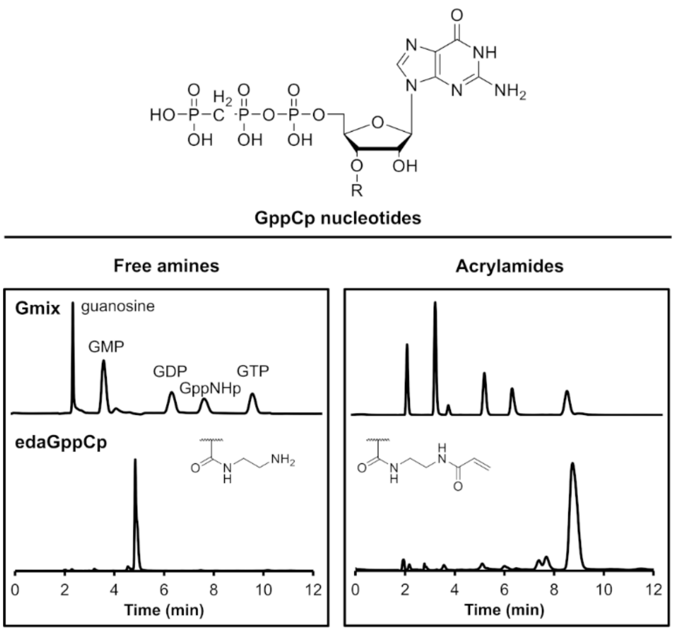


**Figure S6.** Isocratic separation of the GppCp-based nucleotide analogues. Isocratic HPLC runs of (left) the free amine and (right) the acrylamide of the edaGppCp derivatives.

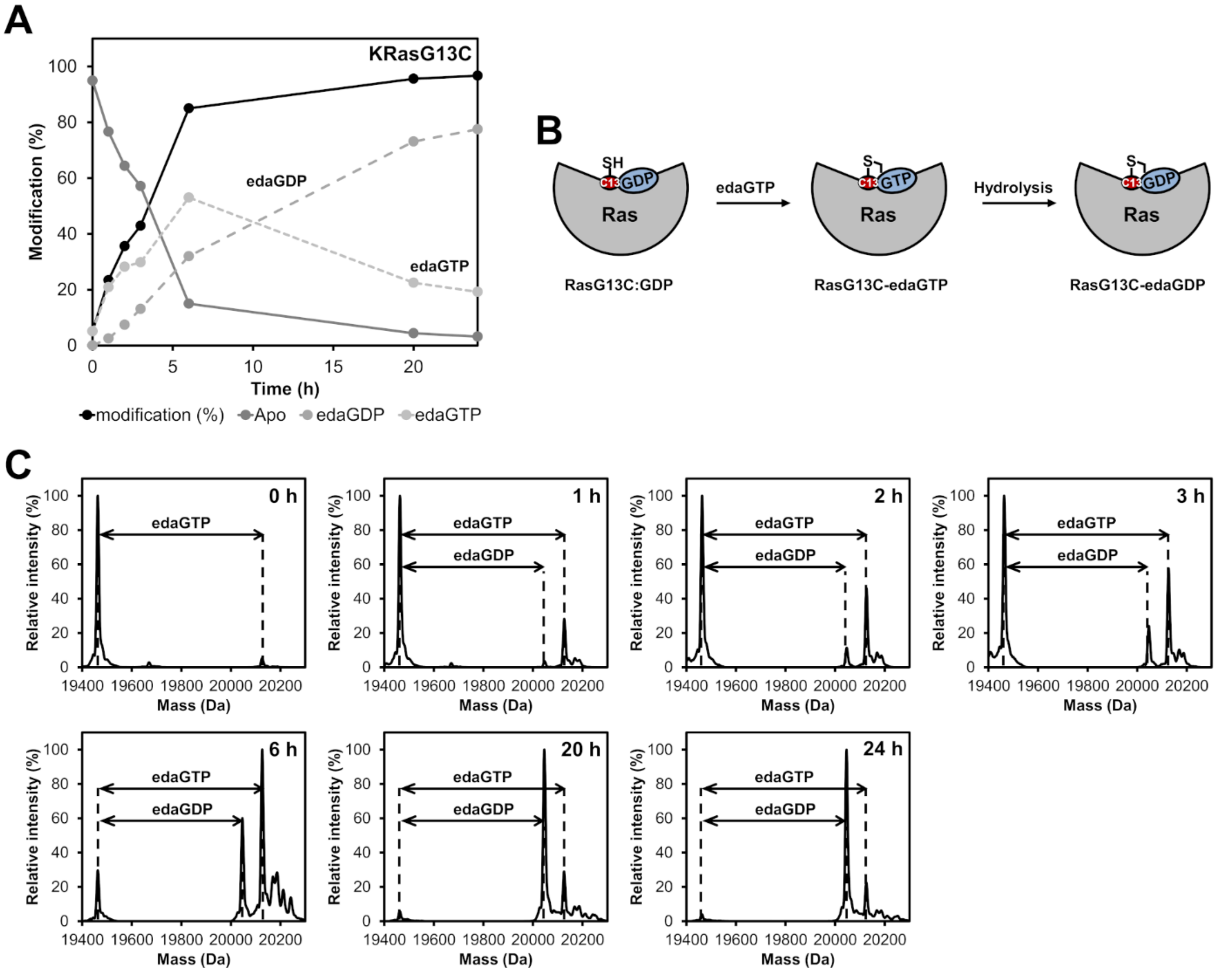


**Figure S7.** Time dependent analysis of the covalent modification of KRasG13C. (**A**) Time-resolved labelling of Ras proteins with acryl-edaGTP showing that the GTP derivative binds in the active site of Ras where it is subsequently hydrolysed. (**B**) Schematic representation of the GDP/GTP hydrolysis. (**C**) Time dependent analysis of the covalent modification of KRasG13C proteins with edaGTP at pH 9.5 at room temperature.

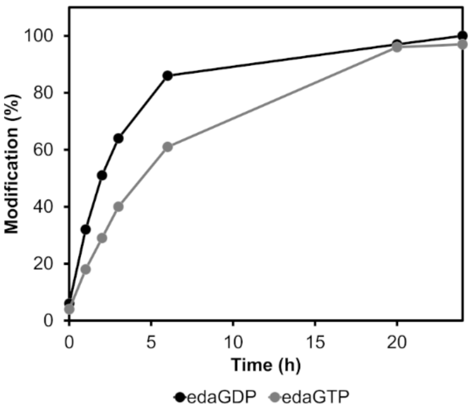


**Figure S8.** Time dependent analysis of the covalent protein modification with edaGDP or edaGTP. Time-resolved labelling of Ras proteins with acryl-edaGDP (black) or acryl-edaGTP (grey).

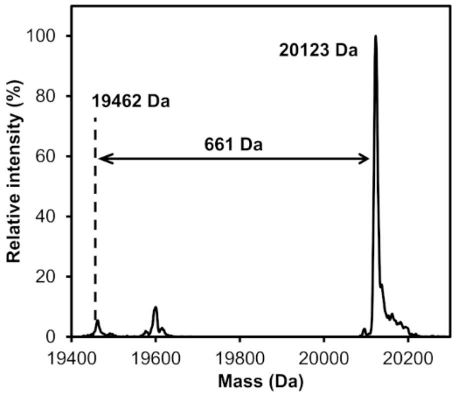


**Figure S9.** Covalent protein modification with edaGppCp. Labelling of KRasG13C$_{1\text{-}169}$ (Cys-light) with acryl-edaGppCp at pH 9.5 and room temperature was controlled after 24 h by ESI-MS.

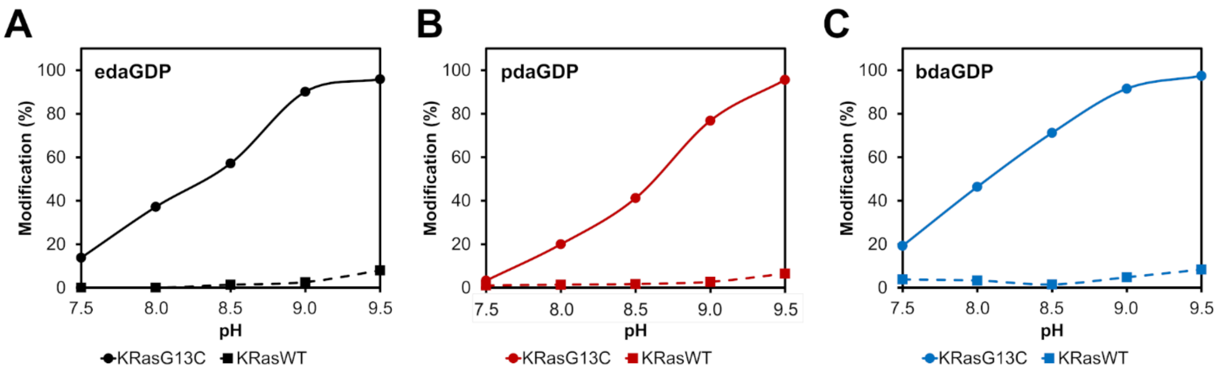


**Figure S10.** pH dependent analysis of the covalent protein modification of KRasG13C and KRasWT. (**A, B, C**) pH dependent labelling of Ras proteins with acryl-edaGDP (black), acryl-pdaGDP (red) and acryl-bdaGDP (blue) after 24 h. To increase the amount of covalently modified protein, the pH value was gradually increased. For KRasG13C$_{1\text{-}169}$ (Cys-light), a slight labelling could already be observed at pH 7.5, which, however, could be increased to almost 100% by increasing the pH to 9.5. For KRasWT$_{1\text{-}169}$ very little unspecific labelling was observed compared to the G13C mutant.

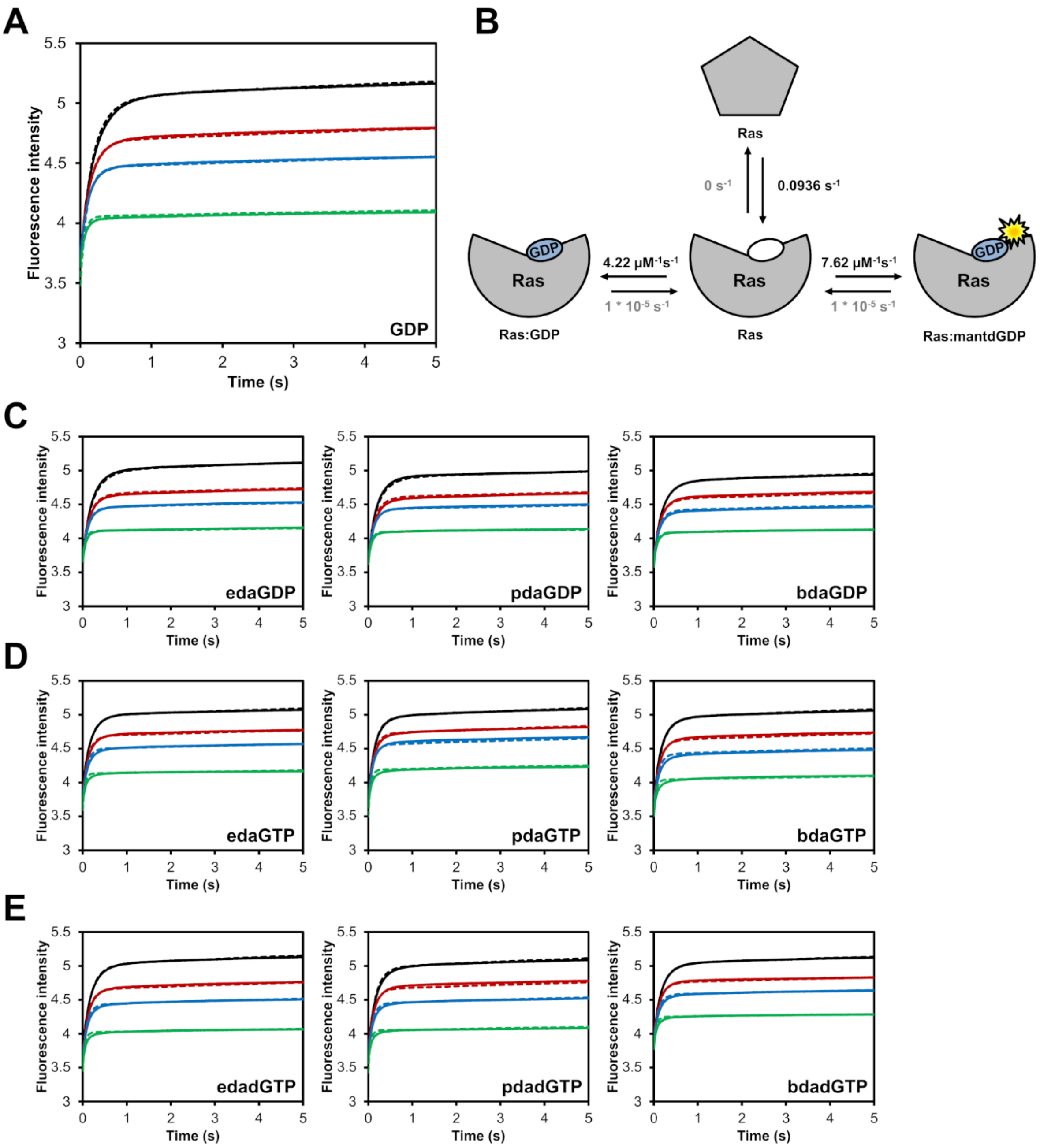


**Figure S11.** Kinetics of the nucleotide association ($k_{on}$). (**A**) Competitive binding experiments of GDP and mantdGDP; 1 µM KRas and 2 µM mantdGDP was used in the absence (black curve) or in the presence of 1 µM (red), 2 µM (blue) or 6 µM (green) competing nucleotides in a stopped-flow instrument. (**B**) The binding curves were globally fit to the indicated model using KinTek Explorer to obtain the corresponding association rates ($k_{on}$). (**C, D, E**) Competitive binding experiments between mantdGDP and GDP, GTP or dGTP analogues. Calculated $k_{on}$ values are shown in Tab. S2.

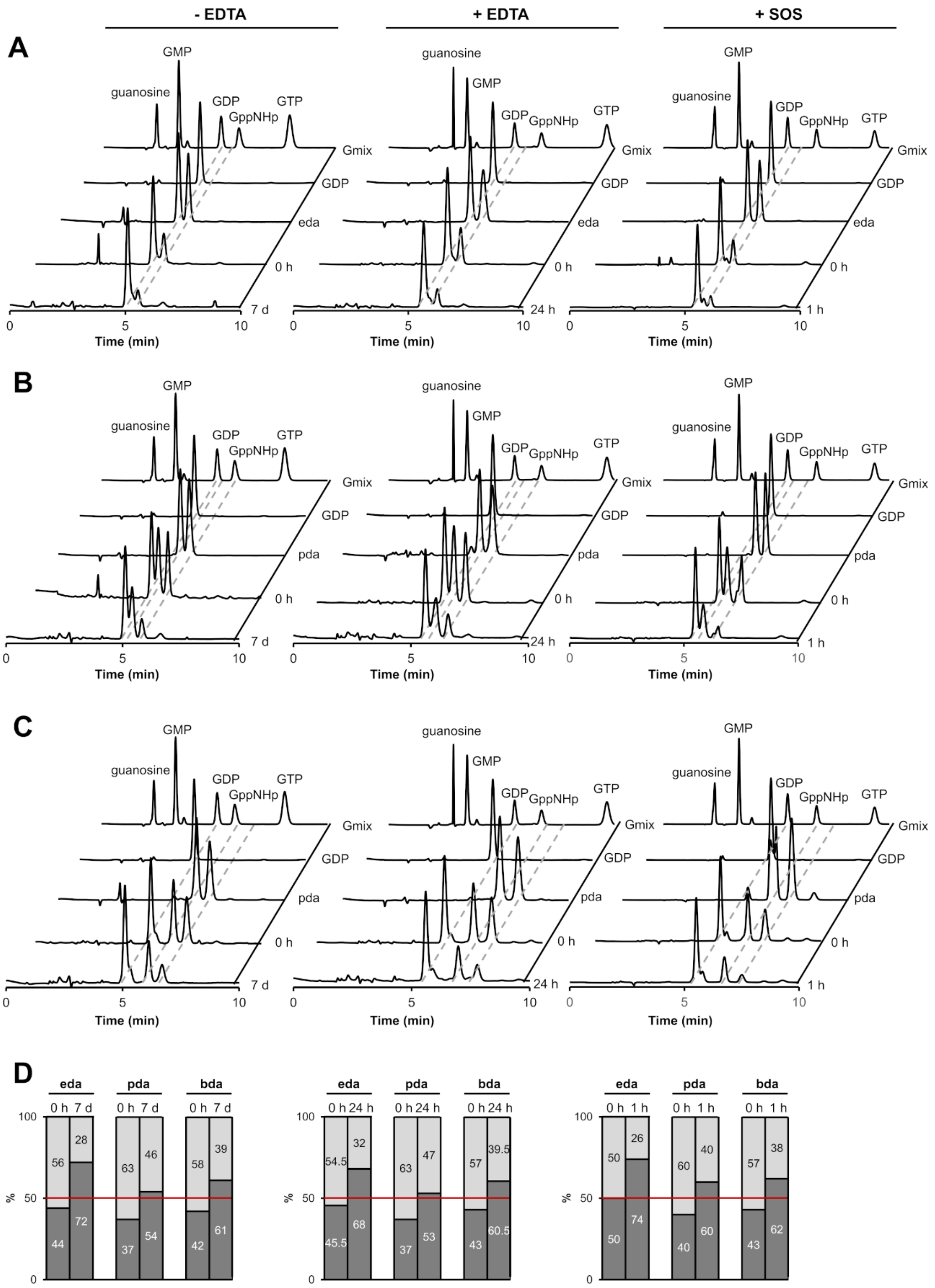


**Figure S12**. HPLC-based approach for determination of affinities relative to GDP. (**A, B, C**) Competitive binding experiments of GDP and the acetyl-derivatives of GDP; 50 µM KRasWT:GDP was mixed with 50 µM of edaGDP, pdaGDP or bdaGDP and incubated for 7 days at room temperature in the absence of EDTA, for 24 h at 4 °C in the presence of 10 mM EDTA or for 1 h in the presence of SOS (0.5 µM). After buffer exchange for removal of any unbound nucleotides, the resulting mixtures were analyzed by isocratic HPLC runs. (**D**) Relative amounts of the nucleotides and GDP bound to KRas were compared. It should be noted that the relative affinities mentioned in the results and discussion section and shown in Tab. S3 are calculated from these experiments and represent an average affinity of the 2' isomers, and that of the 3'-isomer seen to be bound in the X-ray structure is presumably actually higher than this, since it can be seen from Fig. S13 that there is a preference for one of the isomers, presumably the 3'-isomer, so that this species and the derivatives of dGDP/dGTP must have a very similar affinity to that of GDP/GTP.

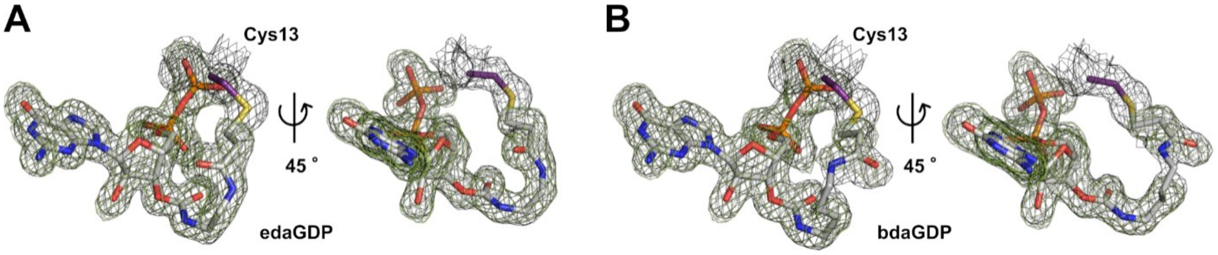


**Figure S13.** Structural insights into the binding mode of nucleotide-based covalent inhibitors. (**A, B**) edaGDP and bdaGDP, respectively, covalently bound to KRasG13C including the $F_O$-$F_C$ simulated annealing omit map (green, r.m.s.d. = 2.5) and the $2F_O$-$F_C$ map (grey, r.m.s.d. = 1.0).

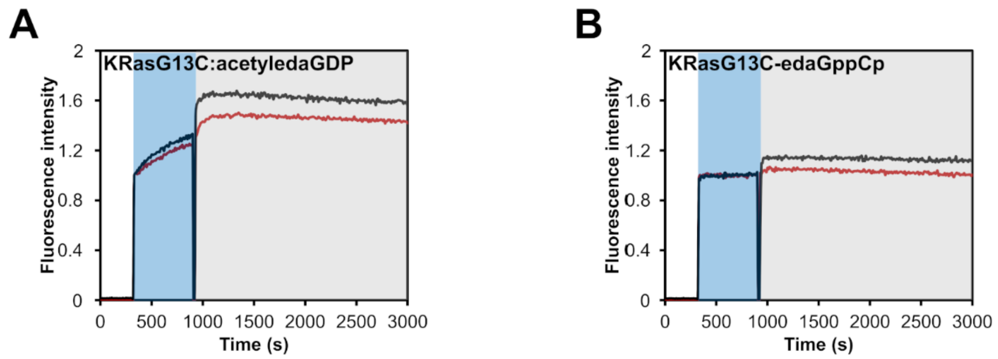


**Figure S14.** GEF-catalysed nucleotide exchange. SOS-catalysed nucleotide exchange of (**A**) KRasG13C:acetyledaGDP and (**B**) KRasG13C-edaGppCp which were mixed with an excess of mantdGDP and subsequently with 0.25 µM (red curve) or 0.5 µM (black curve) of SOS. The intrinsic nucleotide exchange is depicted in the blue box whereas the SOS-catalysed nucleotide exchange is shown in the grey box.

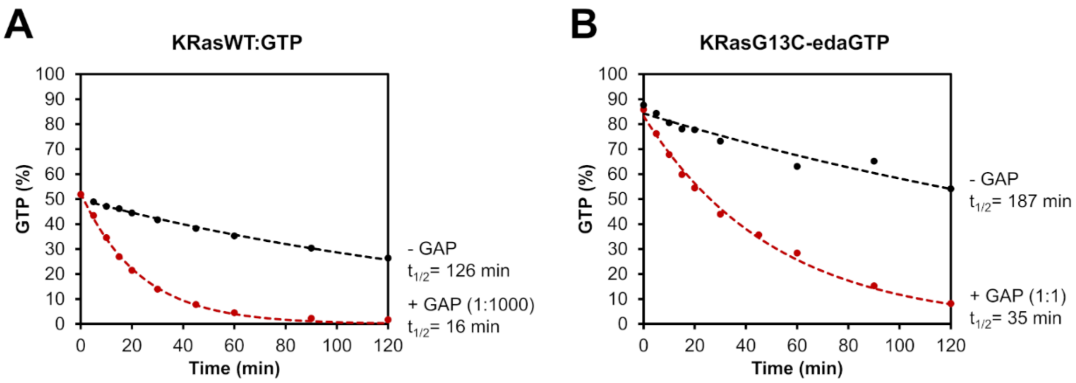


**Figure S15.** GAP-stimulated GTP hydrolysis. (**A**) The intrinsic (black) and the GAP-stimulated (red) GTP hydrolysis of KRasWT:GTP was measured via HPLC. (**B**) The intrinsic (black) and the GAP-stimulated (red) GTP hydrolysis of KRasG13C-edaGTP was obtained via ESI-MS. While GAP-stimulated GTP hydrolysis can be observed for KRasWT:GTP already at catalytic amounts of GAP (GAP 1:1000, $t_{1/2}$= 16 min), KRasG13C-edaGTP requires equivalent amounts of GAP (GAP 1:1, $t_{1/2}$= 35 min). However, the intrinsic GTP hydrolysis of KRasG13C-edaGTP ($t_{1/2}$= 187 min) is comparable to that of KRasWT:GTP ($t_{1/2}$= 126 min).

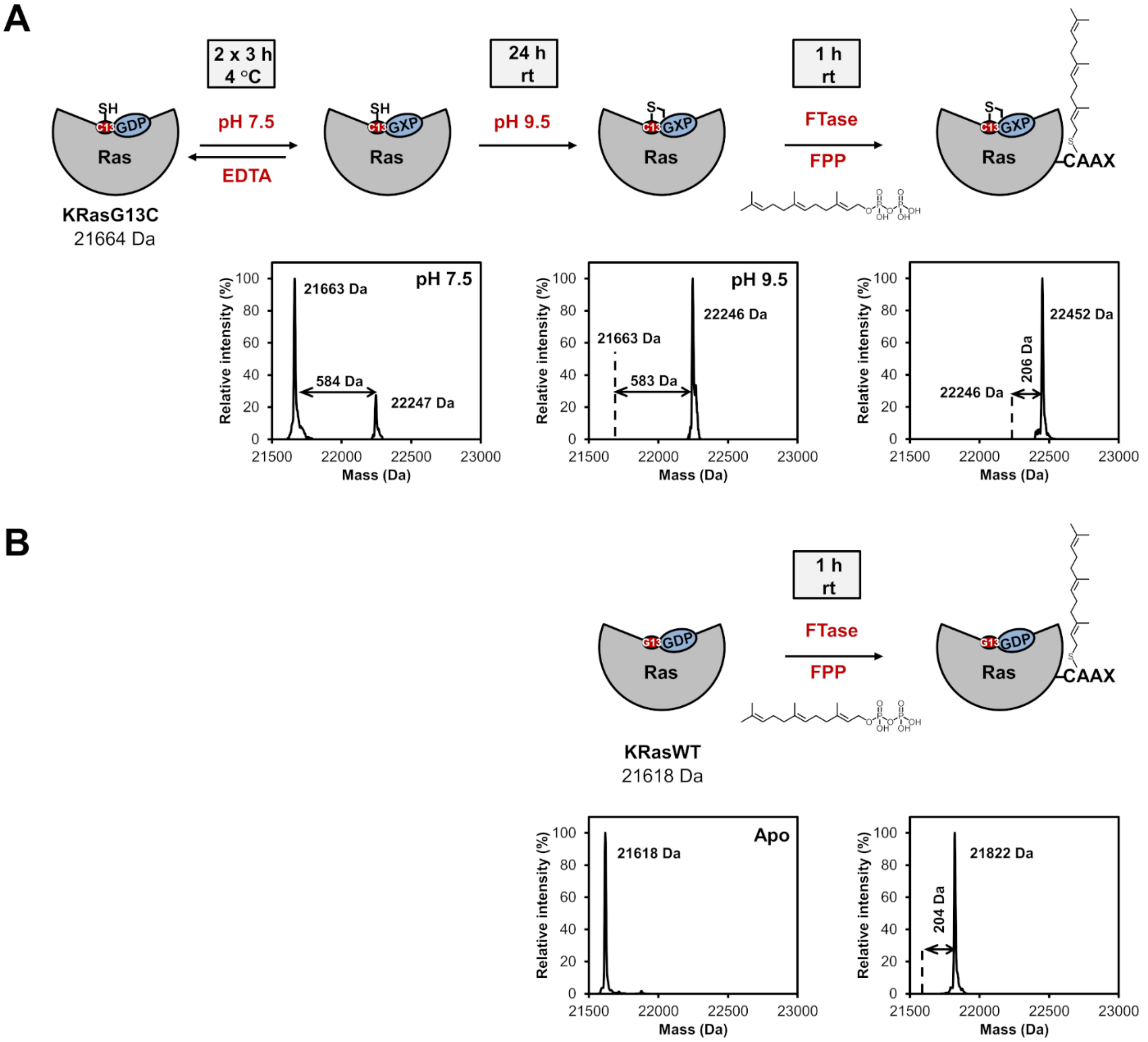


**Figure S16.** Labelling and *in vitro* farnesylation experiments with full-length KRas. (**A**) Schematic representation of covalent modification of full-length KRasG13C followed by *in vitro* farnesylation. Nucleotide exchange was performed in the presence of EDTA at pH 7.5 followed by covalent modification of full-length KRasG13C at pH 9.5 to prevent undesired interaction with the cysteine of the CAAX box. To verify the position of the covalent modification, an *in vitro* farnesylation was afterwards performed, showing complete farnesylation and thus showing that the cysteine of the important CAAX box at the C-terminus of Ras is available for farnesylation and did not react unspecifically with the nucleotide. (**B**) Schematic representation of *in vitro* farnesylation of full-length KRasWT as a control experiment.

**A**

```
                10         20         30         40         50         60         70         80
     GH MTEYKLVVVG AGCVGKSALT IQLIQNHFVD EYDPTIEDSY RKQVVIDGET CLLDILDTAG QEEYSAMRDQ YMRTGEGFLC
                90        100        110        120        130        140        150        160
KRas    VFAINNTKSF EDIHHYREQI KRVKDSEDVP MVLVGNKCDL PSRTVDTKQA QDLARSYGIP FIETSAKTRQ GVDDAFYTLV
               170        180        188
        REIRKHKEKM SKDGKKKKKK SKTKCVIM
```

**B**

| | | | KRasG13C | | | | KRasG13C-edaGDP | | | | | | |
|---|---|---|---|---|---|---|---|---|---|---|---|---|---|
| Sequence | Start position | End position | Intensity s1-a | Intensity s1-b | Intensity s1-c | Intensity s1-d | Intensity s2-a | Intensity s2-b | Intensity s2-c | Intensity s2-d | MW s1 | MW s2 | MW s1/MW s2 |
| GHMTEYK | -2 | 5 | 1.8E+09 | 2.1E+09 | 2.4E+09 | 4.1E+08 | | | 1.0E+08 | | 1.7E+09 | 1.0E+08 | 16.5 |
| GHMTEYKLVVVGAGCVGK | -2 | 16 | 2.1E+09 | 1.6E+09 | 7.1E+08 | 1.7E+09 | | 1.1E+07 | 1.6E+07 | | 1.5E+09 | 1.4E+07 | 109.4 |
| LVVVGAGCVGK | 6 | 16 | 5.6E+10 | 5.3E+10 | 4.6E+10 | 5.3E+10 | | | 3.9E+09 | | 5.2E+10 | 3.9E+09 | 13.5 |
| SALTIQLIQNHFVDEYDPTIEDSYR | 17 | 41 | 4.3E+10 | 5.0E+10 | 3.7E+10 | 4.1E+10 | 4.0E+10 | 4.1E+10 | 4.6E+10 | 2.7E+10 | 4.3E+10 | 3.8E+10 | 1.1 |
| KQVVIDGETCLLDILDTAGQEEYSAMR | 42 | 68 | 1.4E+10 | 1.6E+10 | 1.1E+10 | 1.3E+10 | 9.4E+09 | 1.1E+10 | 1.2E+10 | 5.1E+09 | 1.3E+10 | 9.4E+09 | 1.4 |
| QVVIDGETCLLDILDTAGQEEYSAMR | 43 | 68 | 2.2E+10 | 2.9E+10 | 2.3E+10 | 2.1E+10 | 1.2E+10 | 2.0E+10 | 2.2E+10 | 6.0E+09 | 2.4E+10 | 1.5E+10 | 1.6 |
| TGEGFLCVFAINNTK | 74 | 88 | 2.8E+10 | 2.8E+10 | 2.4E+10 | 2.6E+10 | 1.9E+10 | 2.4E+10 | 2.3E+10 | 1.3E+10 | 2.6E+10 | 2.0E+10 | 1.3 |
| TGEGFLCVFAINNTKSFEDIHHYR | 74 | 97 | 4.5E+08 | 6.0E+08 | 3.9E+08 | 2.3E+08 | 1.7E+08 | 3.7E+08 | 1.2E+08 | 1.1E+08 | 4.2E+08 | 1.9E+08 | 2.2 |
| SFEDIHHYR | 89 | 97 | 9.0E+10 | 5.4E+10 | 5.1E+10 | 4.7E+10 | 2.3E+10 | 4.0E+10 | 3.8E+10 | 2.5E+10 | 6.1E+10 | 3.2E+10 | 1.9 |
| VKDSEDVPMVLVGNK | 103 | 117 | 4.4E+10 | 5.9E+10 | 3.6E+10 | 9.9E+09 | 7.8E+09 | 1.7E+10 | 1.4E+10 | 9.4E+09 | 3.7E+10 | 1.2E+10 | 3.1 |
| VKDSEDVPMVLVGNKCDLPSR | 103 | 123 | 1.4E+08 | 1.2E+08 | 1.8E+08 | 2.2E+08 | 6.4E+07 | 1.3E+08 | 1.3E+08 | | 1.7E+08 | 1.1E+08 | 1.5 |
| DSEDVPMVLVGNK | 105 | 117 | 1.5E+10 | 1.8E+10 | 1.2E+10 | 1.5E+10 | 5.7E+09 | 1.5E+10 | 1.4E+10 | 4.0E+09 | 1.5E+10 | 9.6E+09 | 1.6 |
| TVDTKQAQDLAR | 124 | 135 | 2.7E+08 | 3.5E+08 | 2.4E+08 | 4.4E+08 | 9.4E+07 | 5.7E+07 | 3.2E+07 | 1.1E+07 | 3.3E+08 | 4.8E+07 | 6.7 |
| SYGIPFIETSAK | 136 | 147 | 4.5E+10 | 4.8E+10 | 3.6E+10 | 4.0E+10 | 3.2E+10 | 4.0E+10 | 4.1E+10 | 1.8E+10 | 4.2E+10 | 3.3E+10 | 1.3 |
| QGVDDAFYTLVR | 150 | 161 | 4.0E+10 | 4.6E+10 | 3.7E+10 | 4.3E+10 | 2.7E+10 | 3.6E+10 | 3.6E+10 | 1.7E+10 | 4.1E+10 | 2.9E+10 | 1.4 |

**C**

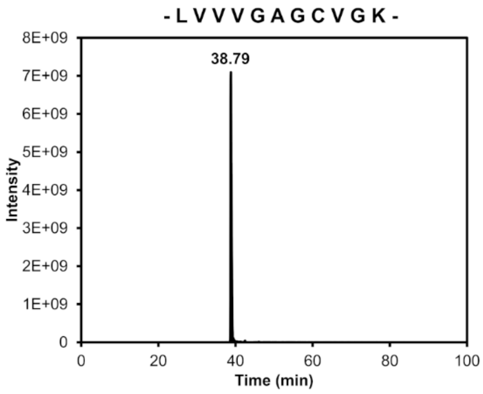


**D**

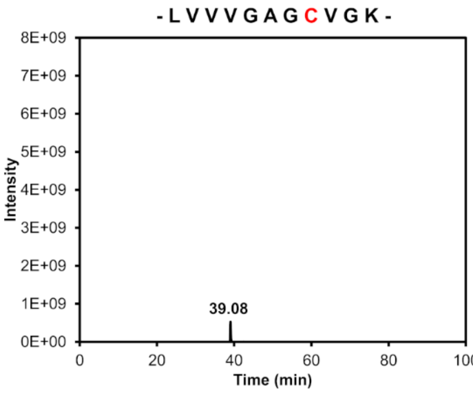


**Figure S17.** MS/MS analysis. (**A**) Overview of the sequence coverage. The identified regions of the protein are marked in green, and only the C-terminus of the protein cannot be identified due to the multiple lysine side chains present in this region. (**B**) Comparison of the intensities of all peptides of KRas after tryptic digestion, which were detected in all four replicates of the unmodified protein with an intensity of at least 1E8. Peptide intensities of the unmodified protein and the modified one are comparable from position 16 on, whereas the intensities of the peptides containing the targeted cysteine residue were significantly lower. By calculating the mean values of the intensities for the replicates of the unmodified (s1) and modified proteins (s2) followed by calculating the ratio of the mean values (MW s1/MW s2), we observed that the N terminus up to position 16 in the samples s2 could not be identified or was detected with much lower intensity than the other detectable regions of KRas. Thus, MS/MS analysis suggests that the targeted cysteine residue at position 13 is labelled by the nucleotide-based inhibitor edaGDP. (**C, D**) Comparison of the intensities of peptide LVVVGAGCVGK (m/z: 529.805+-5 ppm, 2-fold charged) for KRasG13C and KRasG13C-edaGDP. (**D**) Ion chromatogram of the peptide for KRasG13C. (**D**) Ion chromatogram of the peptide for KRasG13C-edaGDP.

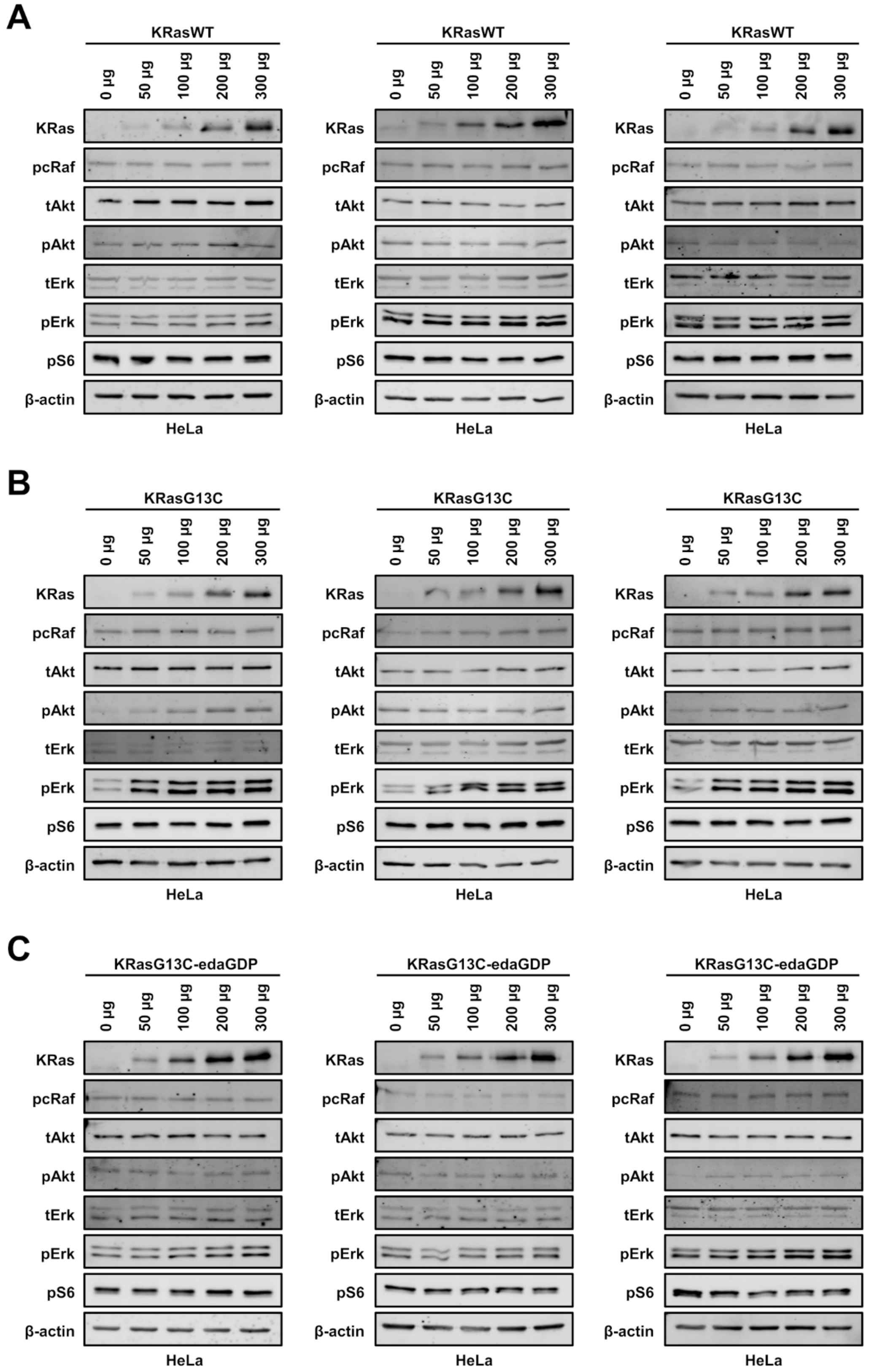


**Figure S18.** *In vitro* immunoblotting experiments. Western blot analysis after electroporation of indicated amounts of full-length (**A**) KRasWT, (**B**) KRasG13C and (**C**) KRasG13C-edaGDP into HeLa cells (n=3 biological replicates).

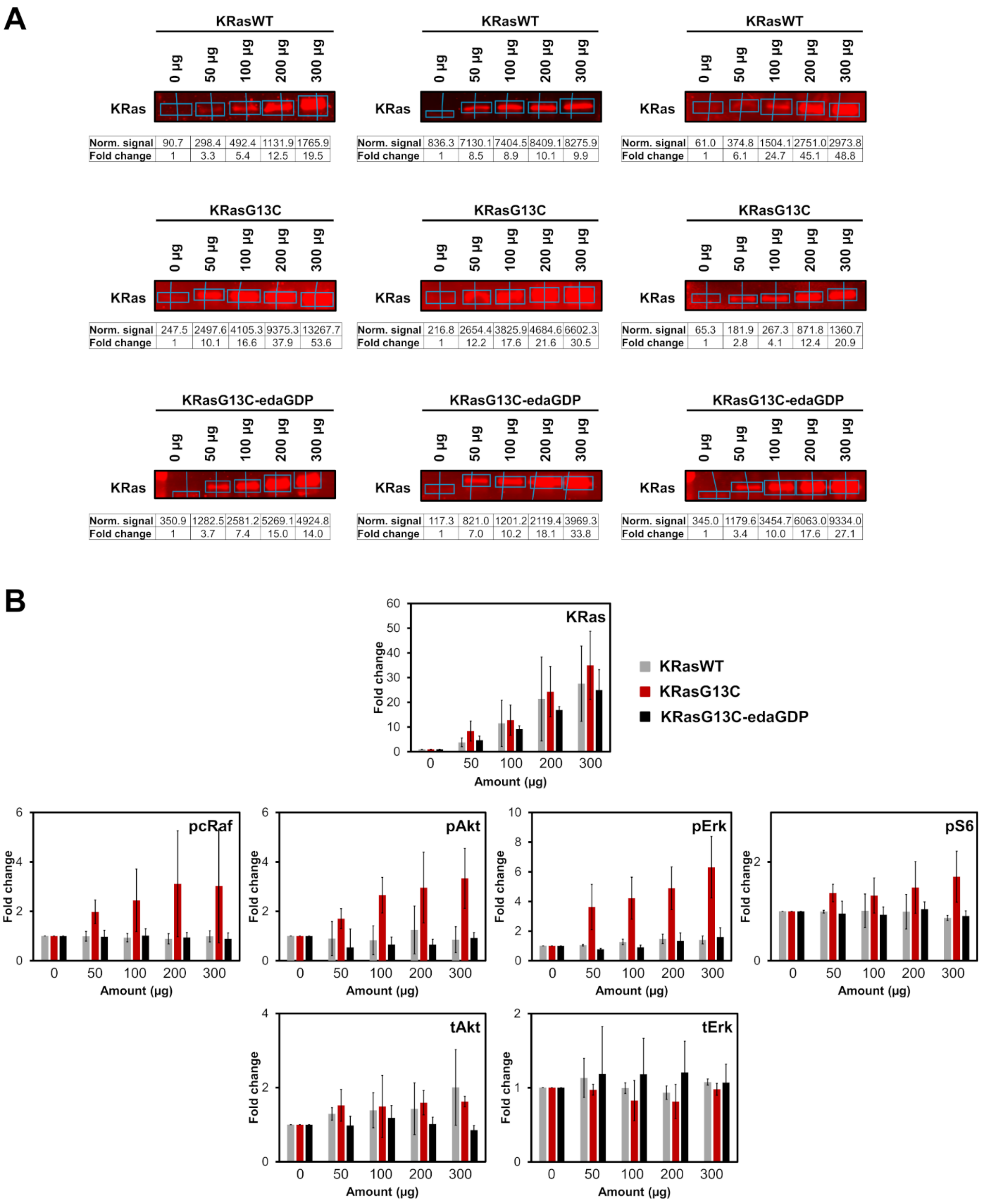


**Figure S19.** Western blot analysis and quantification. Quantification of (**A**) KRas and (**B**) downstream protein levels from western blots after electroporation of indicated amounts of full-length KRasWT, KRasG13C and KRasG13C-edaGDP into HeLa cells using Empiria Studio (Li-Cor). The mean fold change was plotted against increasing amounts of protein. Error bars indicate the standard deviation for each measurement (n = 3). Because of the relatively low expression levels of Ras, the contrast needed to be appropriately increased to enable visualisation of the intrinsically expressed KRas protein.

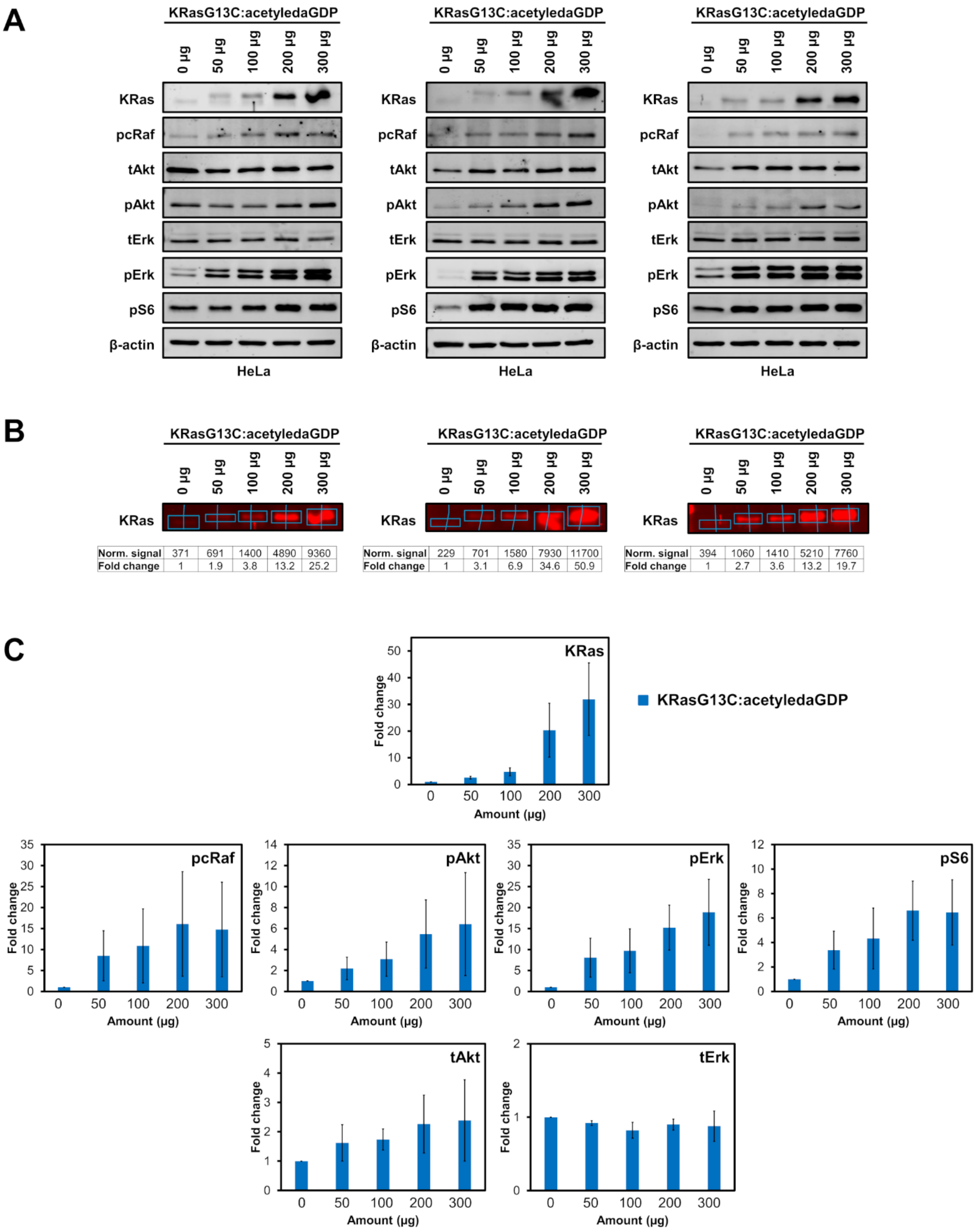


**Figure S20.** Electroporation of KRasG13C:acetyledaGDP. (**A**) Western blot analysis after electroporation of indicated amounts of full-length KRasG13C:acetyledaGDP into HeLa cells (n=3 biological replicates). (**B**) Quantification of KRas and (**C**) downstream protein levels from western blots after electroporation of indicated amounts of full-length KRasG13C:acetyledaGDP (blue) into HeLa cells using Empiria Studio (Li-Cor). The mean fold change was plotted against increasing amounts of protein. Error bars indicate the standard deviation for each measurement (n = 3).

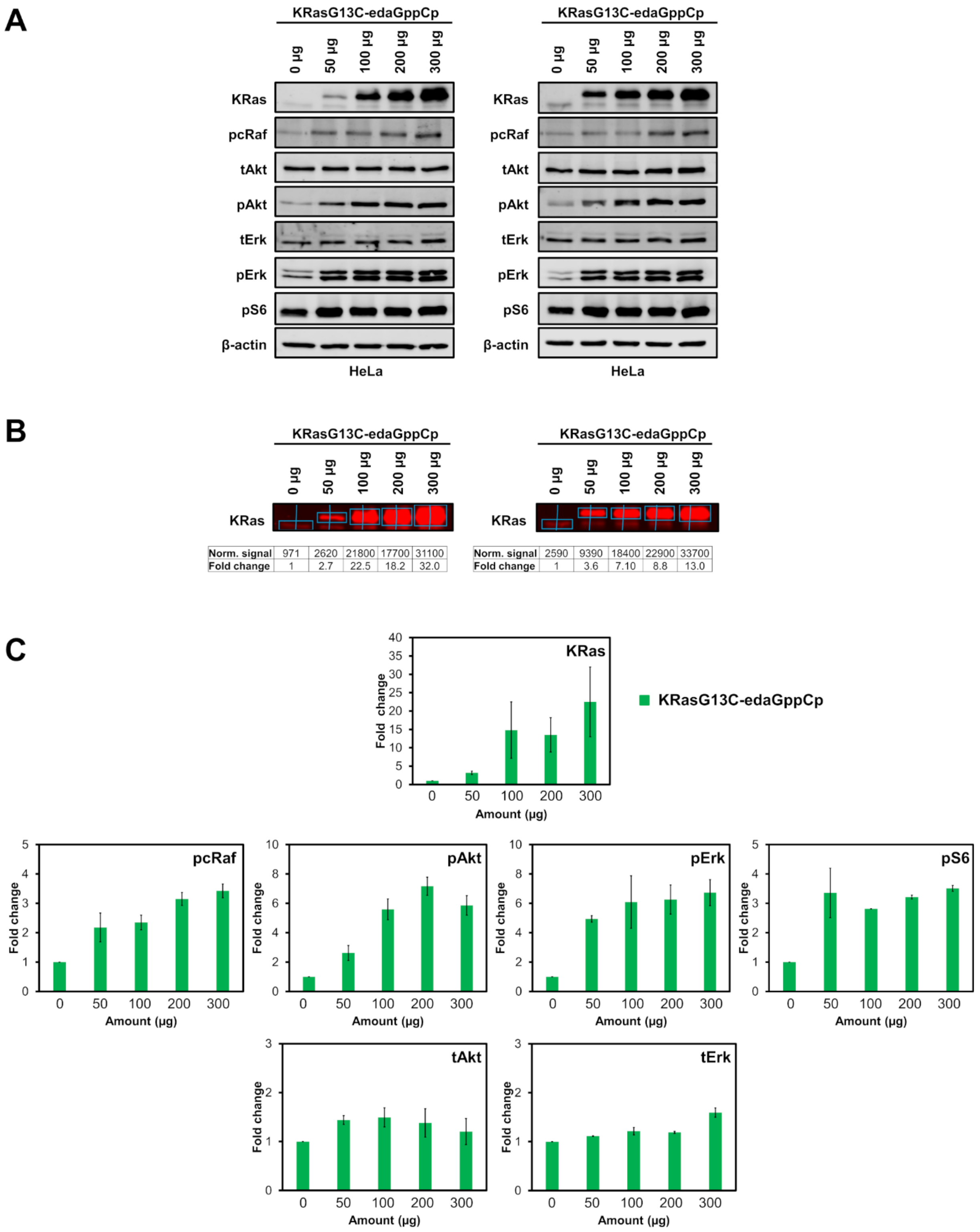


**Figure S21.** Electroporation of KRasG13C-edaGppCp. (**A**) Western blot analysis after electroporation of indicated amounts of full-length KRasG13C-edaGppCp into HeLa cells (n=2 biological replicates). (**B**) Quantification of KRas and (**C**) downstream protein levels from western blots after electroporation of indicated amounts of full-length KRasG13C-edaGppCp (green) into HeLa cells using Empiria Studio (Li-Cor). The mean fold change was plotted against increasing amounts of protein. Error bars indicate the standard deviation for each measurement (n = 2).

**Table S1:** pKa calculations. Overview of calculated pKa values of the KRasG12C and G13C mutants in the presence (+) and absence (-) of GDP. The linker in case of the G13C mutants was removed so that all featured structures had GDP and a free cysteine in its active center. The pKa calculations showed that both cysteines at position 12 and 13 have similar pKa values indicating that also position 13 should generally be addressable by covalent warheads.

| | **KRas structure** | | **pKa Cys (12/13)** | | |
|---|---|---|---|---|---|
| **KRas Mutant** | **PDB** | **Ligand** | **+** | **-** | **Δ** |
| G13C | 7ok3 | GDP (from edaGDP) | 9.85 | 8.85 | 1.0 |
| | 7ok4 | GDP (from bdaGDP) | 9.85 | 8.85 | 1.0 |
| G12C | 4ldj | GDP | 10.65 | 9.35 | 1.3 |
| | 4l8g | GDP | 10.35 | 9.05 | 1.3 |

**Table S2:** $k_{on}$ calculation. Overview of $k_{on}$ rate constants obtained from competitive binding experiments with mantdGDP (Fig. S11).

| | nucleotide | | eda | pda | bda |
|---|---|---|---|---|---|
| **$k_{on}$ [$\mu M^{-1}s^{-1}$]** | **GDP** | 4.22 | 3.73 | 3.34 | 3.12 |
| | **GTP** | 5.23 | 4.39 | 3.36 | 4.21 |
| | **dGTP** | - | 4.51 | 4.38 | 4.06 |

**Table S3:** $K_D$ calculations. Overview of the calculated $K_D$ values of pdaGDP and edaGDP obtained from an HPLC-based approach; For reference, the $K_D$ values of GDP and the nucleotide analogue SML-8-73-1 are also listed (Fig. S12).

| | **nucleotide** | **-EDTA** | **+EDTA** | **+SOS** |
|---|---|---|---|---|
| | **GDP** | | 2.5 | |
| **$K_D$** | **pdaGDP** | 8.0 | 7.4 | 10.4 |
| **[pM]** | **bdaGDP** | 10.0 | 8.9 | 10.0 |
| | **SML-8-73-1** | | ~ 140 nM | |

**Table S4:** Data collection and refinement statistics for KRasG13C-edaGDP and KRasG13C-bdaGDP.

| **Data Collection** | **KRasG13C-edaGDP (PDB 7ok3)** | **KRasG13C-bdaGDP (PDB 7ok4)** |
|---|---|---|
| Space group | P 63 | P 63 |
| Cell constants | | |
| a, b, c (Å) | 73.40, 73.40, 54.20 | 73.90, 73.90, 54.80 |
| α, β, γ (°) | 90.00, 90.00, 120.00 | 90.00, 90.00, 120.00 |
| Resolution (Å) | 41.243 – 1.6 (1.7-1.6) | 41.625 – 1.7 (1.8-1.7) |
| $R_{meas}$ (%) | 10.3 (101.6) | 7.1 (159.6) |
| $R_{merge}$ (%) | 9.8 (96.6) | 7.0 (155.7) |
| I/σ | 12.71 (2.40) | 24.52 (2.26) |
| $CC_{1/2}$ | 99.9 (80.1) | 100.0 (78.4) |
| Completeness (%) | 100.0 (100.0) | 100.0 (100.0) |
| Redundancy | 10.1 (10.4) | 20.3 (20.5) |
| **Refinement** | | |
| Resolution (Å) | 41.243 – 1.6 | 41.625 – 1.7 |
| No. Reflections | 22009 | 18838 |
| $R_{work}$ / $R_{free}$ | 15.80/20.01 (23.30/30.62) | 16.53/18.24 (26.43/35.58) |
| No. Atoms | | |
| Protein | 1386 | 1336 |
| Ligand/Ion | 38 | 40 |
| Water | 187 | 106 |
| B-factors | | |
| Protein | 27.02 | 36.92 |
| Ligand/Ion | 18.91 | 30.77 |
| Water | 36.23 | 45.00 |
| R.m.s deviations | | |
| Bond lengths (Å) | 0.016 | 0.004 |
| Bond angles (°) | 1.483 | 0.711 |
| Wavelength (Å) | 0.91504 | 0.999 |
| Temperature (K) | 100 | 100 |
| X-ray source | X10SA at SLS (Villigen, CH) | X10SA at SLS (Villigen, CH) |
| Detector | Pilatus 6M | EIGER2 X 16M |
| **Ramachandran Plot** | | |
| Outliers (%) | 0.00 | 0.00 |
| Allowed (%) | 2.38 | 2.98 |
| Favored (%) | 97.62 | 97.02 |

### Author Contributions

L.G. carried out the synthesis and performed the biochemical and kinetic experiments. T.K. synthesized the GppCp derivative and contributed to the biochemical experiments. L.G. and M.P.M. solved the crystal structures. S.M. designed and supported the electroporation experiments. L.G. performed and S.K contributed to the cellular assays. A.R. performed the in vitro farnesylation. P.J. performed MS/MS studies. H.V. and P.C. calculated the pKa values for G12C and G13C. L.G. and M.P.M. wrote the manuscript and all authors have given approval to the final version. M.P.M, R.S.G. and D.R. conceived and designed all experiments.
